# PIP5K-Ras bistability triggers plasma membrane symmetry breaking to define cellular polarity and regulate migration

**DOI:** 10.1101/2024.09.15.613115

**Authors:** Yu Deng, Tatsat Banerjee, Satomi Matsuoka, Debojyoti Biswas, Liz A. Kurtz, Jane Borleis, Yu Long, Parijat Banerjee, Huiwang Zhan, Dhiman Sankar Pal, Nathan H. Roy, Masahiro Ueda, Pablo A. Iglesias, Peter N. Devreotes

## Abstract

Symmetry breaking is a fundamental process that underlies key cellular behaviors such as cell polarity and migration, but the mechanism – how a uniform plasma membrane spontaneously transitions to an asymmetric state – is still unknown. Here in this study, using a combination of *Dictyostelium* amoeba, multiple mammalian leukocytes, and human cancer cells and 3D organoid systems, we monitored the localization dynamics of RasGTP and PIP5K, dissected the effects of eliminating, conditionally increasing, or optogenetically manipulating membrane PIP5K levels, screened for key regulators of Ras activation, and tracked single-molecules of PIP5K. Our data converge on a core biochemical circuit involving mutually inhibitory interactions between PIP5K and RasGTP that is both necessary and sufficient for symmetry breaking. The process is initiated by stochastic, localized Ras activation coupled with a decrease in the lifetime of the association of PIP5K with the membrane. A resulting localized reduction in PI(4,5)P2 facilitates the recruitment of a RasGEF to the corresponding domain, amplifying RasGTP production through a positive feedback loop. Dissociated PIP5K relocates to other membrane regions, where it suppresses Ras activation. These events separate the membrane into distinct active and inactive zones, even when receptor inputs or cytoskeletal activities are absent. This same core biochemical circuit controls the spatial organization of downstream PI3K/Akt/Rac signaling and actin/actomyosin activities, which generate the localized protrusions to define polarity and migration mode. While many models have been forwarded to explain polarity or symmetry breaking, the PIP5K-Ras mutually inhibitory bistable circuit we present here is the first universal molecular mechanism to explain the initiation of asymmetry as well as subsequent polarity and migration.

## Introduction

Cells display an extraordinary ability to spontaneously assume an endless variety of morphological forms. A common shared feature is the segregation of the membrane-cortex into two complementary states. During directed migration, for example, a cell maintains distinct front and back regions of the plasma membrane/cortex where a vast array of distinct signaling and cytoskeletal components and activities self-organize ^1^ ^2–4^. Remarkably, the configuration of the same molecular components and events that define these complementary front and back regions during migration is conserved across a variety of processes, including cortical wave propagation, macropinocytosis, phagocytosis, cytokinesis, and apical-basal polarity in epithelial cells ^5^.

The migration field has long recognized the central roles of front-back asymmetry, although conceptual paradigms often conflate the separable phenomena of symmetry breaking, front/rear polarity, motility, and directional sensing. Symmetry breaking can occur even in immobilized, round cells through segregation of otherwise homogeneous surface molecules ^6–20^. When driven by an external gradient, these localizations reflect directional sensing prior to cell shape change or movement. Motility arises as this asymmetry couples to cytoskeletal activity and front and back components segregate to or away from membrane/cortical deformations and protrusions. Polarity establishes a more stable morphological front/back axis, confines protrusions and the associated molecules largely to the front. Directed migration integrates these processes to facilitate the detection and guidance to external cues ^3,21,22^. Numerous models for directional sensing and polarity have been proposed, which typically envision some type of local activator and a more global inhibitor, which allows local responses while blocking responses elsewhere, or some form of a local pseudopod inhibition system ^23-^_36_. Most models for polarity require a functional cytoskeleton and, hence, cannot account for the initial molecular segregation step. Directional sensing models require external cues and therefore cannot explain spontaneous symmetry breaking.

A key initial symmetry-breaking event on the membrane is the spontaneous local activation of the ubiquitous oncogene Ras, as patches of activated Ras appear spontaneously or in response to environmental cues, even in immobilized cells lacking functional cytoskeletons or impaired receptor modules ^14,28,37–39^. Typically, activated Ras is found in the front of the migrating cells ^13,19,40–43^, macropinosomes ^44–46^, and in the activated, “front”-state regions of the propagating cortical waves, which are also enriched in PIP3, Akt, and F-actin ^9,17,18,47–50^. However, the mechanisms, that regulate Ras activation and amplify its stochastic fluctuations at specific membrane regions while curtailing it everywhere else, remain obscure. One clue is that PI(4,5)P2 levels are consistently depleted in membrane regions where Ras activity is high ^17,38,51–54^. Moreover, artificially lowering PI(4,5)P2 increases the size of protrusions and the frequency of spontaneous Ras patches in different cell types ^8,9,40^. Previous studies in *Dictyostelium* have shown that PLC and PI3Ks, which provide two pathways for lowering PI(4,5)P2, are not essential for migration ^55–59^, while disruption of PIP5K has been reported to impair chemotaxis^60^. Reasoning that PIP5K could be a key regulator of PI(4,5)P2 production, we wondered about the roles of this enzyme in defining localized Ras activation, symmetry-breaking and subsequent signaling and cytoskeletal pathways.

In this study, we demonstrate that a core biochemical circuit involving mutually inhibitory interactions between PIP5K and RasGTP, induces symmetry breaking, and thereby, regulates cell polarity and migration. In diverse cell types, PIP5Ks dynamically localize within back regions of the membrane, complementary to domains where the Ras/PI3K/F-actin network is activated. Consistently, knockout of PIP5K results in hyperactivation of signaling activities and a dramatic increase in migration speed. Inducible expression of PIP5K virtually eliminates Ras/PI3K/Rac1 signaling, impairs ventral wave propagation, desensitizes cells to receptor stimulation, and blocks cell migration in two and three dimensions. Using new optogenetic actuators, we showed that the rapid alterations of the level of PIP5K on the membrane can be sufficient to regulate protrusion formation and symmetry breaking. Shifting the balance of PIP5K-RasGTP circuit triggers drastic cytoskeletal remodeling in terms of branched F-actin vs linear actomyosin assembly. Remarkably, this bistable molecular circuit can regulate symmetry breaking and signaling outputs even in the cells that lack cytoskeletal dynamics or receptor modules.

Further dissecting the mechanism, we found that the lowered level of PIP5K/PI(4,5)P2 facilitates the recruitment of a specific RasGEF there, and thereby, amplifies Ras activation. On the other hand, our single-molecule measurements demonstrated that the GTP-bound Ras decreases the lifetime of PIP5K molecules on the membrane. The dissociated PIP5K molecules relocate to other membrane domains and curtail Ras activation there, closing a mutual inhibition driven positive feedback loop. A computational framework, that implements this self-amplification circuit inside an excitable network topology, combined with a viscoelastic model of morphological changes, correctly predicts the PIP5K tuning driven modulation of signaling activation, wave propagation, and cell migration. Together, our study demonstrates that the mutually exclusive localization and interaction of PIP5K and activated Ras forms a core signaling network that is necessary and sufficient to regulate symmetry breaking and generate polarity for cell migration.

## Results

### Loss of PIP5K alters migration and activates the Ras/PI3K network

It is well-documented that PI(4,5)P2 plays a central role in numerous cellular physiological processes ^61–64^. The essential question, however, is whether, under normal physiological circumstances, PI(4,5)P2 is a positive or negative regulator of signal transduction and cytoskeletal activities. The inconsistency of a report that PIP5K had little effect on random cell migration ^60^ with our previous reports of strong migration phenotypes caused by synthetically lowering PI(4,5)P2 ^8^, prompted us to reexamine this issue. A careful observation of *pikI-* cells, which lack the single PIP5K gene in *Dictyostelium* ^60^, revealed a heterogeneous spectrum of migratory patterns (Figure 1A, Video S1). To quantify these migratory modes and morphological features, we developed an automated hierarchical cell classification algorithm (Figure S1A-B, S2A-G, see methods). While a portion of cells retained wild-type amoeboid behavior, a significant fraction exhibited fan-shaped, keratocyte-like movements or oscillatory spreading/contracting movements (Figure S3A), consistent with the previously reported phenotype induced by abrupt synthetic reduction of PI(4,5)P2 in wild-type cells ^8^. These observations were supported by color-coded temporal overlay profiles (Figure S3B). Tracks of migrating cells showed that fan-shaped and oscillatory cells migrate more rapidly than WT (Figure 1B, S3C). Quantification of the basal cell area of the entire population (Figure S3D) revealed an overall 4-fold increase, primarily due to flattening. Interestingly, when *pikI-* cells are grown in shaking culture, most of them become multinucleated (Figure S3E-F), presumably due to cytokinesis defects. However, over 70% of the fan-shaped cells were still mononucleated when cultured adherently (Figure S3G-H). These defects in migration probably explain the inability of *pikI-* cells to carry out chemotaxis, aggregate, and form multicellular structures in a timely fashion (Figure S3I). In addition to PIP5K, PI(4,5)P2 levels can be altered by PI3K, PTEN, and PLC (Figure 1C). While deleting PI3Ks and PLC alters migration, the effects are minor compared to those caused by the loss of PIP5K (Figure S4A-E). Cells lacking PTEN become adherent and “freeze” in place ^65,66^, unlike the extremely active *pikI-* cells.

**Figure 1.**
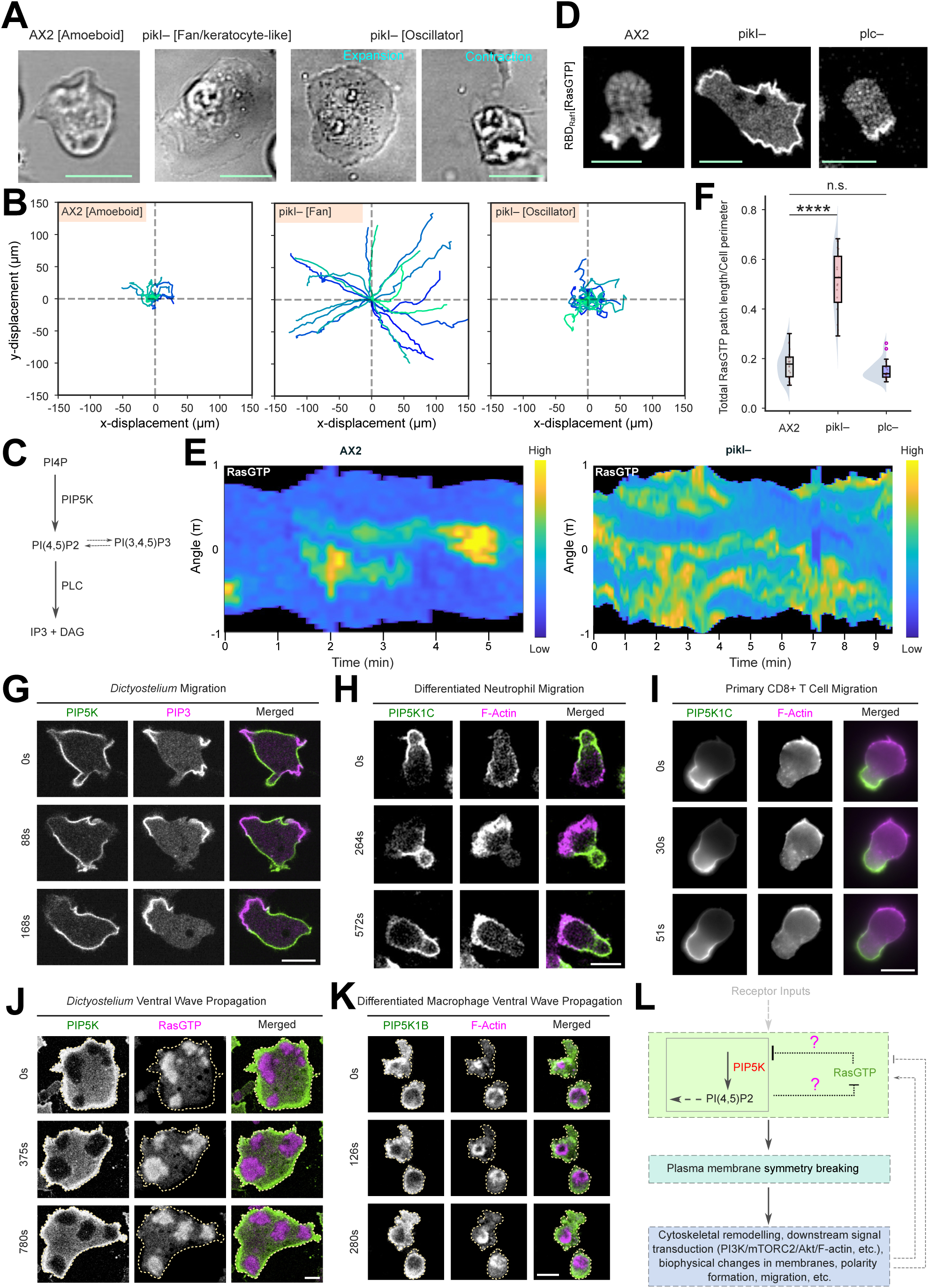
PIP5K knockout results in drastic increases cell migration speed due to hyperactivation of Ras/PI3K/F-actin network, whereas the enzyme itself dynamically localizes into the back-state regions of the plasma membrane in different cell types. **(A)** Representative live-cell DIC images of *Dictyostelium* cells where wild-type AX2 cell exhibits amoeboid morphology and *pikI– Dictyostelium cells exhibit* either fan/keratocyte-like or oscillatory (sequential coordinated expansions and contractions) morphology. For all main and supplementary figures, all scale bars represent 10 μm, unless otherwise mentioned. **(B)** Centroid tracks of randomly migrating AX2 and *pikI– Dictyostelium* cells where AX2 cells migrated predominantly via amoeboid mode whereas *pikI–* cells migrated much faster as they switched to fan/keratocyte-like or oscillatory mode of migration. Here n_c_=20 cells are shown for each migration mode and each cell was tracked for 8 minutes and was reset to the same origin. **(C)** Schematic showing major pathways for synthesizing and degrading PI(4,5)P2 on the plasma membrane. **(D)** Representative live-cell images of wild-typ*e* AX2 cells (left), *pikI–* cells (middle), and *plc-* cells (right) expressing RBD-GFP, demonstrating that *pikI–* cells have elevated Ras activation compared to AX2 and *plc-* cells. Throughout this study, Ras-Binding Domain (RBD) of Raf1 has been as a biosensor of Ras activation. **(E)** Representative 360° membrane kymograph (see “Methods” for details) of RBD-GFP intensity in AX2 (left) and *pikI–* cells (right), respectively, corresponding to Figure 1D. **(F)** Quantification of overall RasGTP level on the cell membrane in AX2, *pikI–*, and *plc–* cells, in terms of the total RBD patch length/cell perimeter. Here, n_c_ =20 (AX2), 21(*pikI–)*, or 24 (*plc–*) cells; here and everywhere for similar quantifications, each of the n_c_ cells were tracked for at least n_f_=5 frames and the average values were obtained. Throughout this study, boxes, whiskers, and outliers are graphed as per Tukey’s convention and the kernel density plots are shown on the left. The p values are computed by the Mann-Whitney U test. **(G)** Representative live-cell images of *Dictyostelium* cells coexpressing PIP5K-GFP and PH_crac_-mCherry (biosensor for PIP3) during migration, showing PIP5K dynamically moves away from PIP3-rich protrusions in migrating cells. **(H, I)** Live-cell images of migrating differentiated HL-60 neutrophils **(H)** or mouse primary CD8+ T cells **(I)** coexpressing PIP5K1C-GFP and LifeAct-iRFP703 (Lifeact is a biosensor for newly polymerized F-actin) or PIP5K1C-mNeon and LifeAct-mScarlet3, respectively **(J)** Live-cell images of *Dictyostelium* cells coexpressing mRFP-PIP5K and RBD-GFP during ventral wave propagation, showing PIP5K dynamically localizes to the back-state regions in ventral waves, maintaining consistent complementarity with respect to RBD. **(K)** Live-cell images of a HL-60 cells (differentiated into macrophages) coexpressing PIP5K1B-GFP and LifeAct-iRFP703, demonstrating that PIP5K dynamically localizes away from F-actin rich or F-actin enclosed front-state regions during frustrated phagocytosis-driven ventral wave propagation. **(L)** Schematic showing a potential core signaling network involving mutually antagonistic interaction between PIP5K and activated Ras. This core signal transduction module, either under stochastic noise or upon receiving upstream receptor input, can initiate plasma membrane symmetry breaking and thereby tune downstream signaling and cytoskeletal cascades to regulate cell polarity and migration. This module can also presumably receive positive and negative feedback from these downstream events.

To further understand the basis of *pikI-* cell phenotypes, we examined a series of signal transduction and cytoskeletal activities using specific biosensors. As expected, PH-PLCδ was mainly found on the membrane of WT cells, suggesting a significant level of PI(4,5)P2, but there was no apparent membrane association of the biosensor in *pikI-* cells (Figure S5A-B, Video S2). Next, we examined the spatial distribution of “front”- state markers starting with Ras activity in WT, *pikI-*, and *plc-* cells. Compared with WT and *plc-* cells, which have characteristic patches of Ras binding domain, RBD, at the cell protrusions (Video S4), *pikI-* cells have much broader regions (Figure 1D-F, Figure S4C, Video S3). LimE, a biosensor reflecting newly formed F-actin, also displayed much broader patches in *pikI-* cells, compared with WT and *plc-* cells (Figure S4C-D, S5C-E, Video S3). Similar results were observed with a biosensor for PIP3 (Figure S5F-G, Video S3). When oscillating cells spread, their activities increase dramatically and disappear upon cell shrinking (Figure S5H).

Next, we examined signal transduction and cytoskeletal activities typically associated with the “back”-state of cells. PH_cynA_ is a biosensor that reports the level of PI(3,4)P2 and has previously been shown to dissociate from regions of high Ras activity _67,68_. Strikingly, there generally was no detectable PI(3,4)P2 level in *pikI-* cells (Figure S5I, Video S2). This reduction is indirect, likely caused by the elevated Ras activity, since PI(3,4)P2 is not a product of PIP5K. Similar results were observed when we examined Myosin II, typically found at the cell back (Figure S5J, Video S2). Thus, our data collectively showed that loss of PIP5K causes a dramatic activation of the Ras/PI3K/Akt/F-actin network, suggesting that PIP5K is a general negative regulator.

### PIP5Ks localize to back-state regions of the membrane

Since PIP5K deletion activates cells, we wondered how PIP5K localizes with respect to activated regions in wild-type cells. We examined protrusion formation and ventral wave propagation in a wide array of cells. First, we co-expressed PIP5K with a typical front-state marker PIP3 in *Dictyostelium* cells. PIP5K was depleted from and localized complementary to PIP3 (Figure 1G, Video S5). This back localization was conserved for PIP5K1B and PIP5K1C in HL60 neutrophils during random migration (Figure S6A, 1H, Video S5), consistent with previous reports about PIP5K1B and PIP5K1C localizing to the uropod during chemotaxis ^53,54^. Consistently, PIP5K1C localized towards the back regions in mouse CD8+ primary T cells, opposite to F-actin (Figure 1I, S6B, Video S6). In addition, in MDA-MB-231 cancer cells, PIP5K1B and PIP3 exhibited complementary localizations (Figure S6C, Video S6).

We then examined the spatiotemporal profile of PIP5K during spontaneous wave propagation in electrofused *Dictyostelium* cells ^47^ and frustrated phagocytosis in HL-60 macrophage-like cells ^52^. In these seemingly diverse scenarios, PIP5K was depleted in front-state regions marked by high PI3K, Ras, or F-actin activity (Figure 1J-K, S6D-F, Video S7). As expected, PI(4,5)P2 was also depleted from front/activated-state regions of the ventral waves in *Dictyostelium* (Figure S6G, Video S7). Collectively, these results demonstrate that, during protrusion formation or wave propagation, PIP5Ks consistently localize into back-state regions, in an evolutionarily conserved fashion (Figure S6H).

To seek regions of PIP5K that control membrane binding and trailing edge accumulation, we generated a series of truncation constructs and examined their localization in *Dictyostelium* cells (Figure S7A, Video S7). Despite its robust membrane association, PIP5K does not contain a recognizable membrane association or transmembrane domain. PIP5K_301-718aa_ was still largely localized at the plasma membrane, as did full-length PIP5K, and some cells showed a relatively tighter localization to the back-state (Figure S7B, Video S8). N-terminal 315aa was not on the membrane, while PIP5K_316-718aa_ appeared significantly in the cytosol, and the protein remaining on the membrane was not localized at the back (Figure S7C-D, Video S8). Upon synthetic recruitment to the membrane, neither fragment relocalized to the back (Figure S7C-D). These results indicate that a region between 301-316 aa is important for localization to the membrane at the back. The membrane and back localization signals were not separable; the back signal may require a coincidental association of two independent sequences within the protein.

Taken together, our results indicate that PIP5K localization is the major regulator of local PI(4,5)P2 levels on the membrane, further suggesting that PI(4,5)P2 is a negative regulator of the Ras/PI3K/F-actin network and that a mutually inhibitory Ras- PIP5K circuit defines crucial evolutionarily conserved symmetry breaking events. We surmise that downstream signaling cascades such as PI3K/mTORC2/Akt reinforce the initial asymmetry, and as cytoskeletal activities remodel, polarity forms, and cells effectively migrate (Figure 1L).

### Overexpressing PIP5Ks suppresses cell protrusions and tunes migration

Reasoning that expression of PIP5K would negatively affect signaling activities and cell migration, we developed a conditional doxycycline-inducible overexpression system for PIP5K (Figure 2A). Without doxycycline (Dox) induction, *Dictyostelium* cells show their normal amoeboid behavior (Figure 2B, Video S9). Upon overnight DOX induction, two striking phenotypes appeared. Approximately 55% of the cells became significantly rounder, lacking protrusions (Figure 2C, S8A, Video S9), while the remaining cells had numerous tiny, broken protrusions, resembling short filopodia, along their perimeter (Figure S8B, Video S9); migration was severely impaired in both cases. These observations were supported by color-coded temporal overlay profiles (Figure 2D-E, S8C) and speed measurements from cell tracking (Figure 2F, S8D). Z-stack imaging showed that the cells are flatter (Figure S8E). They occupied more surface area and were less polarized (Figure S8F-G). When we applied a chemotactic gradient (Figure S9A), overall chemotaxis was strongly impaired in PIP5K OE cells. A fraction of cells migrated poorly towards the folic acid source (possibly the spiky cells), and the rest of the cells did not migrate at all (Figure S9B-D, Video S10). Slight increases in PIP5K activity led to counterintuitive results. Cells with low PIP5K expression after 2h DOX incubation became very elongated and migrated much faster than non-DOX treated cells (Figure 1F, S8F-H, S10A-C, Video S9). A similar increased speed was observed in cells expressing a PIP5K mutant (K681N, K682N), which has impaired kinase activity ^69^ (Figure 1F, S10D-E, Video S9). Next, we measured fluid uptake in cells with different levels of PIP5K. Consistent with the absent or tiny protrusions and impaired migration in cells treated with Dox overnight, FITC-dextran uptake measurements showed that macropinocytosis was significantly reduced (Figure S11A-C). Surprisingly, while *piki-* cells have very elevated PIP3, a marker for macropinocytosis ^70–72^, FITC-dextran uptake was also severely impaired in these cells (Figure S11C). Potentially, it is due to the abnormal size of the protrusions, which cannot close efficiently. Taken together, these results suggest that PIP5K activities tune the cell morphology, macropinocytosis, and migration in a nonlinear, context-dependent fashion.

**Figure 2.**
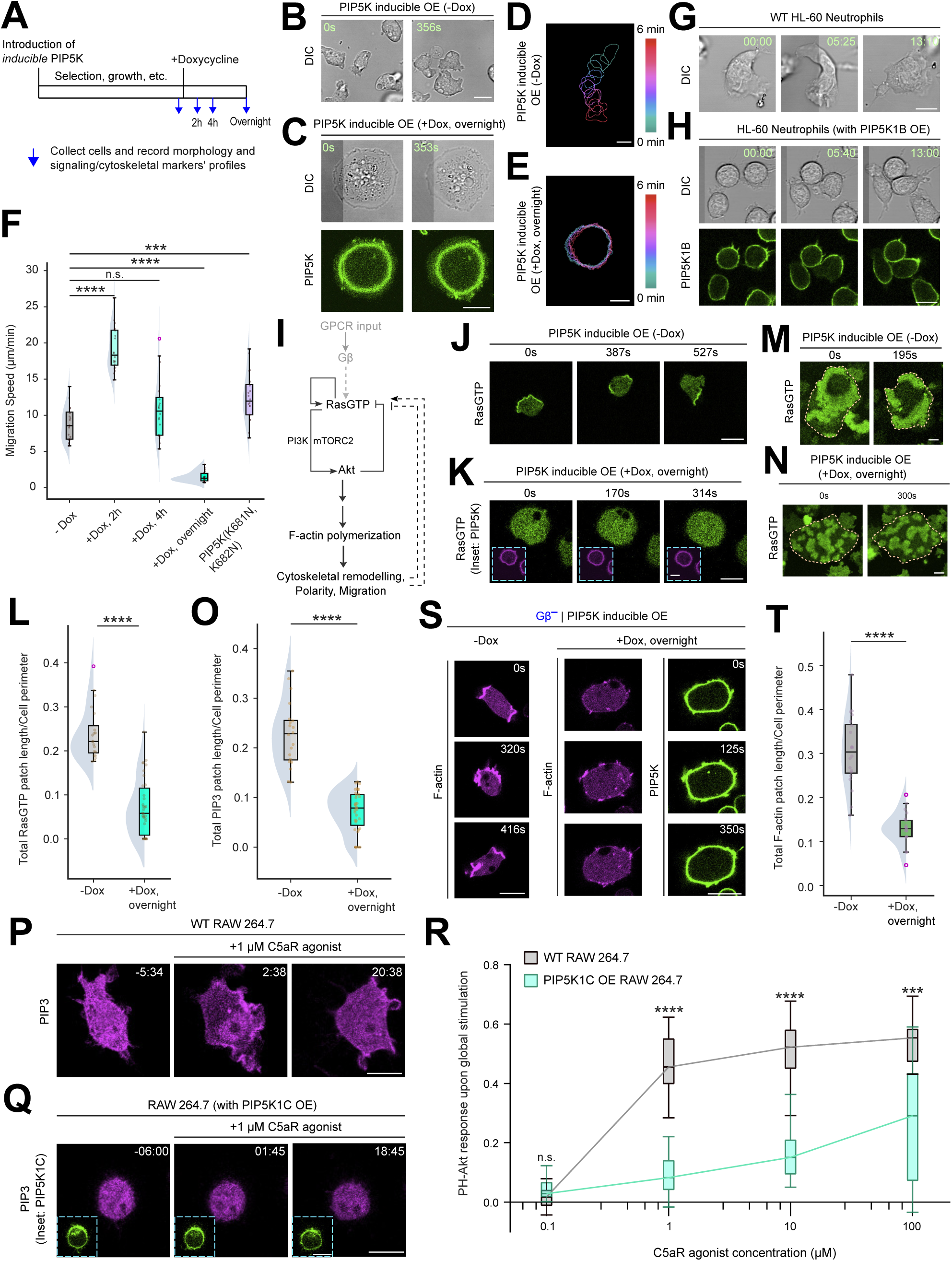
Inducible overexpression of PIP5Ks inhibits spontaneous and receptor driven signal transduction and cytoskeletal activities and severely impairs cell migration and polarity in an evolutionarily conserved fashion. **(A)** Schematic of the experimental strategy involving docycycline-inducible promoter-driven PIP5K overexpression. **(B, C)** Live-cell images of *Dictyostelium* cells expressing doxycycline (Dox)-inducible PIP5K-GFP without Dox induction (B) or with overnight Dox induction (C), showing PIP5K overexpression (OE) can make cells flatter and rounded. **(D, E)** Color-coded temporal overlay profiles of *Dictyostelium* cells expressing Dox-inducible PIP5K-GFP without Dox induction (D) or with overnight Dox induction (rounded) (E), corresponding to 2B and 2C, respectively, showing the migration of PIP5K OE cells is drastically impaired. **(F)** Quantification of the migration speed of *Dictyostelium* cells expressing Dox-inducible PIP5K-GFP upon incubation with Dox for different amount of time periods, demonstrating a nonlinear trend between the PIP5K expression levels and cell migration speeds. Here n_c_ =20 cells were tracked for each case (N≥3 independent experiments); p values by Mann-Whitney U test. **(G, H)** Shape of the migrating differentiated HL-60 neutrophils, wild-type (G) or PIP5K1B overexpressing (H), showing PIP5K1B OE cells display a rounder morphology. Time in min:sec format. **(I)** Schematic of signal transduction network that is known to control cytoskeletal structures, cell polarity, and migration. As the network can act independent of receptor inputs, receptor modules are shown in grey. Dashed black lines show putative positive and negative feedback from the downstream cytoskeletal network to the upstream signaling network. **(J, K)** Live-cell images of *Dictyostelium* cells co-expressing RBD-GFP and Dox-inducible mRFP-PIP5K without Dox induction (J) or with overnight Dox induction (K), showing RasGTP activities nearly completely vanished in rounded PIP5K OE cells (K). Inset of (K) is showing mRFP-PIP5K channel. **(L, O)** Quantification of overall RasGTP level (L) or PIP3 level (O) on the cell membrane in *Dictyostelium* cells with Dox-inducible mRFP-PIP5K overexpression syste,, without or with Dox induction, showing both RasGTP and PIP3 patches are reduced dramatically upon overnight Dox induction. The levels are quantified in terms the total RBD-GFP patch length (L) or PH_crac_-YFP patch length (O) per unit perimeter length. Here, n_c_ = 31 (L, +Dox overnight), 35 (O, +Dox overnight), or 20 (for either -Dox condition in L or O) cells. The p values are computed by the Mann-Whitney U test. **(M, N)** Ventral wave propagation in *Dictyostelium* cells co-expressing RBD-GFP and Dox-inducible mRFP-PIP5K, without Dox induction (M) or with overnight Dox induction (N), exhibiting than even when waves are present on PIP5K OE cells, they exist as many small broken waves (N), compared to their typical shape (M). **(P, Q)** Representative live-cell time-lapse images of RAW 264.7 macrophage cells, wild type (P) or overexpressing PIP5K1C-GFP (Q) where they are responding to global simulation with 1 μM of C5aR agonist. For either (P) or (Q), cells were expressing PH_Akt_-mCherry (biosensor for PIP3) whose cytosol to membrane translocation was monitored over time. Time in min:sec format and C5aR agonist was added at time t=0. **(R)** Normalized responses shown by WT (grey) and PIP5K1C-GFP overexpressing (mint green) RAW 264.7 cells to different doses of C5aR agonist, demonstrating impeded receptor activation driven signaling activation in PIP5K1C OE macrophages. Responses were computed in terms of (1 - decrease in cytosolic intensity of PH_Akt_-mCherry). Here, n_c_≥14 cells for each concentration, for either population, from N≥3 independent experiments. The p values are computed by the Mann-Whitney U test. **(S, T)** Representative live-cell time-lapse images (S) and quantification of the level of newly-polymerized F-actin (T) in Gβ– *Dictyostelium* cells co-expressing Dox-inducible PIP5K-GFP and LimE-mCherry, without Dox induction or with overnight Dox induction. Notably, Dox induction significantly decreased the F-actin polymerization level on membrane-cortex. In (T), n_c_=20 for either condition from N≥3 independent experiments; p value by the Mann-Whitney U test.

Since the localization of PIP5K was conserved from *Dictyostelium* to a series of mammalian leukocytes and cancer cells, we next tested whether the physiological roles of the enzymes were also conserved. First, we overexpressed either PIP5K1B or PIP5K1C, which are reported to be involved in chemotaxis ^53,54^, in differentiated HL-60 neutrophils. Upon overexpressing either PIP5K1B or PIP5K1C, the typical polarized neutrophils displayed either an elongated shape with PIP5Ks enriched on the tail (Figure 2G, S12A-B, Video S11) or a rounded form with PIP5Ks broadly distributed on the membrane (Figure 2H, S12C, Video S11). Cell tracking revealed that the overall average migration speed and cell areas were substantially reduced (Figure S12D-H). Overlayed cell profiles showed that the leading edge of the polarized cells was active, but the tail appeared to adhere more strongly to the surface (Figure S12I-M). Cells with rounder shapes have higher PIP5K expression levels (Figure S12N), consistent with the *Dictyostelium* results, although the round neutrophils are not as flat. Second, we overexpressed PIP5K1B and PIP5K1C in differentiated HL-60 macrophages. This led to a very flat, rounded phenotype and abolished protrusions (Figure S13A-F, Video S12). Third, we overexpressed PIP5K1C in mouse primary CD8+ T cells. Like neutrophils, low-expressing cells were polarized, with PIP5K1C at the rear (Figure 1I), while high-expressing cells were rounded (Figure S13G). The migration of expressing cells is severely impaired (Figure S13H-J).

Next, we extended our inquiry to epithelial-derived cancer cells by studying MDA-MB-231 cells. In scratch-wound closing assays (Figure S14A), cells overexpressing PIP5K1C were rounded, and the speed of closing the wound was significantly delayed (Figure S14B-C). Furthermore, overexpressing PIP5K1C dramatically prevented the dissociation of cells from 3D spheroids (Figure S14D-F). Cells were rounded but remained attached in the center of the spheroid and did not disseminate at least by 48h.

### Overexpressing PIP5Ks suppresses signal transduction and cytoskeletal activities

To dissect how PIP5K levels modulate these feedback loops, we examined the profiles of a series of signal transduction and cytoskeletal activities using biosensors. The biochemical network controlling cell migration is illustrated in Figure 2I. First, we observed the spatial distribution of Ras activity in *Dictyostelium* cells with different Dox incubation times. Without Dox incubation, cells showed characteristic RBD patches at the cell protrusions (Figure 2J, Video S13), whereas cells incubated with DOX overnight had substantially inhibited Ras activities. There were almost no RBD patches in rounded, non-moving cells (Figure 2K, Video S13), whereas in the spiky cells, there were many tiny patches (S15A-B, Video S13). These observations were supported by membrane kymographs (Figure S15C-D). Combining data from rounded and spiky cells, there was an overall 65% reduction in total Ras patch length (Figure 2L). Without Dox, cells displayed broad, sustained waves of RBD on their ventral surface; following overnight DOX, cells showed either no waves or smaller, slower-moving waves (Figure 2M-N, Video S14). Second, similar results were observed when we examined PIP3 activities in cells with different DOX incubation times, with an overall 65% reduction in cells with overnight DOX incubation (Figure 2O, S15E-H, Video S13). Third, we examined cytoskeletal activities in PIP5K-overexpressing cells. Newly polymerized F-actin exhibited analogous phenotypes in cells following overnight DOX induction, with a 60% reduction in these cells (Figure S156-E). Since rounded cells did not exhibit any membrane patches, in this and subsequent figures, we quantified spiky cells only as a conservative measure of the inhibitions. The dynamic behavior of these activities was captured in t-stack kymographs (Figure S16F), indicating that the tiny F-actin patches in cells expressing PIP5K are more transient than patches in untreated cells. Since branched actin is polymerized by Rac1/Arp2/3 activity ^20,36,46,73–75^, we used a Pak1-GBD and ArpC biosensors to examine these activities. The Rac1 activity resembled localization of F-actin (Figure S17A-C). Induced cells showed a 70% reduction in Rac1 patch size (Figure S17D-E). The ArpC patches were also much smaller in PIP5K-expressing cells (Figure S17F-G).

We have previously reported that cells lacking G-actin-sequestering actobindins A,B,C (*abnABC-*) have increased Ras and PI3K activity, suggesting that branched actin is in a positive feedback loop with these signal transduction activities ^76^. We examined the extent to which the inhibitory effect of PIP5K can interrupt this feedback loop by overexpressing the enzyme in *abnABC*-cells. Without DOX incubation, cells showed broad RBD patches as expected (Figure S17H, Video S15). Upon overnight DOX incubation, the RBD patches decreased in length and increased in number (Figure S17I, Video S15), indicating competition between the positive feedback from branched actin and inhibition by PIP5K.

We showed that, counterintuitively, cells showed a more polarized phenotype and migrated faster, compared to WT cells, when PIP5K activity was only modestly increased (Figure 2F). We examined both Ras and PIP3 signaling activities in cells incubated with DOX for only 2 hours and found that the Ras and PIP3 signaling activities were confined to tiny patches at the leading edge or undetectable (Figure S18A-B, Video S13). Similar Ras signaling activities were low or undetectable in fast-moving cells expressing mutant PIP5K (K681N, K682N) (Figure S18C, Video S16). The F-actin patches were smaller and confined to the front of these cells (Figure S18D, Video S16). We previously reported that a RasGAP C2GAPB in *Dictyostelium* cells can similarly inhibit Ras activities and increase polarity and migration speed ^39^ (Figure S19A). A possible explanation is that WT cells have slightly higher Ras activities than optimal ^45^, and reducing that activity brings it into the optimal range for polarizing cells. Since *pikI-* cells have much higher Ras activities, we tested whether attenuating Ras activity in these cells could restore polarity and migration (Figure S19A). Indeed, upon overexpressing C2GAPB, the *piki*- cells changed from fan-shaped or oscillatory cells to more polarized cells, with migration speed exceeding either *pikI-* or C2GAPB overexpressing cells (Figure S19B-D, Video S17). On the other hand, co-expressing PIP5K and C2GAPB completely shut down the migration, presumably due to severe Ras deactivation (Figure S19E-F, Video S17).

As overexpressing PIP5Ks in HL-60 neutrophils showed similar migration and morphological phenotypes to those of *Dictyostelium* cells, we tested whether effects on signaling networks are also conserved. As previously reported ^10,16,40,77,78^, PIP3 and F-actin preferentially localized at the leading edge of the migrating neutrophils (Figure S20A-B). In rounded neutrophils overexpressing PIP5K, F-actin activities were greatly diminished (Figure S20C-D). Even in the polarized populations expressing PIP5K, PIP3 activities remained low (Figure S20E-F). Thus, the inhibition of signal transduction and cytoskeletal activities by expression of PIP5Ks is a general phenomenon.

As we have shown, expressing PIP5Ks inhibits Ras and PI3K activities and leads to smaller, shorter-lived waves (Figure 2M-N). Reduced spontaneous signaling and curtailed wave activity in an excitable system indicate an increased threshold ^8^. To more directly assess the threshold, we tested how the enzyme level interferes with receptor input-driven activities. We first monitored the profile of PIP3 biosensor PH_Akt_ upon C5aR activation in RAW 264.7 macrophages ^18,79^. When cells were stimulated with 1 μM C5aR agonist, PH_Akt_ transiently translocated to the membrane and cells extended protrusions (Figure 2P, Video S18). In contrast, at this concentration, there was no PIP3 increase or protrusion formations in cells overexpressing PIP5K1C (Figure 2Q, Video S18) and there was only a moderate increase at 100 μM C5aR agonist (Figure S21A-G, Video S18). Compared with WT cells, the dose-response spanning 0.1 μM to 100 μM C5aR for PIP5KIC-expressing cells was shifted to higher concentrations (Figure 2R). Similar results were observed when we monitored Ras-GTP profiles upon folic acid (FA) stimulation in WT versus PIP5K overexpressing *Dictyostelium* cells (S22A-D). This shift in dose-response curve raised the remote possibility that seemingly spontaneous Ras/PI3K/F-actin activity is actually triggered by extraneous autocrine or paracrine signaling. To rule this out, we used *Dictyostelium* cells lacking the only Gβ subunit which make them unable to respond to chemoattractants ^11,80,81^. Nevertheless, similarly as in WT cells, overexpressing PIP5K in cells lacking Gβ decreased spontaneous F-actin activities (Figure 2S-T, Video S19). Considering that F-actin is a downstream indicator of network activity, these data indicate the inhibitory role of PIP5K is intrinsic to the system and does not act through receptor module.

### Abrupt perturbation of membrane PIP5K level can regulate symmetry breaking and protrusion formation

The dramatic phenotypes resulting from genetic manipulation of PIP5K occur on a longer timescale than changes in PIP5K distribution during cell migration and polarity. To assess the immediate effects of altering the association of PIP5K with the membrane and rule out secondary factors, such as membrane remodeling or changes in transcription profiles, we designed optogenetic systems to rapidly remove or recruit PIP5K from or to the membrane. First, we tested whether the removal of overexpressed PIP5K from the membrane can break symmetry and induce protrusion formation in PIP5K overexpression-driven highly quiescent RAW 264.7 macrophages. We developed the Opto-SeqP system to rapidly sequester PIP5K from the membrane to mitochondria (Figure 3A). Upon blue light illumination in macrophages with Opto-SeqP, membrane PIP5K levels quickly decreased, and cells began to make many small protrusions (Figure 3B-D, S23A-C, Video S20). These increased protrusion formation events often eventually resulted in an elongated cell shape (Figure 3B-C, S23A, Video S20). Quantification shows that blue light illumination significantly increased protrusion formation frequencies in cells with Opto-SeqP, whereas recruiting a control construct to mitochondria did not elicit any such changes (Figure S23D-F, Video S20).

**Figure 3.**
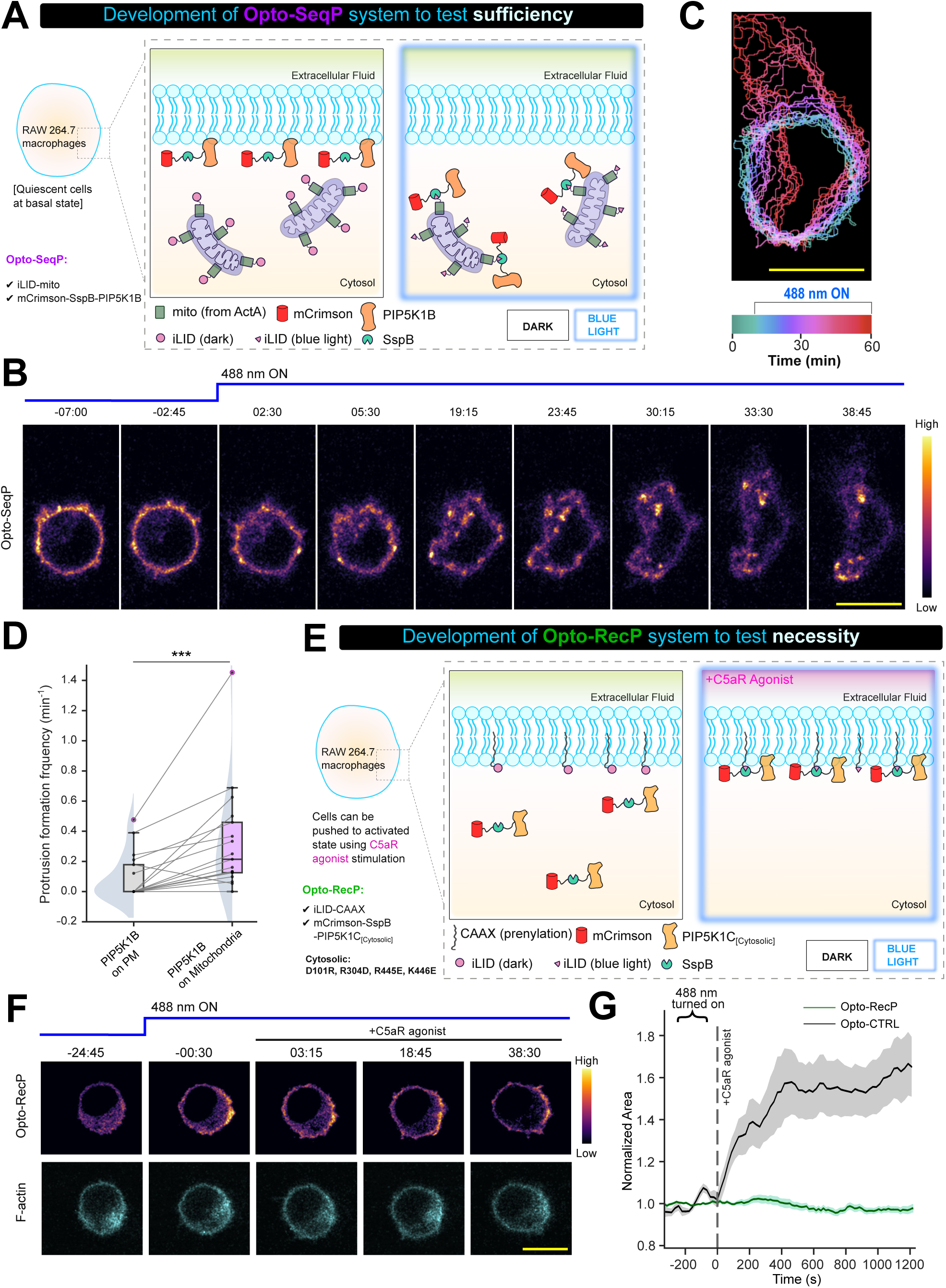
Rapid optogenetic perturbation of PIP5K level on the plasma membrane can be sufficient to regulate symmetry breaking and protrusion formation. **(A)** Schematic of the experimental setup with Opto-SeqP (mCrismon-NES-SspB-PIP5K1B-P2A-iLiD-Mito) recruitment system. To deplete PIP5K on the plasma membrane acutely, iLiD-Mito was used as the anchor so that SspB-tagged protein can be recruited and sequestrated to the outer mitochondrial membrane. This system was expressed in RAW 264.7 macrophage cells (which are largely quiescent in basal state) and optogenetic recruitment was induced using 488 nm laser illumination. **(B)** Representative time-lapse images of RAW 264.7 macrophages expressing Opto-SeqP, before and after 488 nm illumination driven recruitment of PIP5K1B from plasma membrane to mitochondrial outer membrane. 488 nm was turned ON at t=0. Note that, protrusion formation as plasma membrane PIP5K1B was lowered, cells started making protrusions and assumed am elongated, polarized shape. **(C)** Color-coded temporal overlay profiles of RAW 264.7 macrophages before and after Opto-SeqP recruitment, showing that cells spread significantly upon the recruitment of plasma membrane localized PIP5K to mitochondria. **(D)** Quantification of protrusion formation frequency before and after the recruitment of PIP5K1B from plasma membrane to mitochondria. Here, for each of the n_c_ = 18 cells, protrusion formation events were tracked, before and after recruitment. For pairwise comparison, data from the same cell are connected by grey lines. The p values are computed by the Wilcoxon matched-pairs signed-rank test. **(E)** Schematic of the experimental setup with Opto-RecP (mCrismon-NES-SspB- cytosolic PIP5K1C-P2A-iLiD-CAAX) recruitment system. To increase PIP5K on the plasma membrane acutely, iLiD-CAAX was used as the plasma membrane anchor and SspB-tagged cytosolic, functional version of of the enzyme was used as recruitee. protein to plasma membrane. The Opo-RecP system was expressed in RAW 264.7 macrophage cells, then cytosolic PIP5K (or CTRL) was optogenetically recrutied to membrane (using 488 nm laser illumination), and finally cells were globally stimulated with C5aR agonist. **(F)** Representative live-cell time-lapse images of RAW 264.7 macrophage cells co-expressing Opto-RecP system and LifeAct-iRFP703. Time in min:sec format. Note that, cytosolic PIP5K was globally recruited to membrane by turning ON 488 nm laser as indicated, and cells were then stimulated with C5aR agonist at t=0. Images demonstrate that the cell failed to generate new actin-bsed protrusions here or spread upon C5aR agonist stimulation here, even after a saturating dose of C5aR agonist stimulation. **(G)** Quantification of the cell spreading response, in terms of normalized area, upon C5aR stimulation in RAW 264.7 cells where either Opto-RecP or Opto-CTRL was recruited. The C5aR agonist was added at t=0, as indicated by dashed vertical line. Here, n_c_ = 14 cells for either case; mean±SEM are plotted.

Next, to test if rapidly increasing PIP5K level on the membrane would inhibit receptor-mediated protrusions, we developed the Opto-RecP system based on a previously reported cytosolic, functional version of PIP5K ^82^ (Figure 3E). We recruited either cytosolic PIP5K or a cytosolic control construct to the membrane of RAW 264.7 macrophages and then stimulated them with C5aR agonist (Figure 3E). Compared to the control recruitment, where C5aR stimulation induced large protrusions and strong cell spreading (Figure S24A-B, Video S21) ^17,79,83^, recruitment of the cytosolic PIP5K completely blocked the response, and cells remained quiescent (Figure 3F-G, S24C, Video S21). Furthermore, globally recruiting cytosolic PIP5K1C to the plasma membrane in MDA-MB-231 cells (pretreated with saturating doses of EGF) induced cell retraction (Figure S25A-G, Video S22), whereas recruiting the control did not induce any change in cell area or aspect ratio (Figure S26A-D, Video S22). Taken together, these observations demonstrate that the acute perturbation of the membrane PIP5K level is necessary and sufficient to regulate the plasma membrane symmetry breaking and protrusion formation events and these effects are not mediated via the long-term changes in cell physiological responses.

### Inhibiting actomyosin contractility counteracts PIP5K overexpression-induced phenotypes

Since there is a shift from front-state towards back-state activities in PIP5K overexpressing cells, we reasoned that there might be a selective inhibition of branched actin relative to linear, cortical actin. ABD120 is a protein that labels total F-actin in *Dictyostelium* cells ^84–86^, while LimE more selectively labels branched actin. We expressed both constructs in a PIP5K-overexpressing cell line and used the ABD120-LimE patch ratio to indicate the relative level of inhibition. Before inducing PIP5K, ABD120 is primarily labeled uniformly on the plasma membrane, with some enrichment at the protrusions, whereas LimE is localized almost exclusively to protrusions (Figure S27A, Video S23). Upon inducing PIP5K, cells changed to a rounder shape, and LimE patches nearly disappeared (Figure S27B, Video S23). However, ABD120 was still uniformly distributed on the plasma membrane, resulting in a higher ABD120-LimE ratio (Figure S27B-C, Video S23), suggesting a selective inhibition of branched actin. Consistent results were observed when cells were stained with phalloidin. In uninduced cells, phalloidin primarily stained the protrusions, whereas in cells expressing PIP5K, phalloidin staining colocalized with PIP5K around the perimeter (Figure S27D-E). Note that earlier (Figure 2), we found that PIP5K-expressing cells that showed the “spiky” phenotype changed to the “rounded” phenotype with CK666, further indicating that round-shaped cells have less branched actin (Figure S27F, Video S24).

In previous experiments (Figure 2), we showed that expressing PIP5Ks can eliminate signal transduction and cytoskeletal activities and cell migration. We hypothesized that under normal conditions, the inhibitory effects of PIP5Ks would be reinforced by actomyosin-based cytoskeletal activities. To test the extent to which lowering Myosin II would counteract PIP5K expression and restore migration, we co-expressed PIP5K and a recruitable myosin heavy chain kinase C (MHCKC) that was previously shown to abruptly decrease myosin II bipolar thick filament assembly in *Dictyostelium* cells ^76^(Figure S28A-B, Video S25). In PIP5K overexpressing cells, global recruitment of MHCKC to the plasma membrane caused cells to polarize and migrate (Figure S28C-F, Video S25). As shown in Figure S28G and Video S26, analogous results were observed in propagating waves of Ras activity on the ventral surface of fused giant *Dictyostelium* cells. Expressing PIP5K completely suppressed the waves, but upon adding Myosin II inhibitor, blebbistatin, the RBD waves started to burst and gradually increased until the entire surface displayed activity. These results are consistent with an independent study from our laboratory demonstrating negative feedback from myosin II to Ras ^39^.

### A bistable Ras-PIP5K interaction mediates the symmetry breaking independent of cytoskeletal dynamics

While, as we have just shown, there is an interaction between the levels of PIP5K and actomyosin, and as outlined earlier, models for cell polarity typically rely on cytoskeletal activities, symmetry breaking can occur in immobilized cells without cytoskeleton. To investigate the roles of Ras-PIP5K interaction in symmetry breaking, we applied cytoskeletal activity inhibitors to cells with different PIP5K levels. We first treated *pikI- Dictyostelium* cells with Latrunculin A to remove F-actin activities and monitored patches of RBD, which appear as dynamic crescents moving around the cell perimeter. Consistent with the large patches seen in *pikI-* cells with intact cytoskeletal dynamics, the Ras activity crescents were much wider than in WT cells and maintained typical dynamics (Figure 4A-B, Video S27).

**Figure 4.**
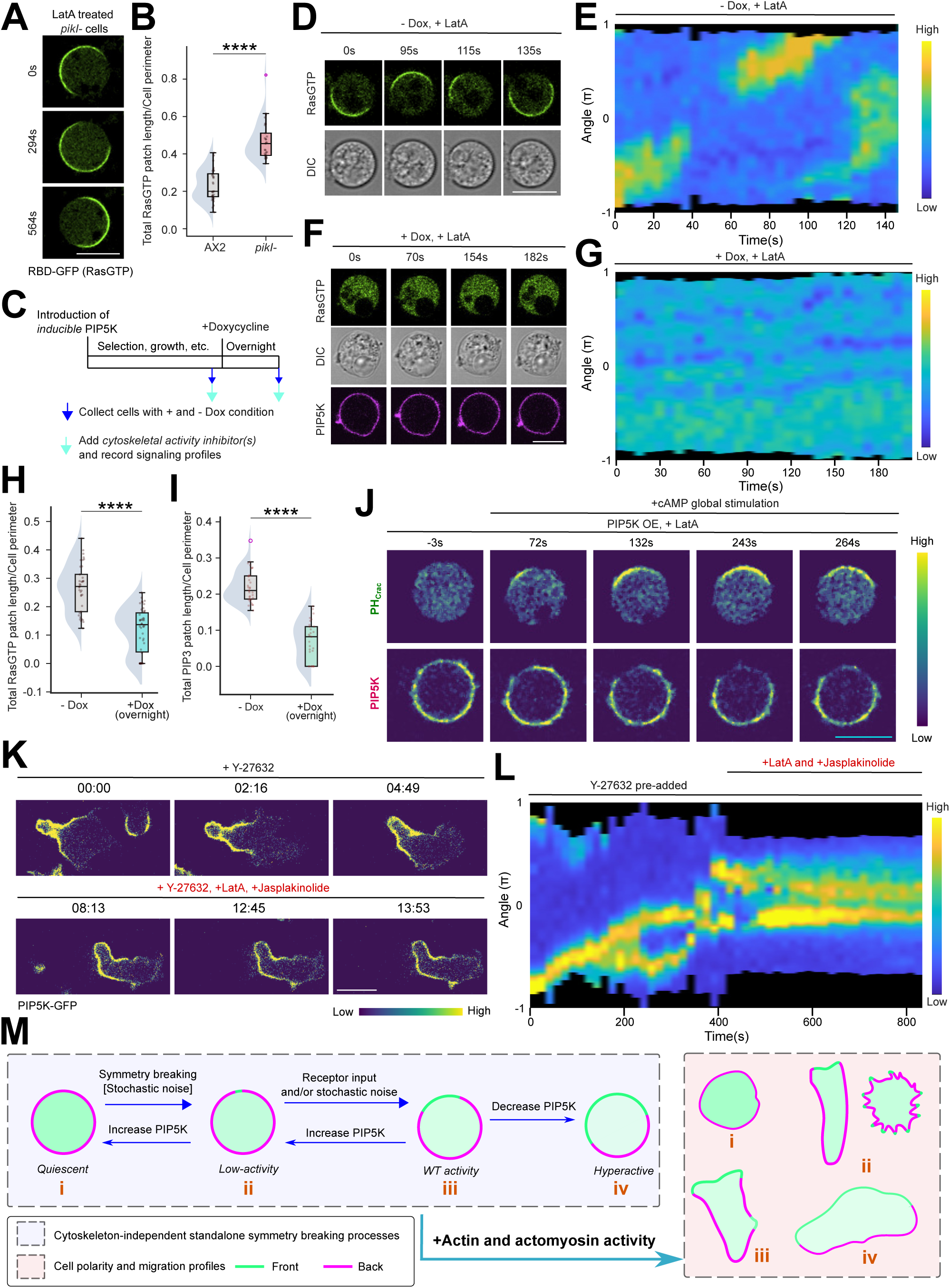
Mutually inhibitory actions of Ras-PIP5K can function in the absence of cytoskeletal dynamics. **(A)** Live-cell images of *pikI- Dictyostelium* cells expressing RBD-GFP, pre-treated with actin polymerization inhibitor Latrunculin A (final concentration 5µM) and caffeine (final concentration 4mM), showing large, asymmetric RasGTP activities on the plasma membrane, in the absence of actin cytoskeleton. **(B)** Quantification of overall RasGTP level on the cell membrane in AX2 or *piki- Dictyostelium* cells after Latrunculin A treatment, showing that RasGTP patches are significantly higher in *piki-* cell populations. The levels are quantified in terms of the total RBD-GFP patch length per unit perimeter length. Here, n_c_ = 33 for AX2 populations, and n_c_ = 29 for *piki-* cells populations. The p-values are computed by the Mann-Whitney U test. **(C)** Schematic of the experimental strategy involving docycycline-inducible promoter-driven PIP5K overexpression, followed by inhibition of cytoskeletal activities. **(D, F)** Live-cell images of *Dictyostelium* cells co-expressing RBD-GFP and Dox-inducible mRFP-PIP5K without Dox induction (D) or with overnight Dox induction (F). Cells were pre-treated with actin polymerization inhibitor Latrunculin A (final concentration 5µM) and caffeine (final concentration 4mM) for 20min, showing that Dox-treated cells have negligible RasGTP activities, compared to the untreated population. **(E, G)** Representative 360° membrane kymograph of RBD-GFP, corresponding to Figure 4D and 4F, respectively. **(H, I)** Quantifications of overall RasGTP level (H) or PIP3 level (I) on the cell membrane in *Dictyostelium* cells with Dox-inducible PIP5K overexpression system, without or with Dox induction, showing both RasGTP and PIP3 patches are reduced significantly upon overnight Dox induction, even in the absence of cytoskeleton. The levels are quantified in terms the total RBD-GFP patch length (H) or PH_crac_-YFP patch length (I) per unit perimeter length. Here, n_c_ = 37 (H, -Dox overnight), 46 (H, +Dox overnight), 35 (I, -Dox overnight), or 29 (H, +Dox overnight) cells. The p values are computed by the Mann-Whitney U test. **(J)** Live-cell images of *Dictyostelium* cells co-expressing PHcrac-RFP and Dox-inducible GFP-PIP5K with overnight Dox induction. Cells were pre-treated with actin polymerization inhibitor Latrunculin A (final concentration 5µM) and caffeine (final concentration 4mM) for 20min and then stimulated with 5µM cAMP at t=0, showing that, even in the absence of actin cytoskeletal dynamics, plasma membrane symmetry can be broken and PIP5K can dynamically localizes on the membrane domains where front signaling activities are low. **(K)** Live-cell images of differentiated HL-60 neutrophils expressing GFP-PIP5K. Cells were treated with 5µM ROCK inhibitor Y-27632, followed by 8µM Jasplakinolide and 5µM latrunculin B, showing PIP5K maintains back localization even in the absence of actin and actomyosin-based cytoskeletal dynamics. Time in min:sec format. **(L)** Representative 360° membrane kymograph (see “Methods” for details) of GFP-PIP5K in differentiated HL-60 neutrophils, corresponding to Figure 4K. Inhibitors were added as indicated on top. **(M)** Schematic showing different signaling states (blue box) and corresponding cellular morphologies (salmon box) in the absence or presence of cytoskeletal activities. The notations “i”-”iv” denotes the correspondence. Note that, increase and decrease in PIP5K level in cell can enable transition among signaling states and when cytoskeleton is engaged, it can regulate the cell migration mode, polarity, and morphologies.

Next, we asked whether the inhibitory effects of PIP5K expression depended on the cytoskeletal activity. We treated PIP5K overexpressing cells with Latrunculin A (Figure 4C) and noted that the typical oscillatory Ras and PIP3 patches were severely diminished in the overexpressing cells that were incubated overnight with Dox, compared to cells without Dox incubation (Figure 4D-I, S29A-D, Video S27). Furthermore, to study whether external stimulus which are known to activate signal transduction network can override PIP5K-mediated inhibition and thereby, break symmetry, we globally stimulated Latrunculin A-treated PIP5K overexpression cells with cAMP. We observed that PIP3 patches appeared in localized regions and importantly, PIP5K was depleted from those particular membrane domains (Figure 4J, S29E). Taken together, these data collectively suggest that the complementary patterns of PIP5K and Ras are tightly maintained in the absence of cytoskeletal dynamics, and this bistable interaction is fundamental to symmetry breaking.

It has been shown that neutrophils can maintain polarized form in the absence of cytoskeletal dynamics when treated with a cocktail of Jasplakinolide, Latrunculin B, and ROCK inhibitor Y-27632 (JLY) ^16,87^. Since actin turnover and myosin-based contractility are halted, the cells do not move (Figure 4K-L, S29F-G, Video S28), but they maintain their polarized sensitivity to receptor stimulation ^16^. Remarkably, we found that PIP5K persistently maintained its back localization in JLY-treated neutrophils (Figure 4L, S29F-G). Given our evidence that PIP5K is a negative regulator of Ras activity and downstream signal transduction activity, its persistent back localization can explain the reduced sensitivity at the back. These results prove that cytoskeletal dynamics are not required to maintain the insensitivity of the cell rear, although cytoskeletal structure is likely key in maintaining PIP5K at the rear.

Our data so far indicates that the inherent Ras-PIP5K bistability is critical for symmetry breaking on the plasma membrane (Figure 4M). In an initially quiescent cell, PIP5K is uniformly distributed on the membrane. Ras and other downstream signaling activities are absent. We propose that stochastic noises turn on a “spark” of Ras signaling, or depletion of PIP5K, on a local membrane domain, which initiates positive feedback. This mutually inhibitory positive feedback loop amplifies the initial event and the spread of activities. Increasing or decreasing PIP5K levels can drastically inhibit or activate Ras accordingly (Figure 4M). When connected to other downstream signaling and cytoskeletal pathways, this core circuit can tune polarity, and thereby, generate distinct morphological states and migratory modes, as we have observed (Figure 1-2).

### RasGEFX mediates PIP5K/PI(4,5)P2 driven inhibition of Ras activation

Our results have demonstrated that the RasGTP-PIP5K bistability acts as a core circuit regulating symmetry breaking and polarity, but it remains unclear how PIP5K/PI(4,5)P2 inhibits RasGTP. To address this question, we speculated that there are specific RasGEFs whose action is inhibited by high PIP5K/PI(4,5)P2 level (Figure 5A) and that if such RasGEFs were deleted, cells would subvert the phenotype driven by lowering PI(4,5)P2. To test this idea, we conducted a screen in *Dictyostelium,* which has 25 known RasGEFs (Figure 5B). A recent study performed a large-scale RasGEF overexpression screen where all RasGEFs have been hierarchically clustered based on their ability to affect Ras spontaneous excitability dynamics ^19^. Cluster 1 contained four RasGEFs, viz. RasGEFB/M/X/U, all of which, upon overexpression, enhanced spontaneous Ras activity significantly. Reasoning that one or more of these four GEFs mediates the enhanced Ras activity that is observed in membrane domains with low PI(4,5)P2, we focused our screen on this cluster.

**Figure 5.**
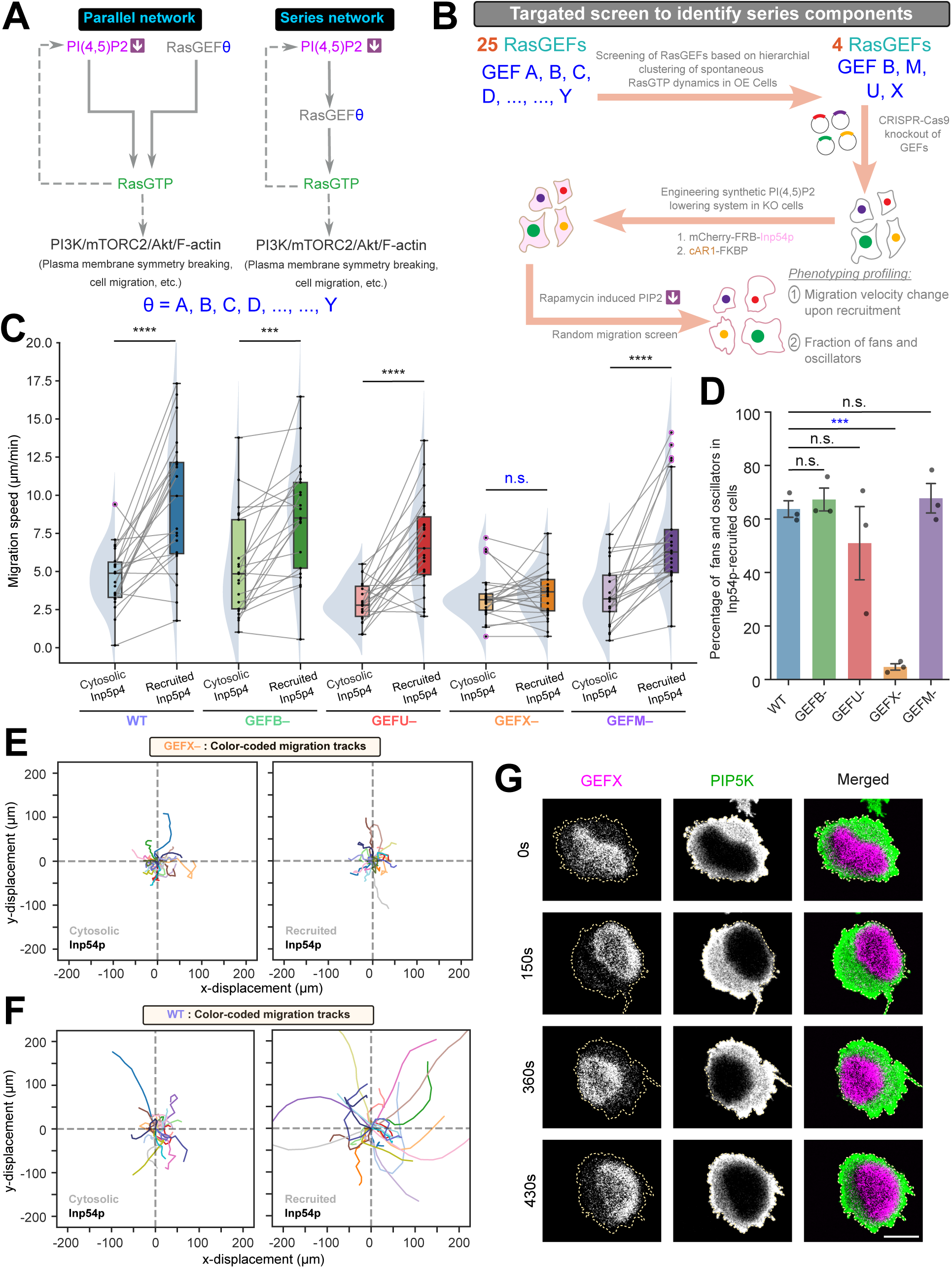
Genetic screen identifies that PIP5K/PI(4,5)P2 inhibits Ras activation on the plasma membrane by regulating RasGEFX profile. **(A)** Schematic showing two potential topologies by which Ras can be activated via lowered PI(4,5)P2 and/or via different RasGEFs. Note that *Dictyostelium* genome encodes 25 known RasGEFs, from A to Y. It is possible that a few of these GEFs act independently of decreased PI(4,5)P2 level to activate Ras (Parallel circuitry), while others directly facilitate the Ras activation in response to lowered PI(4,5)P2 level (Series circuitry). **(B)** Schematic showing the workflow of targeted genetic screening to identify the RasGEFs that directly activate Ras in response to a decrease in PI(4,5)P2. First, 4 GEFs were selected from the list of 25 GEFs, based on their ability to regulate spontaneous Ras excitable dynamics when overexpressed, by analyzing large-scale hierarchical clustering data from recent Iwamoto K et al. 2025 study. Next, a chemically inducible dimerization based synthetic PI(4,5)P2 lowering system (based on Inp54p, a yeast inositol polyphosphate 5-phosphatase that specifically hydrolyzes PI(4,5)P2) was incorporated in the *Dictyostelium* cells where one of these 4 GEFs were CRISPR knocked out. Finally, cell migration was profiled in these cells, before and after rapamycin induced acute PI(4,5)P2 lowering. **(C)** Quantification of average migration speed of wild-type AX2, *RasGEFB–*, *RasGEFU–*, *RasGEFX–*, and *RasGEFM– Dictyostelium* cells, before and after rapamycin induced recruitment of Inp54p from cytosol to membrane and acute PI(4,5)P2 lowering, demonstrating that knockout of RasGEFX completely abrogate PI(4,5)P2 lowering driven increase in migration speed, while knockout of any other GEFs has negligible effect on the phenotypic transition. Here, n_c_= 23 (WT), 21(*RasGEFB–*), 23 (*RasGEFU–*), 25 (*RasGEFX–*), 22 (*RasGEFM–*) cells and each cell has been tracked for t=22 min for before and after recruitment. For pairwise comparison between before and after recruitment, data from the same cell are connected by gray lines; the p values are computed by the Wilcoxon signed-rank test. **(D)** Quantification of percentage of cells migrating as fans and oscillator in wild-type AX2 and different RasGEF knocked out *Dictyostelium* cell populations, after rapamycin induced Inp54p recruitment driven PI(4,5)P2 lowering. Note that, even upon acute PI(4,5)P2 lowering, *RasGEFX– Dictyostelium* cells did not switch their migration mode to fans or oscillators. For definition of fans and oscillators, please see Figure S1 and S2. Here, each data point indicate one independent experiment where at least 40 cells were profiled. The p values by Welch’s t-test. **(E, F)** Centroid tracks of migrating *RasGEFX–* (E) and wild-type AX2 (F) *Dictyostelium* cells, before and after rapamycin induced recruitment of Inp54p. For pairwise comparison, tracks from the same cell are plotted in same color in before and after recruitment populations. Note that, while WT AX2 cells migrated much faster and switched their migration mode upon Inp54p recruitment, *RasGEFX–* cells did not exhibit any discernible changes. Here, n_c_= 25 (*RasGEFX–*) and 23 (AX2) cells and each cell has been tracked for t=22 min for before and after recruitment. All the tracks were reset to the same origin. **(G)** Ventral wave propagation in *Dictyostelium* cells co-expressing mCherry-RasGEFX and GFP-PIP5K, demonstrating that upon RasGEFX overexpression, PIP5K overexpressing cell can break symmetry and display propagating waves on the plasma membrane-cortex. Note that, GEFX and PIP5K maintain consistent complementarity as waves propagate.

We expressed a chemically inducible dimerization-based synthetic PI(4,5)P2 depletion system in cells lacking each of these RasGEFs and observed random migration before and after rapamycin-driven acute PI(4,5)P2 depletion (Figure 5B). In WT cells, decreasing PI(4,5)P2 significantly increased cell migration speed, and a majority of cells switched their migration mode to fans and oscillators (Figure 5C-D), as reported previously ^8,9^. Strikingly, in *gefx-* cells the effects of lowering PI(4,5)P2 were nearly abrogated (Figure 5E-F). There was a slight effect in *gefb-* cells, whereas the phenotypic changes caused by PI(4,5)P2 lowering were negligible in *gefm-* and *gefu-* cells (Figure 5C-F, S30A-C).

Results from these screens collectively suggest that synthetically lowered PI(4,5)P2 activates signaling and cytoskeletal networks primarily via RasGEFX. We found that RasGEFX itself localizes to membrane protrusions that have high RasGTP and low PIP5K levels, consistent with its role as a positive regulator of Ras activation (Figure S30D, Video S29). Importantly, even when cytoskeletal dynamics are inhibited, RasGEFX and RasGTP patches co-localized (Figure S30E, Video S29). Furthermore, we overexpressed RasGEFX and PIP5K, and we observed cell can break symmetry and maintain consistent complementarity during wave propagation (Figure 5G). Taken together, our data suggest that, as PI(4,5)P2 is generated by PIP5K, it strongly inhibits RasGEFX access, and consequently, Ras cannot be activated in the membrane domains where PI(4,5)P2 level is high. Whenever and wherever PI(4,5)P2 level decreases on the membrane, RasGEFX can be recruited and locally activate Ras and downstream signaling and cytoskeletal networks.

### Single-molecule measurements show activated Ras increases dissociation rate of PIP5K from the membrane

We have demonstrated that PIP5K/PI(4,5)P2 inhibits Ras activation via RasGEFX; however, how Ras modulates PIP5K localization remains unclear (Figure 6A). It is possible that activated Ras increases diffusion of PIP5K within the membrane or increases dissociation of the enzyme to the cytosol (Figure S31A-B) ^5,18^. One way to distinguish these is to apply a global stimulation, which activates Ras uniformly on the membrane. In response to a global stimulation, proteins that are considered to “shuttle”, clearly translocate to the cytosol, while others that compartmentalize by diffusing differentially in the membrane in a process called “dynamic partitioning”, stay on the membrane ^18^(Figure S32A). In our initial experiments, where we stimulated *Dictyostelium* cells with cAMP, we did not observe significant translocation of PIP5K to the cytosol (Figure S32B-C; Video S31, also see Figure S21-S22 for observing PIP5K dynamics during global receptor activation in different systems). Furthermore, selectively photoconverted membrane PIP5K molecules did not significantly appear in the cytosol (Figure S32D-G, Video S32), initially suggesting that PIP5K may re-localize by dynamic partitioning rather than shuttling.

**Figure 6.**
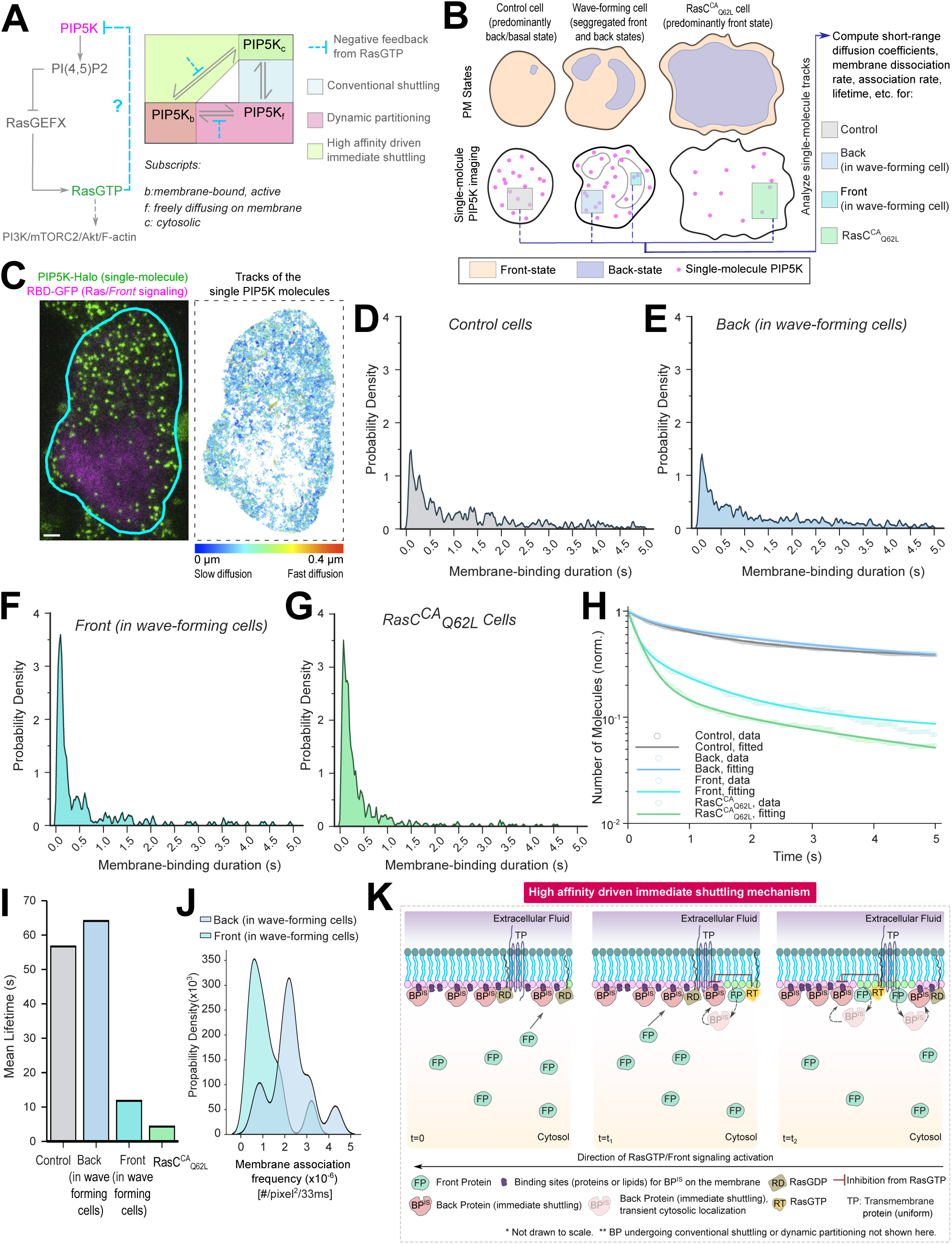
Single-molecule Imaging of PIP5K under different Ras activation levels reveals RasGTP inhibits PIP5K by limiting its binding to specific membrane domains via a rapid, differential dissociation and association based mechanism. **(A)** Left: Schematic of the core signaling network that trigger plasma membrane symmetry breaking and engages with downstream PI3K/mTORC2/Akt/F-actin network to properly polarize cell and generate localize protrusions. The mechanism of the blue inhibitory loop remained unknown and is therefore represented by a question mark. Right: The possible states that PIP5K can assume in the membrane and cytosol are shown, along with the potential mechanisms by which PIP5K activity on the membrane can be limited. For details on *Conventional shuttling* and *Dynamic Partitioning*, please see Figure S30. The *High affinity driven immediate shuttling* mechanism is delineated in Figure 6K. Blue inhibitory loop, consisting of actions from RasGTP driven negative feedback to PIP5K, can potentially act in different ways, as shown here. **(B)** Illustration of the multiscale imaging setup where PIP5K-Halo (TMR conjugated) is tracked at the single-molecule level while front signaling state regions on the membranes are tracked by tracking the coordinates of the RBD-GFP biosensor during ventral wave propagation. This assay was performed in different conditions where Ras is increasingly more activated, such as basal/quiescent-state cells (which exhibit predominantly back-state), cells with propagating waves (where membrane dynamically segregates into front and back state regions), and cells with constitutively active Ras (where is membrane is largely in front or activated state). From these conditions, short-range diffusion coefficients, lifetimes, membrane association and dissociation rates were computed for four different categories of membrane domains, as shown in the figure. **(C)** Left: Representative live-cell multiscale TIRF microscopy image of a *Dictyostelium* cell, where magenta shows the demarcated front signaling regions and green shows individual PIP5K-Halo-TMR molecules. Cell boundary is shown with the cyan outline. Right: The trajectories of single PIP5K molecules motion detected in 4s from the cell shown in left. The colormap below indicates the amount of diffusion. Scale bar indicates 2 um. **(D-G)** Kernel density estimation plots showing the probable density distribution of the membrane-binding durations of PIP5K in four different categories of membrane domains as shown in (B): control cells (D), back-state in wave-forming cells (E), front-state in wave-forming cells (F), and cells with constitutively active Ras (G). For all of these plots, *scott* was used as the bandwidth method and bw_adjust=0.04 was used to maintain consistent scaling. Note that, substantially more PIP5K molecules dissociate from the membrane domains as Ras activation increases. **(H)** Temporal profile of dissociation of PIP5K molecules from four categories of membrane domains (as depicted in (B)), demonstrating that the dissociation rate of PIP5K molecules from membrane increases as Ras gets increasingly more active. Means are shown and lines represent the fitted data. **(I)** Mean lifetimes of PIP5K in our categories of membrane domains (as depicted in (B)). **(J)** Kernel density estimation plots showing the probability density distribution of the membrane association of PIP5K in back vs front state regions in cells, demonstrating PIP5K molecules bind more effectively to the back-state, compared to the front-state. Here *scott* was used as the bandwidth method and bw_adjust=0.5 was used. **(K)** Schematic showing the suggested *High affinity driven immediate shuttling* mechanism. Lipid headgroups associated with the back-state and front-state are shown in mauve color and green color, respectively. At t=0, the membrane is in complete back-state and wave of signaling activation initiated at the rightmost point and as time passed (t_2_>t_1_>0), wave propagated from right to the left. As wave propagates and Ras switches from GDP to GTP bound state in specific membrane domains, front proteins translocate to those membrane domains from the cytosol. Simultaneously, in those domains, RasGTP provides negative feedback to membrane binding of BP^IS^ (PIP5K is a BP^IS^ molecule) and force it to transiently move to cytosol; due to presence of oversaturating amount of binding sites (shown in dark violet), these dissociated BP^IS^ molecules quickly bind to other parts of the membrane. Thus, Ras maintain complementarity with PIP5K to facilitate mutual antagonism-based signal amplification and to spatiotemporally regulate signaling, in a mechanism distinct from *Conventional shuttling* and *Dynamic Partitioning*. TP molecules are shown as fixed, fiducial component. Solid arrows are shuttling between the membrane and cytosol and the dashed arrows are showing transient immediate shuttling.

To test whether PIP5K diffuses heterogeneously in the front and back state regions of the membrane, we designed a multiscale imaging setup where we tracked front state using a RasGTP biosensor and tracked single-molecule diffusion of PIP5K on the other channel ^88^. We tracked diffusion under various experimental conditions, such as in *Dictyostelium* cells which are quiescent, wave-forming, or RasC_Q62L_ expressing cells, which have increasingly more Ras activities (Figure 6B). PIP5K-Halo was observed as diffusive fluorescent spots (Figure S33A). We verified that the individual fluorescent spots of PIP5K-Halo-TMR detected using TIRF microscopy represent single molecules, as demonstrated by single-step photobleaching patterns (Figure S33, Video S33) and the distribution of fluorescence intensities of the spots (Figure S33C-G). Subsequently, we measured the movement profiles of individual PIP5K molecules in both the front- and back-state regions of the membrane using single-molecule tracking (Figure 6C). Surprisingly, tracking shows that the short-range diffusion coefficient of PIP5K is not substantially different between front and back states of the membrane, and furthermore, was slightly slower in RasC_Q62L_ expressing cells (Figure S34A-H), questioning our dynamic partitioning hypothesis.

Next, we measured the lifetime of PIP5K molecules under all experimental conditions. The lifetime in the front state regions and RasC_Q62L_ expressing cells was significantly shorter than in control cells and back state regions (Figure 6D-I). Furthermore, membrane association frequency of PIP5K in back state regions was nearly three times higher than in the front state regions (Figure 6J). Consistently, a significant amount of PIP5K was moved to the cytosol of cells expressing RasC_Q62L_ (Figure S34I). Taken together, these data suggest that the dissociation constant in activated front state regions is nearly two orders of magnitude larger than in the back state regions. This large difference in dissociation constant will lead to the depletion of the molecules at the front and enrichment at the back. This explains why PIP5K molecules dynamically re-localize away from the activated regions (Figure 6K). The affinity of PIP5K molecules for the membrane must be extremely high or there must be a large excess binding sites on the membrane since so little amount of enzyme is found in the cytosol under normal physiological circumstances.

### The stochastic, reaction-diffusion model explains PIP5K driven tuning of signaling activities

Figure S35A summarizes the molecular interactions that we propose to account for symmetry breaking and signal amplification, leading to polarity establishment. The core of the network consists of the mutual inhibition between PI(4,5)P2 and RasGTP. PI(4,5)P2 inhibits Ras activation, and in turn, activated Ras lowers PI(4,5)P2 by removing PIP5K from the membrane. The total amount of PIP5K is constant on the plasma membrane. This mutual antagonism-based positive feedback, in conjunction with delayed negative feedback from Akt, confers biochemical excitability to the core loop. These events can occur independently of cytoskeletal activities. However, under normal physiological conditions, Akt triggers branched actin polymerization, which feeds back to Ras. Furthermore, increased PI(4,5)P2 promotes myosin, which provides negative feedback to Ras.

To assess the feasibility of these schemes *in silico*, we developed a mathematical model that combines the key molecular interactions with level-set frameworks, which move the cell boundaries ^8,89,90^. In WT conditions, stochastic noise triggers transient Ras activation (Figure 7), which spreads as V-shaped waves in kymographs. PIP5K and PI(4,5)P2 are depleted from those areas, whereas Akt follows them (Figure 7, S35). Consequently, branched actin appears in the regions where Ras is activated, and actomyosin is depleted (Figure S35). When PIP5K is overexpressed, Ras and branched actin are confined to small patches, or extinguished, and the entire cortex is shifted toward the back state. When PIP5K is absent, Ras is highly activated, and, once triggered, signal transduction events and branched actin spread across most of the cortex and oscillate. With increased polarity, the signaling remains on across half of the cell. When the feedback loops from actin and myosin are removed, this core symmetry breaking network remains excitable (Figure S36).

**Figure 7.**
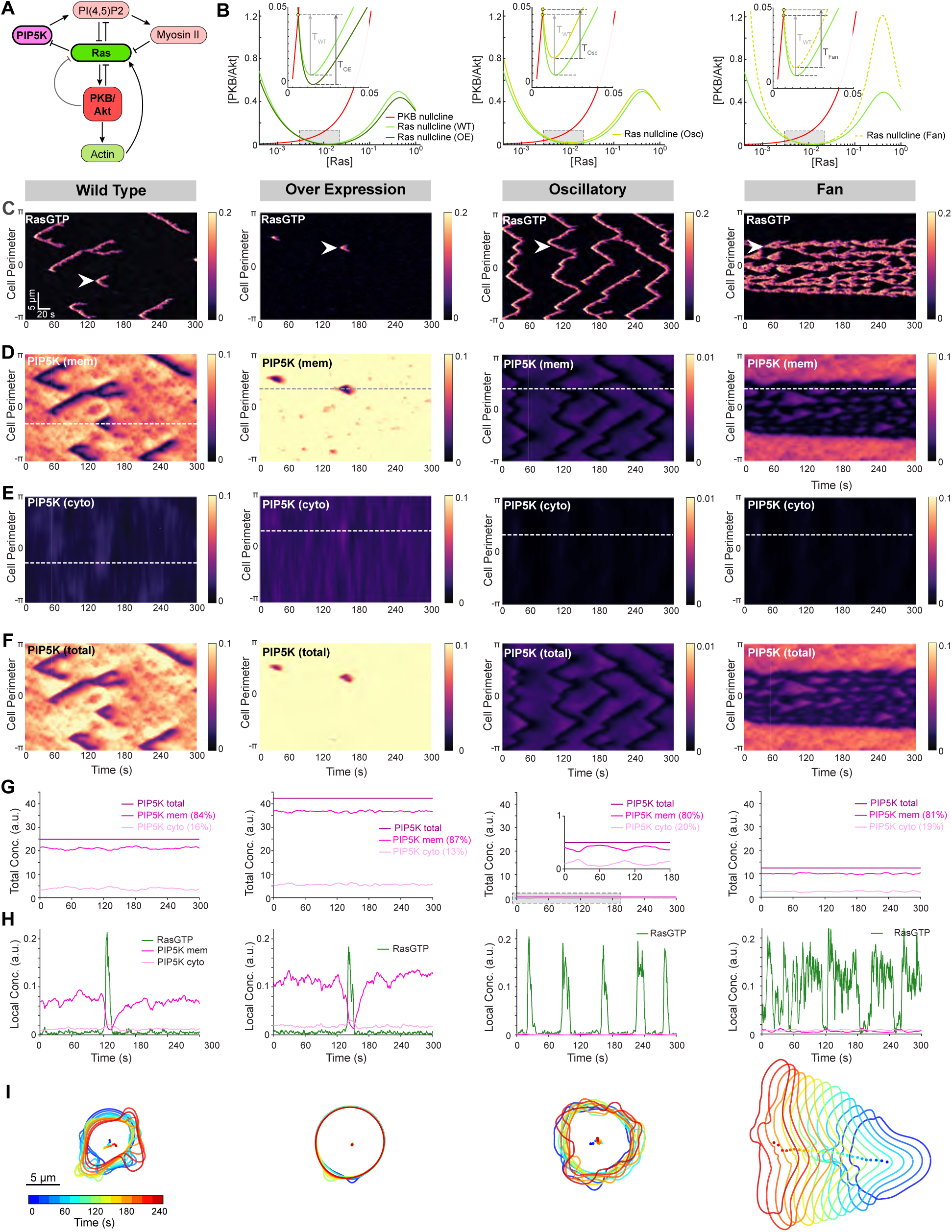
A stochastic reaction-diffusion model predicts signaling state transition-driven cellular morphological changes. **(A)** Schematic illustrating the key signaling interactions included in the model. **(B)** Phase-portraits showing nullclines for Ras (shades of green) and PKB/Akt (red) under different perturbation scenarios. Yellow markers denote equilibrium points; the inset shows the firing thresholds (𝑇_∗_), indicated by arrows. **(C–F)** Kymographs of molecular dynamics over a 300 s simulation: **(C)** Ras, **(D)** membrane-associated PIP5K, **(E)** cytosolic PIP5K, **(F)** total PIP5K. Columns correspond to different conditions: wild type (1st), PIP5K overexpression (2nd), *piki-* oscillatory phenotype (3rd), and *fan* phenotype (4th). **(G)** Time series of total PIP5K (dark purple), membrane-associated (purple), and cytosolic (light purple) fractions, summed across the cell perimeter. Total PIP5K remains conserved over time, but differs across phenotypes, mimicking the experimental manipulations. **(H)** Local concentration profiles of Ras (green), membrane-associated (purple), and cytosolic (light purple) PIP5K at the spatial location marked by the arrow in **(C)** and dashed lines in **(E,F)**. These dynamics highlight a causal relationship: Ras activation correlates with decreased membrane association and a transient increase in cytosolic PIP5K. **(I)** Simulated cell morphologies for control and perturbed conditions, illustrating phenotypic differences predicted by the model. Colors indicate different time points in the simulation.

To visualize the effects of these perturbations on cell migration, we developed a system in which the output (i.e., Akt, branched actin) drove the expansion of the viscoelastic boundary. As shown in Figure 7G and Video S34, in WT cells, each wave of Ras activation creates a bilateral protrusion with the cup-like morphology. The cup is formed because, as evidenced by the kymograph, activation occurs at the spreading tips, while inhibition occurs at the valley. When PIP5K is high, protrusions become smaller or disappear entirely and cells stop migration. When PIP5K is absent, PI(4,5)P2 levels are low, protrusions are wide, and cells switch their migratory behaviors from amoeboid to oscillatory or fan-shaped phenotypes.

## Discussion

Figure 8A illustrates the network in greater detail, highlighting the molecular interactions that we propose to account for symmetry breaking and signal amplification, ultimately leading to the establishment of polarity. PI(4,5)P2 inhibits Ras activation by preventing RasGEFX recruitment to the membrane. Activated Ras increases the dissociation rate of PIP5K from the membrane, perhaps by reducing the level of anionic lipids ^17,91^. Activated Ras couples to the PI3K/mTORC2/Akt pathway, triggering branched actin polymerization and inhibiting myosin assembly. Increased PI(4,5)P2 promotes linear actomyosin. Branched and linear actin provide positive and negative feedback loops to Ras, respectively. As shown in Figure 8B, in the basal state, PIP5K is at the membrane, PI(4,5)P2 levels are high, RasGEFX is in the cytosol, Ras-GTP activity is low, and the cell cortex is mostly actomyosin. Due to local stochastic fluctuations, some RasGEFX molecules approach the membrane, or some PIP5K molecules spontaneously dissociate from it. Ras becomes activated, leading to further PIP5K dissociation, which lowers PI(4,5)P2 and initiates a positive feedback loop. Meanwhile, more RasGEF molecules are recruited to the membrane, resulting in increased Ras activation. Actomyosin is replaced by branched actin in this region. Crucially, dissociated PIP5K molecules rapidly bind to other regions of the membrane, preventing RasGEFX from accessing the membrane and activating Ras and initiating a new cycle there.

**Figure 8.**
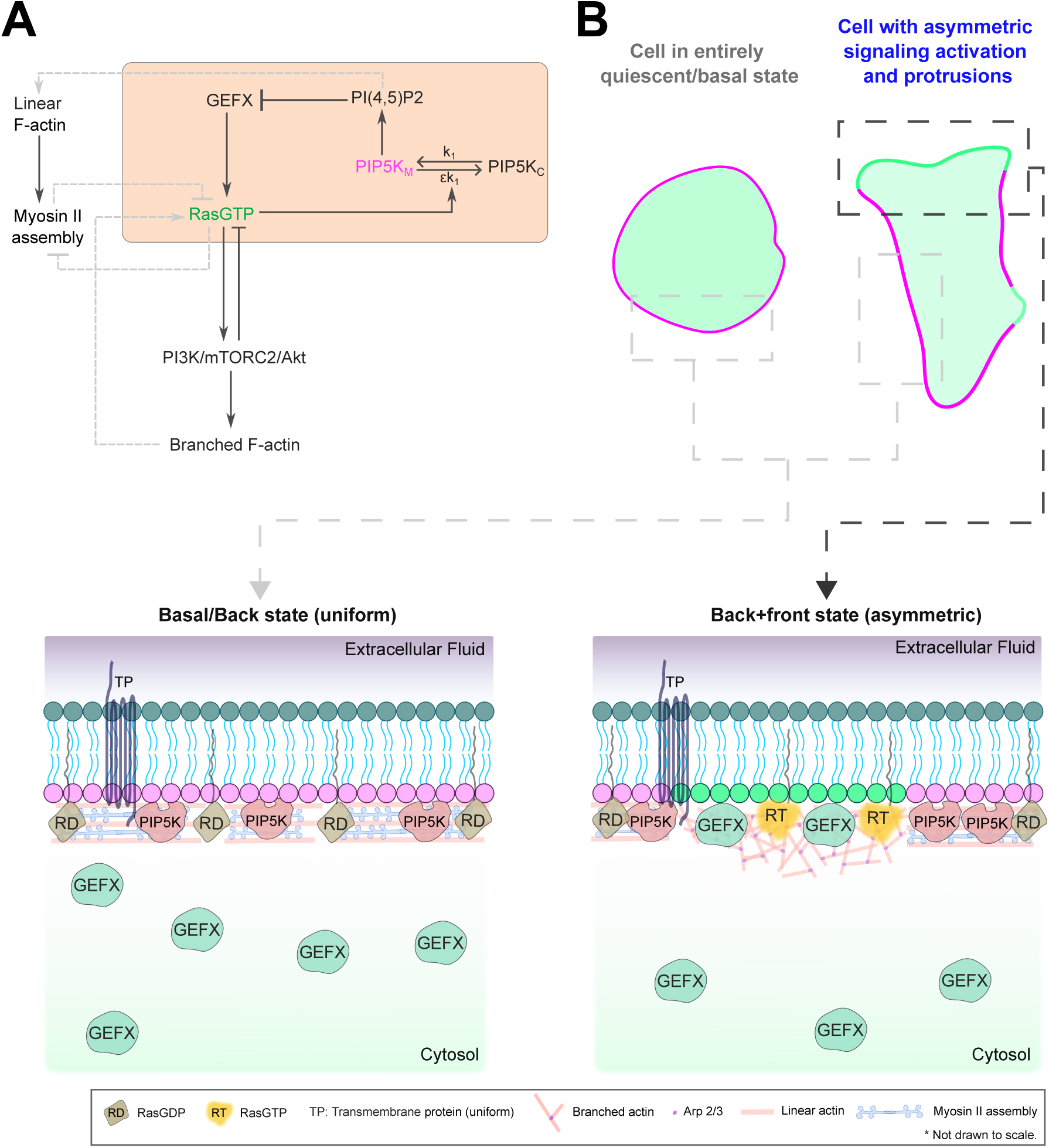
Schematic illustration showing the biophysical and biochemical bases of plasma membrane symmetry breaking. **(A)** Signal transduction network that regulate cell polarization and migration. Core signaling components that define the fundamental symmetry breaking and signal amplification is highlighted with a peach colored box. M and C subscripts denote membrane and cytosol, respectively. Here k_1_ is the membrane binding rate of PIP5K, ε is a small number that can be increased by RasGTP activity. Grey connections indicates positive and negative interactions with cytoskeletal components that are not essential for the symmetry breaking of the signaling network. **(B)** Note that, as a cell transitions from basal/back/inactivated to front/protrusions/activated state via a symmetry transition, PIP5K molecules are removed from the front-state regions via RasGTP activities (which depletes its binding sites, presumably a combination of anionic lipids) and those dissociated molecule re-associate with the remaining or new back-state regions of the membrane. RasGDP is converted into RasGTP on the membrane via the action of recruited RasGEFX. The head groups of the inner leaflet lipids that form the back or basal-state (such as PI(4,5)P2, PI(3,4)P2, PS, PA, etc.) are shown in magenta and the head groups of the front-state associated lipid molecules are shown in green. Indicated transmembrane protein does not localize preferentially.

Several elegant models for directional sensing and cell polarity have been proposed, but few of these explain the basic mechanisms of spontaneous symmetry breaking. Models envisioning some type of local <u>e</u>xcitor and a more <u>g</u>lobal inhibitor, generally referred to as LEGI models, allow local responses while blocking responses elsewhere ^24,25,92^. In one version the levels of the regulators are tied to receptor occupancy. ^26,92^ Other recent examples employ membrane ^29,30^ or cortical ^93^ tension, which, as protrusions form, increases rapidly and globally, preventing protrusions elsewhere or membrane-cortical detachment, which specifically releases global inhibition at the front of the cell ^33,34,94^. Other models involve membrane bending, where recruitment of F-BAR-domain-containing proteins have sought to explain wave propagation ^95^. Since all these systems require specific external cues or cytoskeletal activity, they cannot explain the initial fundamental symmetry breaking steps, which in our model stems from differential localizations and mutual inhibitions between activated Ras and PIP5K.

The concept of mutual inhibition-driven symmetry breaking has been theorized from the systems perspective and tested in synthetic or *in vitro* systems, but only a few models have focused on specific molecules in living cells. These include mutual inhibition between Rac1 and RhoA^96^, branched actin and linear actin/actomyosin^97^, or PI3K/PIP3 and PTEN^7^. The first two models, again, rely on cytoskeletal activity, so while these interactions may be important downstream, they cannot account for the initial steps. Loss of PIP3 has only transient effects on migration, but over production of PIP3 in *pten-* cells inhibits signaling, so that this model deserves consideration. However, it seems secondary to the highly sensitive core circuit presented here where Ras or PIP5K alters multiple downstream signaling pathways in propagating cortical waves, extended protrusions, macropinocytosis, polarized migration, and chemotaxis.

Moreover, these features are conserved across evolutionally diverse cell types and organisms. Thus, this Ras-PIP5K bistability circuit is potentially the most fundamental concept to explain symmetry breaking and control a plethora of subsequent cell physiological processes. Our study shows that PI(4,5)P2 serves as one of the strongest negative regulators of Ras activity identified to date. Although various regulatory pathways for PI(4,5)P2 synthesis and degradation have been reported, our findings, together with prior literature, underscore the overarching importance of PIP5K in setting PI(4,5)P2 levels. In principle, PIP5K and PI(4,5)P2 can attenuate Ras signaling either by recruiting RasGAPs or by competing with RasGEFs, thus inhibiting Ras activation. The spontaneous appearance of RasGEFs, primarily GEFX in *Dictyostelium*, on protrusions and the requirement of GEFX for the hyperactivation of Ras induced by PI(4,5)P2 lowering, suggests that high PI(4,5)P2 may inhibit Ras by preventing the access of GEFs to Ras. Some of the key questions for future research include: How are GEFs regulated by PI(4,5)P2? How does activated Ras increase the dissociation of PIP5K from front regions? Scores of additional network components display similar localization dynamics to RasGTP and PIP5K and many have been shown to regulate the network, but none of these have displayed as penetrant phenotypes.

PI(4,5)P2 itself has been broadly implicated in cell migration and cancer signaling, serving not only as the precursor for PLC-dependent generation of IP3 and DAG, which are classic activators of the Ca2+/PKC pathway, but also as a modulator of numerous actin-binding proteins. Despite this, we found that PLC and PI3K, both of which degrade PI(4,5)P2, play only a minimal role in Ras activation and actin polymerization. Moreover, while a few studies report that PI(4,5)P2 levels rise at the front, the collective effects of PI(4,5)P2 on the actin cytoskeleton is hard to predict *in vivo* since the reported effects *in vitro* span both activating and inhibitory roles ^98–109^. Our data, however, provide clear evidence that PIP5K acts as a negative regulator in multiple physiological contexts across diverse cells and organisms.

These observations have significant implications for understanding disease and developing therapeutic strategies. On the surface, one might predict that PIP5K acts as an oncogene, given that PI(4,5)P2 is a substrate for PI3K and PLC, which promote oncogenic signaling axes, such as Akt and PKC pathways ^110–115^. However, overexpression of PIP5K robustly suppresses Ras and PI3K activity and reduction of PI(4,5)P2 clearly elevates PIP3 and hyperactivates Ras. As Ras and Akt are among the most important oncogenes, according to our results, one would expect that PIP5K to be a tumor suppressor. Consistently, overexpression of PIP5K in cells and organoids strongly inhibits migration and dissemination. Therefore, increasing PIP5K, rather than inhibiting it, in cancer cells might be an effective therapeutic approach.

## Acknowledgments

We thank all members of the Devreotes, Iglesias, and Robinson laboratories (Schools of Medicine and Engineering, JHU) for helpful discussions and for providing resources. We thank Jonathan Kuhn and Yiyan Lin (Devreotes Lab), and Elena Parajon (Robinson Lab) for providing constructs. We thank Wakiko Iwata (Iijima Lab, School of Medicine, JHU) for sharing experimental reagents and advice for performing spheroid assay. We thank Orion Weiner (UCSF) for providing HL-60 cell line. We thank Sean Collins (UC Davis) for providing transposon plasmids. We thank Gerald Hammond (U Pittsburgh) for providing the cytosolic PIP5K1C construct. We thank Stephen Gould (School of Medicine, JHU) for help with instrumentation. We appreciate services of DictyBase and Addgene for providing plasmids. This work was supported by NIH grant R35 GM118177 (to P.N.D.), DARPA HR0011-16-C-0139 (to P.A.I. and P.N.D.), AFOSR MURI FA95501610052 (to P.N.D.), as well as NIH grant S10OD016374 (to S. Kuo of the JHU Microscope Facility).

## Author Contributions

Y.D., T.B., and P.N.D. conceived and designed the project. Y.D. engineered constructs/stable cell lines. Y.D. and T.B. designed and executed the majority of the experiments. T.B. and Y.D. performed quantification and the majority of the data analysis. S.M. and M.U. conducted single-molecule imaging assays, performed the relevant analysis, and helped with RasGEF screening. L.A.K. and N.H.R. conducted primary T cell migration experiments and relevant quantifications. D.S.P. and H.Z. provided resources for mammalian experiments. J.B. and Y.L. made a few constructs and helped with cell culture. D.B. and P.A.I. developed computational models, and D.B. performed simulations with inputs from P.A.I., P.B., P.N.D., T.B., and Y.D.. T.B., Y.D., and P.N.D. wrote and revised the manuscript with comments from the rest of the authors. P.N.D. supervised the overall study.

## Competing Interests

The authors declare no competing interests.

## Methods

### Preparation of reagents and inhibitors

Rapamycin (Sigma-Aldrich, 553210) was dissolved in DMSO to prepare the 5 mM stock solution. Then,1 µl aliquots were diluted 1:100 in the development buffer (DB) to a 10x concentration (50 µM). CK666 (Sigma-Aldrich, 182515) was dissolved in DMSO to prepare the 100 mM stock solution. Then, on the day of the experiment, 0.5 µl aliquots were diluted 1:100 in the warm development buffer (DB) to a 10x concentration (1 mM). 237 µM latrunculin A in ethanol (Cayman, 10010630) was diluted in the development buffer (DB) to prepare the 50 µM (10x) stock solution. 10 mM latrunculin B (Millipore Sigma, 428020) stock solution was made in dimethylsulfoxide (DMSO). Jasplakinolide (Sigma-Aldrich, 420127) was available as a ready-made 1 mM stock. Folic acid (Sigma-Aldrich, 329823065) was dissolved in sterile water, with the addition of 2 M NaOH, to prepare the 1.25 mM stock solution. Caffeine (Sigma-Aldrich,1085003) was dissolved to 1 M in sterile water and then diluted to a 10x concentration (40 mM) in the development buffer (DB). Fibronectin (Sigma-Aldrich, F4759-2MG) was dissolved in 2ml sterile water, followed by dilution in 8ml Ca^2+^/Mg^2+^ free PBS to prepare the 200 µg ml^−1^ stock solution. N-Formyl-Met-Leu-Phe (fMLP; Sigma-Aldrich, 47729) was dissolved in DMSO (Sigma-Aldrich, D2650) to prepare the 50 mM stock solution. Then, 1 µl aliquots were diluted in RPMI to a 10x concentration (2 µM). Phorbol 12-myristate 13-acetate (PMA; Sigma-Aldrich, P8139) was dissolved in DMSO to prepare the 1 mM stock solution. Then, 1 µl aliquots were diluted in RPMI to a 10x concentration (320 nM). FKP-(D-Cha)-Cha-r (ChaCha peptide, Anaspec; 65121) was dissolved in PBS to prepare the 2.5 mM stock solution. Then, 1 µl aliquots were diluted in DMEM to a 10x concentration (320 nM). A stock solution of EGF (Sigma-Aldrich, E9644) was prepared by dissolving it in 10 mM acetic acid to a final concentration of 1 mg ml^−1^. The anti-BSA mouse monoclonal antibody was acquired from Sigma-Aldrich (SAB4200688, clone BSA-33). Blebbistatin (Peprotech, 8567182) was dissolved in DMSO to prepare the 50mM stock solution. Then, 1 µl aliquots were diluted 1:100 in the development buffer (DB) to a 10x concentration (500 µM). 5 mM Y-27632 (Sigma-Aldrich, 688001) was dissolved in DMSO to prepare the 5 mM stock solution. Then, 1 µl aliquots were diluted in RPMI to a 10x concentration (100 µM). Hygromycin B (Thermo Fisher Scientific, 10687010) or G418 sulfate (Thermo Fisher Scientific, 10131035) was purchased as 50 mg ml^−1^ stock solution. Then, 10 µl Hygromycin B or G418 sulfate was added in a 10ml cell culture. Blasticidin S (Sigma-Aldrich, 15205) or puromycin (Sigma-Aldrich, P8833) was dissolved in sterile water to prepare the stock solutions of 10 mg ml^−1^ or 2.5 mg ml^−1^, respectively. Then, 10 µl Blasticidin S or 4 µl puromycin was added in a 10ml cell culture. Doxycycline hyclate (Sigma-Aldrich, D9891-1G) was dissolved in sterile water to prepare the stock solution of 5 mg ml^−1^. Then, 100 µl Doxycycline hyclate was added in a 10ml cell culture before the experiment. TRITC-dextran or FITC-dextran (Sigma-Aldrich, T1162) was dissolved in sterile water to prepare the 50 mg ml^−1^ stock solution. Then, the stock solution was further diluted to 2 mg ml^−1^ in the HL5 culture medium before the experiment. All stock solutions were aliquoted and stored at −20 °C.

### Cell culture

Wild type *Dictyostelium discoideum* cells of the AX2 strain (dictyBase, strain ID DBS0235521) were obtained from the R. R. Kay laboratory (MRC Laboratory of Molecular Biology) and cultured in HL5 medium at 22 °C for a maximum of 2 months after thawing from frozen stock. The *pip5k-* (piki-) cells were purchased from Dictybase (strain ID DBS0350270), which was originally generated by Fets et al. (Strain ID HM1513) ^60^. pikI− cells were grown on a *Klebsiella aerogenes* lawn on an SM plate and transferred to HL5 medium supplemented with heat-killed *Klebsiella aerogenes* before the experiment. *gefX-, gefB-, gefU-, and gefM-* cells were generated at the Ueda lab, as described previously_19_. *Gβ-* cells were generated in the Devreotes Lab previously ^80^. abnABC- cells were generated previously in the Devreotes Lab ^116^. Growth phase cells were used for all the experiments. GFP-arpC cells were purchased from Dictybase (strain ID DBS2036065).

The human HL-60 cell line (ATCC CCL-240; RRID:CVCL_0002) was obtained from the Weiner laboratory (UCSF) and cultured in RPMI 1640 medium (Gibco, 22400-089) supplemented with 15% heat-inactivated fetal bovine serum (Thermo Fisher, 16140071). To obtain migration-competent neutrophils, WT or stable lines were differentiated in the presence of 1.3% DMSO with a total of 1.5 million cells over 5-7 days_117_. Differentiated cells are an effective model to study human neutrophils ^118^. To differentiate HL-60 cells into macrophages, cells were incubated with 32 nM PMA for 48-72h ^119^. Cells were grown in humidified conditions at 5% CO2 and 37 °C.

RAW 264.7 macrophage-like cells were obtained from the N. Gautam laboratory (Washington University School of Medicine in St. Louis). RAW 264.7 cells were cultured in Dulbecco’s modified Eagle’s medium (DMEM) containing 4,500 mg l^−1^ glucose, l-glutamine, sodium pyruvate and sodium bicarbonate (Sigma-Aldrich, D6429), supplemented with 10% heat-inactivated fetal bovine serum (ThermoFisher Scientific, 16140071) and 1% penicillin–streptomycin (ThermoFisher Scientific, 15140122). When 70–90% confluency was reached, RAW 264.7 cells were gently lifted using cell scrapers (SARSTEDT, 83.1830) and subcultured using 1:5-1:10 split ratio.

MDA-MB-231 breast cancer cell lines were obtained from M Iijima laboratory (School of Medicine, Johns Hopkins University). These cells were cultured in DMEM medium with high glucose and GultaMAX^TM^ (ThermoFisher Scientific, 10566024), supplemented with 10% heat-inactivated fetal bovine serum (ThermoFisher Scientific, 16140071) and 1% penicillin–streptomycin (ThermoFisher Scientific, 15140122). When 70–90% confluency was reached, MDA-MB-231 cells were lifted using 0.05% Trypsin-EDTA (ThermoFisher Scientific, 25300054) and subcultured using 1:5-1:10 split ratio.

All mammalian cells were grown in humidified conditions at 5% CO2 and 37 °C. All the experiments were conducted using low passage number cells.

### DNA constructs

All DNA oligonucleotides were purchased from Sigma-Aldrich. *Dictyostelium* PIP5K (referred to as PI5K below) was PCR amplified from genomic DNA and cloned into the pCV5-GFP expression plasmid to generate PI5K-GFP (pCV5). Two-point mutations K681N and K582N were introduced to PI5K-GFP (pCV5) to generate PI5K (K581N, K682N)-GFP (pCV5). PI5K-GFP was then PCR amplified and cloned into doxycycline-inducible pDM335 plasmid (dictyBase, ID no. 523) using BglII/SpeI restriction digestion to generate mRFPmars-PI5K (pDM335). mRFPmars-PI5K was than PCR amplified and cloned into doxycycline-inducible pDM359 plasmid (dictyBase, ID no. 518) using BglII/SpeI restriction digestion to generate mRFPmars-PI5K (pDM359). mRFPmars was then replaced with EGFP to generate EGFP-PI5K (pDM359). EGFP was then replaced with KikGR to generate KikGR-PI5K (pDM359). DNA sequences encoding the 1-315aa, or 316-718aa, or 369-718aa or 1-718aa amino acids of PI5K, were PCR-amplified from PI5K-GFP (pCV5) and cloned into mCherry-FRB-MCS (pCV5) expression plasmid (previously created in the Devreotes Lab) using NheI/XhoI restriction digestion to generate mCherry–FRB-PI5K (1-315aa) (pCV5), mCherry–FRB- PI5K (316-718aa) (pCV5), mCherry–FRB-PI5K (369-718aa) (pCV5) and mCherry–FRB-PI5K (1-718aa) (pCV5). DNA sequences encoding the 316-718aa amino acids of PI5K, were PCR amplified and cloned into mRFPmars-SspBR73Q-MCS (pCV5) using NheI/NotI restriction digestion to generate mRFPmars-SspBR73Q-PI5K (316-718aa). DNA sequences encoding the 301-718aa amino acids of PI5K, were PCR amplified and cloned into pCV5- GFP expression plasmid to generate PI5K (301-718aa)-GFP (pCV5). PH-PLCδ-YFP (pCV5), CynA-KikGR (KF2), RBD-EGFP (pDM358), PHcrac-RFP (pDRH), LimEΔcoil- mCherry (pDM181), PHcrac-YFP (pDM358), LimEΔcoil-RFP (pDRH), cAR1-FKBP-FKBP (pDM358 or pCV5), LimE-GFP-FKBP-FKBP (pDM358), mCherry-FRB-MHCKC (pCV5), and PKBR1_N150_-iLiD (pDM358) were previously created in the Devreotes Lab. Myosin II-GFP (pDM181) was obtained from the Robinson laboratory (School of Medicine, JHU). PAK1(GBD)-YFP (pDEXH) was a gift from the C. Huang lab (School of Medicine, JHU). RBD-RFP (pDM358) GFP-ABD120 (pDXA) was purchased from Dictybase (ID no. 472).

For transient transfection in RAW 264.7 macrophage-like cells and MDA-MB-231 cells, PH_Akt_-mCherry and CIBN-CAAX (Addgene #79574) were obtained from Devreotes Lab stock. EGFP-PIP5K1B (Addgene #202722), GFP-PIP5K1γ90 (Addgene #22299) TagBFP-FKBP-PIP5K1C (Addgene #220090) (cytosolic PIP5K1C), and mCherry-NES-SspB-MCS-iLiD-CAAX (Addgene #173869) were purchased from Addgene. mCherry was replaced with Crimson fluorescent protein, and a point mutation R73Q was introduced to SspB to generate Crimson-NES-SspBR73Q-MCS-P2A-iLiD-CAAX. Cytosolic PIP5K1C was PCR amplified and cloned into the XhoI/EcoRI sites of the Crimson-NES-SspBR73Q-MCS-P2A-iLiD-CAAX expression plasmid to generate Crimson-NES-SspBR73Q-PIP5K1C-P2A-iLiD-CAAX (Opto-RecP). Venus-iLiD-Mito (Addgene #60413) was purchased from Addgene. iLiD-CAAX was replaced with iLiD-Mito to generate Crimson-NES-SspBR73Q-MCS-P2A-iLiD-Mito. MCS was replaced with full-length PIP5K1B to generate Crimson-NES-SSPBR73Q-PIP5K1B-P2A-iLiD-Mito (Opto-SeqP).

For stable lines in HL-60 cells, stable lines co-expressing LifeAct–miRFP703 (pLJM1) (Addgene #201750) and CIBN-CAAX (pLJM1) (Addgene #201749) or expressing RFP-PHAkt (pFUW2) were previously created in the Devreotes Lab ^120^. EGFP-PIP5K1B and EGFP-PIP5K1C were PCR amplified and cloned into the BspEI/SalI sites of the PiggyBac transposon plasmid to generate EGFP-PIP5K1B (pPB) and EGFP-PIP5K1C (pPB). Two-point mutations K359N, K260K or K407N, K408N were introduced to EGFP-PIP5K1B (pPB) or EGFP-PIP5K1C (pPB), to generate EGFP-PIP5K1B (K359N, K360N) and EGFP-PIP5K1C (K407N, K408N), respectively. For stable lines in MDA-MB-231 cells, stable lines expressing GFP-PIP5K1B (pCW57.1, Addgene #71782) or GFP-PIP5K1C (pPB, Addgene #104454) were made under a doxycycline-inducible promoter.

### Transfections, nucleofections, and lentiviral transduction

*Dictyostelium* AX2 cells were transfected using a standard electroporation protocol. Briefly, for each transfection, 5 × 10^6^ cells were pelleted, washed twice with ice-cold H-50 buffer (20 mM HEPES, 50 mM KCl, 10 mM NaCl, 1 mM MgSO4, 5 mM NaHCO3, 1 mM NaH2PO4, pH adjusted to 7.0) and subsequently resuspended in 100 μl ice-cold H-50 buffer. 2 μg of total DNA was mixed with the cell suspension, which was then transferred to an ice-cold 0.1-cm-gap cuvette (Bio-Rad, 1652089) for two rounds of electroporation at 850 V and 25 μF with an interval of 5 s (Bio-Rad Gene Pulser Xcell Electroporation Systems). After a 10 min incubation on ice, the electroporated cells were transferred to a 10-cm Petri dish containing HL-5 medium. The cells were selected by the addition of hygromycin B (50 μg ml−1) and/or G418 (20–30 μg ml−1) after 24h as per the antibiotic resistance of the vectors.

RAW 264.7 cells and MDA-MB-231 cells were transiently transfected by nucleofection in an Amaxa Nucleofector II device using Amaxa Cell line kit V (Lonza, VACA-1003) by slightly modifying pre-existing protocols ^121^. For each transfection, 3 × 10^6^ cells were pelleted, washed, and resuspended in 100 μl supplemented Nucleofector Solution V. A total of 2-3 μg of DNA mixture was added and immediately transferred to a Lonza cuvette for electroporation using the program settings D-032 and X-013 for RAW 264.7 cells and MDA-MB-231 cells, respectively. Immediately, pre-warmed pH-adjusted culture medium (500 μl) was added to the electroporated cells in the cuvette. The cell suspension was then transferred to a 1.5 ml vial and incubated at 37 °C and 5% CO2 for 10 min and 30min for RAW 264.7 cells and MDA-MB-231 cells, respectively. Next, 50– 100 μl solution containing cells was transferred to a coverslip chamber and allowed to adhere for 1 h. Finally, approximately 400 μl of pre-warmed pH-adjusted culture medium was added to each chamber. RAW 264.7 cells were further incubated for 4–6 h before imaging. MDA-MB-231 cells were imaged the next day.

Stable expression lines in HL-60 cells or MDA-MB-231 cells were generated by a combination of 3^rd^ generation lentiviral- and PiggyBac^TM^ transposon-integration based approaches ^73,120^. For transposon integration in HL-60 or MDA-MB-231 cell line, 5 μg transposon plasmid was co-electroporated with an equal amount of transposase expression plasmid into two million cells using Neon^TM^ transfection kit (Invitrogen; MPK10025B). Cells and DNA mix were resuspended in buffer ‘R’ before electroporation in 100 μl pipettes at 1350V for 35 ms in Neon^TM^ electroporation system (Invitrogen; MPK5000). Cells were resuspended in mixed culture medium in a 6-well plate and allowed to recover for 24 hours. Post-recovery, transfected cells were selected in presence of 10 mg/mL blasticidine S or 2.5 μg /mL Puromycin for 5-6 days. Once blasticidine S was removed, resistant cells were transferred to 48-well cell culture plate (Sarstedt; 83.3923), and further grown over 3-4 weeks into stable cell lines. Stable cells were maintained throughout in Blasticidine S or Puromycin.

For lentivirus transfection, 2^nd^ generation virus was prepared in HEK293T cells grown to about 80% confluency in 10 cm cell culture dish (Greiner Bio-One; 664160). For each reaction, a mixture of 2 μg pMD2.G (Addgene #12259), 8 μg psPAX2 (Addgene #12260), and 10 μg expression constructs in transfer plasmids were transfected using Lipofectamine 3000 as per manufacturer’s instructions (Invitrogen; L3000-008). After 96 hours, virus containing culture medium was harvested at 3000 rpm for 20 mins at 4°C. In a 6-well plate (Greiner Bio-One; 657160), entire viral medium was added to 4 x 10^6^ HL- 60 cells (seeded at a density of 0.25 x 10^6^ cells/mL) in presence of 10 μg /mL polybrene (Sigma; TR1003). After 24 hours, viral medium was removed, and cells were introduced to a mix of fresh and conditioned (mixed) culture medium. For selecting high expressors, infected cells were sorted after 5 days, and subsequently, grown to 80% confluency. Briefly, cells were pelleted, resuspended in sorting buffer (1x PBS, Ca2+/Mg2+ free; 0.9% heat-inactivated FBS; 2% penicillin-streptomycin) at a density of 15 x 10^6^ cells/ml, and sorted using 100 mm microfluidic sorting chip. High expressors (top 1%-5%) were collected in fresh culture medium (containing 2% penicillin-streptomycin) and grown to confluency.

### Retroviral production for primary T cell transduction

To produce retroviral particles for overexpression of PIP5K1C, 293T cells (originally obtained from ATCC) were grown to 70% confluency in 6-well plates before being co-transfected with 4.5µg nNeon-PIP5K1C (or mNeon alone as a control, in the MSCV backbone, VectorBuilder) along with 3.5µg pCL-eco packaging vector (Addgene #12371). Transfection was done using calcium phosphate purchased from Invitrogen (cat# K278001). Transfection mixes remained on 293T cells overnight before being replaced with fresh media, after which cells were incubated for an additional 24-30hrs. Retroviral containing supernatants were collected and centrifuged for 10min at 1000g to remove cellular debris, and polybrene was added at a final concentration of 8µg/mL. Retroviral supernatants were then immediately applied to T cells for transductions.

To generate lifeact-mScarlet retroviral particles, 4.5µg lifeact-mScarlet (MSCV backbone, VectorBuilder) and 3.5µg pCL-eco packaging vector were co-transfected into 293T cells and harvested as above.

### Primary CD8+ T cell isolation and transduction

Primary CD8+ T cells were isolated from spleen and lymph nodes of WT B6 mice (C57BL/6J, strain #000664, 8-12 weeks old). Mice were purchased from The Jackson Laboratory and housed at Upstate Medical University mouse facility in accordance with the Institutional Animal Care and Use Committee guidelines. Briefly, spleens and lymph nodes were homogenized and filtered before incubation with ACK lysis buffer (77mM NH_4_Cl, 5mM KHCO_3_, and 55µM EDTA pH 7.2-7.4) for 1 min to remove red blood cells (RBC). After RBC lysis, cells were washed and incubated with anti-CD4^+^ (clone GK1.5) and anti-MHCII (clone M5/114) for 20min at 4°C. followed by magnetic separation using BioMag Goat anti-rat IgG beads (Qiagen cat#310107). Isolated CD8^+^ T cells were then immediately activated on a 24-well plate coated with 5µm/mL anti-CD3 (clone 2C-11) and 2µm/mL anti-CD28 (clone PV-1) at 1 X 10^6^ cells per well. Activation was done in T cell media composed of Corning DMEM supplemented with 5% FBS, 1% each penicillin/streptomycin, non-essential amino acids, and Glutamax (all purchased from Gibco), plus 2µL 2-mercaptoethanol (obtained from Sigma Aldrich). 24hrs later, the T cell media was collected and put aside for cell culture post-transduction. 800uL of viral supernatants were gently added per well and plates were incubated at 37°C for 10min. Plates were centrifuged at 1100*g* for 2hrs at 32°C. Plates were then incubated at 37°C for 10min and then viral supernatants were removed and replaced with 1.5mL of pre-collected conditioned media mixed with fresh complete T cell medium at a 2:1 volume ratio. T cells were cultured for an additional 24hrs before being removed from activation and mixed at a 1:1 volume ratio with fresh T cell media containing recombinant 80 U/mL IL-2 to create a final concentration of 40U/mL. T cells were used on days 5-6 post-harvest for all experiments.

To generate lifeact-mScarlet expressing T cells, CD8^+^ T cells were transduced as above with 200uL of lifeact-mScarlet viral supernatants along with 600uL of mNeon-PIP5K1C or mNeon alone.

### Microscopy

All *Dictyostelium* experiments were performed on a 22 °C stage. All RAW 264.7, MDA-MB-231 and HL-60 experiments were performed inside a 37 °C heated chamber with 5% CO2 supply. All time-lapse live-cell imaging experiments were performed using one of the following microscopes: (1) Zeiss LSM 780-FCS Single-point, laser scanning confocal microscope (Zeiss Axio Observer with 780-Quasar; 34-channel spectral, high-sensitivity gallium arsenide phosphide detectors), (2) Zeiss LSM800 GaAsP Single-point, laser scanning confocal microscope with wide-field camera, (3) Nikon Eclipse Ti-E dSTROM total internal reflection fluorescence (TIRF) microscope (Photometrics Evolve EMCCD camera). The Zeiss 780 confocal microscope was controlled using ZEN Black software; Zeiss 800 confocal microscope was controlled using ZEN Blue software; the Nikon TIRF was controlled using Nikon NIS-Element software. The 40X/1.30 or 63X/1.4 Plan-Neofluar oil objective (with proper digital zoom) was used in all microscopes. The 488 nm (Ar laser) excitation was used for GFP and YFP, 561 nm (solid-state) excitation was used for RFP and mCherry, and 633nm (diode laser) excitation was used for miRFP703 in Zeiss 780 and 800 confocal microscopes. The 488nm (Ar laser) excitation was used for GFP and 561 nm (0.5W fiber laser) excitation was used for mCherry and RFP in Nikon TIRF.

### Cell preparation for live-cell imaging

Before starting the image acquisition, growth phase vegetative *Dictyostelium* (unless otherwise mentioned) were allowed to adhere on an eight-well coverslip chamber (Nunc Lab-Tek) for ∼15 min and then HL-5 media was aspirated and replaced with 400-500 μL DB media (5 mM Na2HPO4 and 5 mM KH2PO4 supplemented with 2 mM MgSO4 and 0.2 mM CaCl2) Cells were incubated in DB for ∼40-60 min before starting imaging experiments. For details on the preparation of developed or electrofused *Dictyostelium* cells for imaging, please see “*Cell differentiation”* and “Electrofusion” section below.

Differentiated HL-60 macrophages were allowed to adhere on similar surface for ∼40 min. Differentiated HL-60 neutrophils, pre-treated with heat-killed *Klebsiella aerogenes*, were allowed to adhere on fibronectin-coated chambers for ∼20 min. Next, for HL-60 neutrophils or macrophages, the culture media was replaced with fresh RPMI 1640 medium and adhered cells in fresh media were used for imaging. MDA-MB-231 cells were introduced to 8-well chamber the ∼12 hour before the experiment. Spent culture media was replaced with fresh culture media before the experiments. For more details on wound healing and 3D spheroid assays with MDA-MB-231 cells, please see the relevant sections below. Generally, after nucleofection and adhesion to the substrate (see above), RAW 264.7 cells were imaged in Hank’s balanced salt solution (HBSS buffer) supplemented with 1 g/L glucose. For more details on RAW 264.7 cell preparation for microscopy, please see relevant sections below.

To induce PIP5K expression, doxycycline (50 µg ml^−1^ for Dictyostelium cells, 1 µg ml^−1^ for MDA-MB-231 cells) was added 12 h before imaging. For macropinocytosis assay, *Dictyostelium* cells were incubated with 2 mg/ml TRITC-dextran or FITC-dextran for 10 min, washed twice with DB and imaged subsequently. All live- or fix-cell imaging was acquired with 0.3–0.8% (*Dictyostelium*) or 1–5% (HL-60) laser intensity. In single cells, confocal imaging was performed at a middle z-plane of the cells. Images were aquired at a frequency of 5-60s/frame as appropriate. To visualize cortical/ventral waves in *Dictyostelium* and HL-60 macrophage cells, confocal microscopes were focused on the very bottom of the cell to capture the substrate-attached ventral surface of the cell.

For the cytoskeletal dynamics inhibition experiments, neutrophils were treated with 10 µM Y-27632 for at least 10 min before imaging and 8 µM jasplakinolide and 5 µM latrunculin B were added during imaging. In *Dictyostelium,* for myosin II or branched actin inhibition experiments, 50 µM blebbistatin or 100 µM CK666 was added during imaging, respectively. For arresting actin polymerization in *Dictyostelium* cells, 5 µM latrunculin A and 4 mM caffeine were added at least 20 min before imaging (note that, caffeine can induce increased wave propagation in cells ^6,7,18^.

To image primary CD8+ T cells, Ibidi 8-well, glass-bottomed chamber slides were coated with ICAM-1 at 2µg/mL (Bio-Techne) overnight at 4°C. Activated, transduced CD8^+^ T cells were washed and resuspended in Leibovitz’s L-15 medium (Gibco) containing 2mg/mL D-glucose and 5 x 10^4^ cells were added to each well. After 20min, wells were gently washed to remove non-adherent cells and debris and mounted on a Marianas system (3i) consisting of a Ziess Axio Observer 7 equipped with a X-Cite mini+ light source (Excelitas) and a Prime BSI Express CMOS camera (Photometrics) enclosed in an environmental chamber (Okolab). For cell tracking, movies were collected every 30 seconds for 10min using a 20x objective at 37°C with SlideBook software (3i) and exported to ImageJ for analysis. mNeon expressing cells were tracked using the manual tracking plugin to generate XY coordinates for each time point. We only included cells that stayed in the field-of-view for the entire imaging period. 20x movies were also obtained with higher time resolution (2 seconds per timepoint). 63x imaging was performed every 3 seconds for 1 min total.

### Electrofusion

*Dictyostelium* cells were elecrofused to form giant cells using an established protocol ^17,47^. First, the growth phase cells were collected from suspension culture, washed twice, and finally resuspended in 10 ml SB buffer (17 mM Soerensen buffer containing 15 mM KH_2_PO_4_ and 2 mM Na_2_HPO_4_, pH 6.0) at a density of 1.5 × 10^7^ cells/mL. Next, to facilitate the formation of cell clusters, the cells were placed in a conical tube and gently rolled for 30 to 40 minutes. Following this, 800 μL of the rolled cells were moved to a Bio-Rad cuvette with a 0.4 cm gap. This transfer was carried out using pipette tips with their edges cut off to ensure the cell clusters remained intact. To induce hemifusion, electroporation was conducted using a BioRad Gene Pulser (Model 1652098). The procedure involved one pulse at 1kV and 3 μF, followed by two more pulses at 1kV and 1 μF. A time interval of 3 seconds was maintained between each pulse. After electroporation, approximately 35 μL of the electrofused cells were extracted from the cuvette and placed into a Nunc Lab-Tek 8-well chamber. The cells were incubated for 5 minutes, after which 450 μL of SB buffer supplemented with 2 mM CaCl2 and 2 mM MgCl2 was added. Finally, the cells were left for about 1 hour to recover, settle, and adhere to the substrate before the image acquisition was initiated.

### Frustrated phagocytosis and osmotic shock

To visualize ventral waves in HL-60 macrophages that had been differentiated with PMA, we made minor adjustments to an existing protocol ^18,52^. First, the Nunc Lab-Tek eight-well coverslip chambers were prepared. This involved an initial pre-wash with 30% nitric acid, followed by a 3-hour coating with 1 mg ml⁻¹ BSA. The chambers were then washed with PBS and incubated for 2 hours with 5 μg ml⁻¹ anti-BSA (a 1:200 dilution). A final double wash with PBS was performed to remove any excess antibodies. Before imaging, the transfected and PMA-differentiated macrophage cells were starved for 30 minutes in a suspension of 1×Ringer’s buffer. This buffer consisted of 150 mM NaCl, 5 mM KCl, 1 mM CaCl2, 1 mM MgCl2, 20 mM HEPES, and 2 g l⁻¹ glucose, with a pH of 7.4. Finally, these starved cells were transferred to the prepared, opsonized chambers. The cells were allowed to spread on the antibody-coated surface for 5 to 10 minutes. Just before the imaging, a hypotonic shock was induced by applying a 0.5×Ringer’s solution.

### Optogenetics

Optogenetics perturbationexperiments were performed in *Dictyostelium*, RAW 264.7, or MDA-MB-231 cells using slightly modified protocols ^17,120,122^. To test the necessity and sufficiency of plasma membrane PIP5K in regulating symmetry breaking and protrusion formation, Opto-SeqP was introduced in RAW 264.7 macrophages and Opto-RecP was introduced in RAW 264.7 macrophages as well as MDA-MB-231 cells, respectively, using nucleofection (please see “*DNA constructs*” and “*Transfections, nucleofections, and lentiviral transduction*” sections for more details). In parallel, Opto-CTRL system was also introduced in these cell lines as a control for optogenetic phenotypes. Note that, cytosolic, functional version of PIP5K1C had been recently reported ^82^ which we used to engineer Opto-RecP. In all cases, 561 nm excitation laser was used to track the localization dynamics of mCrimson-SspB or mCherry-SspB tagged recruitee and 488 nm excitation laser was used to trigger the recruitment into specific subcellular location. The 633 nm excitation was used to record the Lifeact-miRFP703 dynamics throughout the experiment. Before turning 488 nm laser ON, the basal activities was recorded for 5-10 min, whereas after turning 488 nm laser ON, recruitment induced activities were recorded for another 30-40 min. For Opto-Rec system, the RAW 264.7 cells were globally stimulated with saturating dose of C5aR agonist immediately after tagged PIP5K was localized to plasma membrane by turning ON 488 nm laser; the MDA-MB-231 cells were maintained in saturating dose of EGF throughout the experiment. For Opto-SeqP recruitment, cells were not stimulated and maintained in standard HBSS media.

### Chemically induced dimerization and targeted screening through RasGEFs

To identify which RasGEFX mediate lowered PI(4,5)P2 driven Ras activation, we screened across the *Dictyostelium* RasGEF space which has 25 members (RasGEFA to RasGEFY). First, we analyzed the data from Iwamoto et al. ^19^, where all RasGEFs were hierarchical clustered based on their ability to induce spontaneous Ras activity when overexpressed. We selected all four members of “Cluster 1” from there and decided to trigger acute PI(4,5)P2 lowering in cells where these RasGEFs are knocked out. To this end, RasGEFB, RasGEFM, RasGEFU, and RasGEFX were knocked out from wild-type AX2 Dictyostelium cells at the Ueda lab, as described previously ^19^ and following plasmids were stably expressed in the knockout and control wild-type cells: a) cAR1-FKBP-FKBP (pDM358) as the membrane anchor and b) mCherry-FRB-Inp54p (pCV5) as recruitable enzyme for rapidly hydrolyzing PI(4,5)P2. Note that, this Inp54p-based chemically inducible dimerization system is a well-established synthetic PI(4,5)P2 lowering tool ^8,9,123^. Finally, on the day of imaging experiment, engineered *Dictyostelium* cells were transferred to an eight-well coverslip chamber and incubated for 10–15 min to allow adhesion. The HL-5 medium was then replaced with 450 μl DB buffer and cells were incubated for 15-20 min before starting live-cell imaging. After imaging a certain number of frames, rapamycin was added gently to the chamber (to a final concentration of 5 μM) during image acquisition to facilitate the recruitment of Inp54p to the plasma membrane and imaging was continued. Cell migration speed (before and after recruitment) and percent of fans and oscillators were quantified using a custom-written Python based workflow. Please see Image Analysis section and GitHub repo for further details.

To facilitate myosin II disassembly from the membrane-cortex in PIP5K overexpression cells (Figure S28), an analogues rapamycin inducible dimerization system was used that was previously developed at the Devreotes Lab by incorporating MHCKC as recruitee ^76^. To generate Figure S7D, mCherry–FRB-PIP5K_1-315aa_ (pCV5) was used as recruitee. In both cases, cAR1-FKBP-FKBP (pDM358) was used as membrane anchor.

### 2D wound healing assay

The wound healing assay was performed using Ibidi Culture-Insert 4 Well in μ-dish^35mm, high^ (Ibidi, catalog # 80466). 6 x 10^5^ MDA-MB-231 cells expressing doxycycline-inducible PIP5K1C were seeded in each insert well and incubated for 24 hours until a full monolayer was created. Next day, 1 µg/ml Dox was added to each monolayer to induce PIP5K1C expression. After another 16 hours Dox incubation, the 4-well insert was removed, creating a 500 μm cell-free area between cell monolayers. Cells were imaged using Zesis 780 Confocal microscopy with 20x objective, 5 x 5 tile function. The open scratch area was quantified at 1h, 15h, and 24h time points.

### 3D Spheroid assay

MDA-MB-231 cells expressing doxycycline-inducible PIP5K1C were trypsinized and resuspended at a concentration of 1 × 10^^5^/ml in normal media (10% FBS, DMEM, 2 mM glutamine). Spheroids were formed by incubating the cells in 20 μl hanging drop onto the underside of a 10-cm culture dish for 72 hours. At 72h, 2 µg/ml Dox was added to each drop to induce PIP5K1C expression. Next day, spheroids were collected in a tube previously coated with 2.5% (vol/vol) BSA solution in DPBS. After centrifugation at 400 xg, 5 min, spheroids were resuspended in growth-factor-reduced Matrigel (Corning, 354230) as previously described. Spheroids were embedded at a density of 1 spheroid per μl into the Matrigel solution and 50 μl of suspension per well was plated into an 8-well glass-bottomed chamber on a 37°C heating block. After 1-hour incubation at 37°C, 0.5 ml of normal media was added to the wells. Spheroids were disseminated into Matrigel over a period of 8h to 3 days. Spheroid images were acquired by a Zesis Confocal LSM 780 microscope with a 40x water objective. Dissemination of spheroids was quantified using ImageJ to outline the core of the spheroid and the total spheroid area (dissemination strand and core).

### Cell differentiation

For *Dictyostelium* cell development, 1 × 10^8^ cells in the exponential growth phase were collected from suspension culture and pelleted as previously described ^124,125^. After washing with DB twice, the cells were resuspended in 4 ml DB and shaken at 105 r.p.m. for 1 h. The cells were then pulsed with 100 nM cAMP (5s pulse every 6 min) using a time-controlled peristaltic pump for 5-6 h with continual shaking. This allowed the cells to develop and polarize. After development, Around 10–15 μL of differentiated cells (in DB media) was collected were transferred from the shaker to an eight-well coverslip chamber, resuspended thoroughly in 450 μl DB and incubated for 20–30 min before imaging. For development on non-nutrient agar, cells were washed with developmental buffer (DB) and plated on solidified DB agar at a density of 5.2 × 10^5^ cells/cm^2^.

For development on non-nutrient agar, cells were washed with developmental buffer (DB) and plated on solidified DB agar at a density of 5.2 × 10^5^ cells/cm^2^. For most experiments, development in suspension was carried out by pulsing cells resuspended in DB at 2 × 10^7^ cells/ml with 50 to 100 nM cAMP every 6 min starting from 1 hour.

### Global receptor activation assay

For cAMP receptor (cAR1) activation experiments, *Dictyostelium* cells were first differentiated in suspension as described in “Cell differentiation” section. Cells were subsequently incubated with 5 μM Latrunculin A for around 20 min before imaging. Using a confocal laser scanning microscope, 10-20 frames were first acquired to record the basal activity of the proteins, then cAMP was added to the chamber (to a final concentration of 1 μM) to activate all the cAR1 receptors, and the image acquisition was continued. An imaging frequency of 3 s/frame was maintained throughout the experiment.

For folate receptor (fAR1) activation experiments, vegetative growth phase *Dictyostelium* cells, cells were seeded on the eight-well coverslip chamber and incubated for 20–30 min. Then, the HL5 medium was replaced with DB and incubated cells with an additional 20 minutes before the imaging. Cells were subsequently incubated with 5 μM Latrunculin A for around 20 min before imaging. Using a confocal laser scanning microscope, 10-20 frames were first acquired to record the basal activity of the proteins, then folic acid was added to the chamber (to a final concentration of 0.1, 1, 10, or 100 nM) to activate all the fAR1 receptors, and the image acquisition was continued. An imaging frequency of 3 s/frame was maintained throughout the experiment.

For RAW 264.7 cells, cells, with either PH_Akt_-mCherry only or with PIP5K-GFP and PH_Akt_-mCherry coexpression, were collected after 4-6 hours of nucleofection (as described in “*Transfections, nucleofections, and lentiviral transduction*” section). The culture medium was then replaced with Hank’s balanced salt solution (HBSS buffer) supplemented with 1 g/L glucose and cells were incubated for another 30 min before the imaging. Using a confocal laser scanning microscope, 10-20 frames were first acquired to record the basal activity of the proteins, then C5aR agonist was added to the chamber (to a final concentration of 0.1, 1, 10, or 100 μM) to activate the C5a receptors, and the image acquisition was continued. An imaging frequency of 15 s/frame was maintained throughout the experiment.

### 2D Chemotaxis Assay

We slightly modified a pre-existing protocol to perform chemotaxis assay for vegetative *Dictyostelium* cells ^126^. Briefly, cells were seeded on the 1-well Nunc Lab-Tek chamber as a drop at the center and incubated for 20–30 min. Then 2 mL DB buffer was added to the chamber and cells were dispersed by pipetting multiple times. A Femtotip microinjection needle (Eppendorf) was loaded with 10 μM of filtered folic acid solution and then connected to a FemtoJet microinjector (Eppendorf). The microinjector was operated in continuous injection mode with a compensation pressure of 15 hPa. To initiate the chemotaxis, the micropipette was brought to the (x,y,z) coordinate of cells using a programmed micromanipulator. The imaging was continued with an acquisition frequency of 30 s/frame.

### Multinucleation assay

Cells that have been in suspension at 200 rpm for 2–3 days, or cultured on adherent surface, were seeded at ∼70% confluency on glass coverslips in HL-5 medium and allowed to adhere for 20 min. Next, HL-5 medium was replaced with ice-cold fixative solution (2% PFA + 0.01% Triton X-100 in 1.5× HL-5 medium) for 10 min. Cells were then washed quickly with 1× PBS, and the nuclei were stained with 10 µg/ml Hoechst for 10 min. Afterwards, cells were washed three times, 10 min each time, with 1× PBT (1× PBS + 0.01% Triton X-100). Before visualization, cells were mounted on coverslips using Invitrogen SlowFade^TM^ Gold antifade reagent. Cells were imaged using Zeiss LSM 780-FCS Single-point, laser scanning confocal microscope with 405 nm laser.

### Immunofluorescence

*Dictyostelium* cells were seeded on glass coverslips in HL-5 medium and allowed to adhere for 20 min. Next, HL-5 medium was replaced with fixative solution (2% buffered paraformaldehyde with 0.25% Glutaraldehyde and 0.1% Triton X-100 in HL-5 medium) for 10min at room temperature (RT), then quenched in 1mg/mL Sodium Borohydride for 10min, then washed three times with 1x PBT (1x PBD supplemented with 0.5% BSA and 0.05% Triton X-100), each time for 3min. Cells were then blocked in blocking buffer (3% BSA in 1× PBT) for 1 h at room temperature. Blocking buffer was removed and cells were washed with 1x PBT quicky then incubated with Invitrogen Alexa Fluor ^TM^ 647 phalloidin (1: 200 dilution) for 1h at RT. Cells were then washed three times with 1x PBT, each time for 3min. Before visualization, cells were mounted on coverslips using Invitrogen SlowFade^TM^ Gold antifade reagent. Cells were imaged using Zeiss LSM 780-FCS Single-point, laser scanning confocal microscope with 405 nm laser.

Migrating T cells were fixed with 3.7% paraformaldehyde (Electron Microscopy Sciences) and blocked and permeabilized for 15mins at room temp (1% BSA and 0.15% Triton X-100 in PBS). Cells were then incubated with fluorescent phalloidin (alexa 568) and DAPI for 45mins at room temp. Cells were then washed three times in block/perm solution, and put in PBS for imaging. 0.5µm *z*-stacks for a total of 8um were collected at 63x from the coverglass moving up through the cell. Each z-slice was background subtracted and the image was projected.

### Cell preparation for single-molecule imaging

Cells were cultured in HL5 medium in the presence of 50 μg/mL Doxycycline for 8 hours at 21°C. During the final 1 hour, the medium was supplemented with HaloTag® TMR ligand (Promega) at the final concentration of 40 nM. The cells were washed with DB buffer and resuspended in DB buffer at a cell density of 3 × 10^6^ cells/mL. A 10 μL cell suspension was placed on a coverslip (25 mm diameter, 0.12–0.17 mm thick; Matsunami) that was washed by sonication in 0.1 N KOH for 30 min and rinsed with 100% ethanol prior to use. After a 10 min incubation, the cells were overlaid with an agarose sheet (8 mm × 8 mm, Agarose II; Dojindo). After 20 min of incubation, the coverslip was set in an Attofluor^TM^ cell chamber (Invitrogen) and observed by TIRF Microscope.

### Microscopy setup for single-molecule imaging

The methods for single-molecule imaging are described elsewhere [Banerjee et al., 2023]. In brief, objective-type TIRFM was constructed on an inverted fluorescence microscope (Ti; Nikon) equipped with two EM-CCD cameras (iXon3; Andor), solid-state CW lasers (OBIS 488-150 LS and OBIS 561-150 LS; Coherent), and an objective lens (CFI Apo TIRF 60X Oil, N.A. 1.49; Nikon). The PIP5K-Halo and RBD-GFP images were acquired at 30 and 1 frames/s, respectively, with a software (iQ2; Andor).

### Statistical analysis of single-molecule trajectories

The fluorescent images of single molecules were tracked with a laboratory-made software to yield the *x*- and *y*-coordinates of each molecule. The methods for estimating lateral diffusion coefficient and membrane-binding lifetime are described elsewhere ^18,88^. Briefly, diffusion coefficient was estimated by fitting the statistical distribution of the displacement, Δ*r*, that a single molecule moved during a time interval, Δ*t* = 33.3 ms, to the following probability density function,

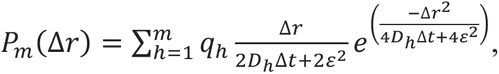

where *D_h_*, *q_h_*, *m*, and *e* denote the diffusion coefficient and fraction of the *h*-th mobility state, total number of mobility states, and standard deviation of the measurement error, respectively. Short-range diffusion coefficient, *D_SRD_*, of an excerpt of a trajectory for 0.5 sec was estimated by fitting the mean square displacement (MSD) with the linear function,

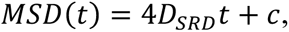

where *c* is a constant representing the measurement error. The membrane-binding lifetime was quantified based on the statistical distribution of the time a single molecule was detected on the membrane. The distribution in cumulative form was fitted to an exponential function as,

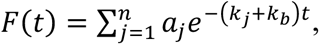

where *k_j_*, *a_j_*, *n*, and *k_b_* denote the decaying rate constant of the *j*-th binding state, the fraction of the *j*-th binding state, the total number of binding states, and the rate constant of photobleaching of TMR, respectively. The fraction of the molecules that adopt the *j*-th state at an arbitrary time point was calculated as,

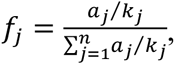

whereas a mean of the membrane binding lifetime was calculated as,

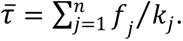

### Reaction-diffusion model

#### Core excitable network

The simulations are based on a model whose core consists of three interacting species–Ras, PIP2 (PI(4,5)P2), and PKB (PKBA)–which together form an excitable network ^8^. The concentrations of these species are governed by stochastic reaction-diffusion partial differential equations (RD-PDEs):

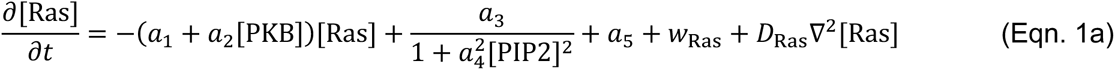

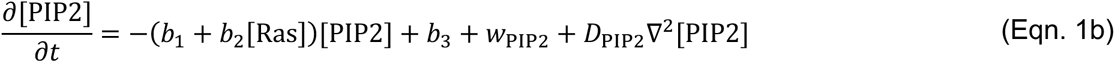

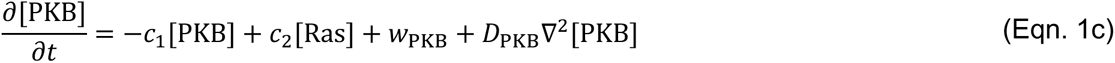

In each equation, the final term represents the diffusion of the species, where 𝐷_∗_denotes the diffusion coefficient of the respective species, and ∇^2^ is the spatial Laplacian (in one dimension). The penultimate term 𝑤_∗_ accounts for intrinsic molecular noise. Noise is modeled using a Langevin approximation, with its magnitude determined by the corresponding reaction terms (Gillespie, 2000)^127^. For example, the noise term for PKB is given by:

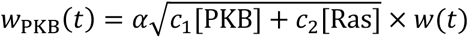

Here, 𝑤(𝑡) is Gaussian white noise process with zero mean and unit variance. The parameter 𝛼 is an empirical scaling factor used to adjust the magnitude of the noise in simulations.

#### PIP5K Dynamics via Ras-Regulated State Transitions

We assume that PIP5K exists in two distinct states: a membrane-bound fraction (PIP5K_mem_) and a cytosolic fraction (PIP5K_cyto_), characterized by low and high diffusivity, respectively. The total amount of PIP5K is assumed to be conserved within the cell, though its local membrane distribution may vary over space and time. The interconversion between membrane-bound and cytosolic states is regulated by Ras activity, which promotes the transition from PIP5K_mem_ to PIP5K_cyto_. The spatiotemporal dynamics of PIP5K are governed by the following coupled RD-PDEs:

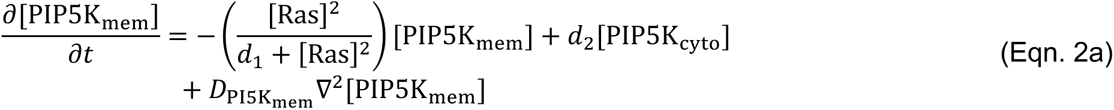

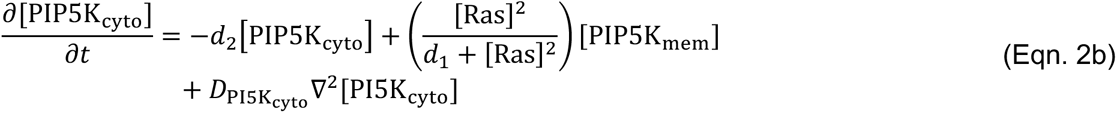

We do not include noise terms in these equations. The conservation of total PIP5K concentration follows from the fact that the sum of the reaction terms in the Eqn. (2a) and (2b) is zero. Consequently, changes in local concentration arise solely from diffusion. Integrating over the complete spatial 1D domain and using the fact that boundary conditions are periodic guarantees that the total concentration of PIP5K remains constant. Functionally, the main role of PIP5K in the model is to enhance the production rate of PIP2, implemented as a modulation of the parameter 𝑏_1_ via:

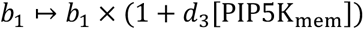

#### Cytoskeleton feedback loops

Finally, we incorporate three other terms. The first represents local positive feedback from branched actin onto Ras. Since the actin cytoskeleton is not directly modeled, we model the origin of this feedback from PKB, which is upstream to the cytoskeleton ^8^, and incorporated the actin dynamics as:

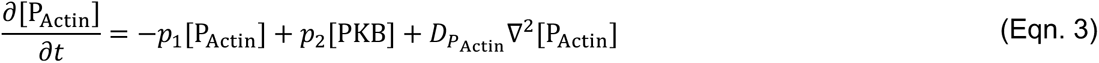

The second feedback loop accounts for the local inhibitory regulation by actomyosin on Ras that originates from PIP2:

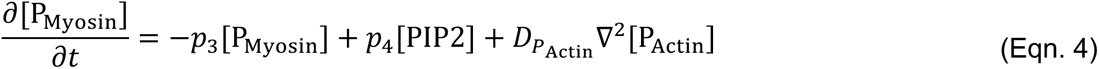

The third feedback loop models global inhibition from membrane tension ^5,30^. Membrane tension is treated as a spatially uniform variable influenced by average PKB activity:

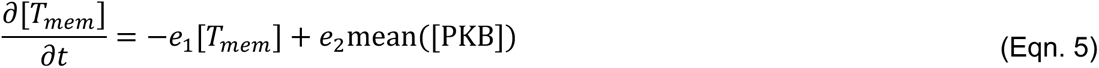

Together, these feedbacks modify the Ras dynamics by altering the reaction terms in Eqn. (1a) as follows:

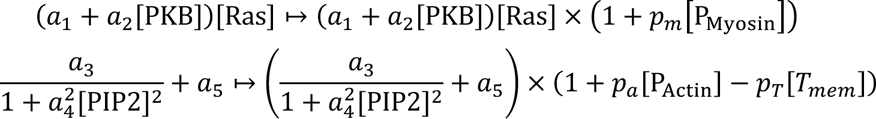

#### Modeling perturbations

The perturbations in Fig. 7, Supplementary Fig. S35, S36 were done by modifying the reaction terms associated with the relevant species during the simulation. Specifically:

|  |  |  |
| --- | --- | --- |
| over expressing PIP5K | PIP5K_total = 0.1508 | The total amount of PIP5K was increased approximately 1.9-fold compared to the wild type. |
| piki- fan phenotype | PIP5K_total = 0.0397,<br>$p_a = 1.25$<br>$p_T = 0.75$ | Compared to wild type:<br>- PIP5K total reduced by ~50%<br>- Ras activation by Actin increased 4.5-fold<br>- Membrane tension-mediated Ras inhibition reduced by 20% |
| piki- oscillator phenotype | PIP5K_total = 0.0016,<br>$p_a = 0.1250$<br>$p_m = 0.0750$<br>$p_T = 0.75$ | Compared to wild type:<br>- PIP5K total greatly reduced (~2% of wild type)<br>- Ras activation by Actin reduced 2-fold<br>- Ras inhibition by Myosin reduced 2-fold<br>- Membrane tension-mediated Ras inhibition reduced by 20% |
| Latrunculin treated | $p_1 = 0$<br>$p_3 = 0$ | Actin and Myosin were depleted by blocking their production via PKB and PIP2, respectively. |

a. Eliminating PIP5K: 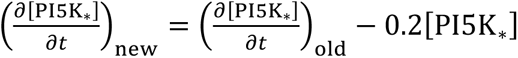, for both types of PIP5K
b. Increasing PIP5K: 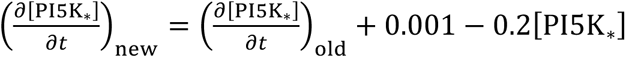, for both types of PIP5K.
c. Eliminating myosin: 𝑝_4_ = 0 and 𝑝_𝑚_ = 0.
d. Eliminating actin: 𝑝_2_ = 0 and 𝑝_𝑎_ = 0.

### Level Set-Based Modeling of Cell Shape Evolution

To evaluate the impact of molecular perturbations on cell morphology, we simulated cell behavior using level set methods (LSM), based on the framework as previously described _8,92_. In this approach, the cell boundary is represented as the zero-level set of a potential function 𝜑(𝑥, 𝑡), 𝑥 ∈ ℝ^2^. The evolution of this potential function over time is governed by:

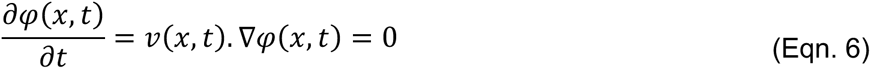

Here, 𝑣(𝑥, 𝑡)denotes the local velocity field derived from mechanical responses of the cell to applied stresses. To compute this velocity, we employed a viscoelastic model, which accounts for the deformation of both the membrane and the underlying cell cortex:

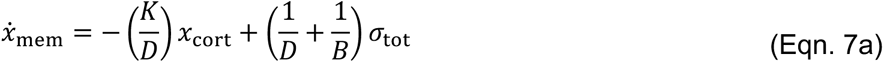

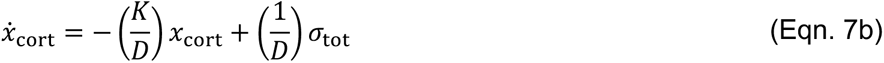

In these equations, 𝜎_tot_ represents the total stress applied to the cell boundary, while 𝑥_mem_ and 𝑥_cort_ are the displacements of the membrane and cortex, respectively. The parameters in Eqn. (7) denote various viscoelastic attributes of the cell: the elasticity (𝐾) and viscosity (𝐷) of the membrane, and the viscosity of the (𝐵) of the cytoplasm. The boundary velocity 𝑣 used in Eqn. (6) is obtained from 𝜕𝑥_mem_/ 𝜕𝑡.

The total stress 𝜎_tot_ includes contributions from:

1. **Cortical tension**: 𝜎_ten_ = 𝛾𝜅(𝑥), where 𝛾is the local cortical tension, and 𝜅(𝑥) is the local membrane curvature.
2. **Volume conservation**: 𝜎_vol_ = 𝑘_area_(𝐴(𝑡) − 𝐴_0_), enforcing approximate conservation of the enclosed area by the cell boundary 𝐴(𝑡) relative to the initial rest area 𝐴_0_.
3. **Protrusive forces**: 𝜎_pro_ = 𝜎_0_𝑅(𝜃), where 𝑅(𝜃) is the scaled input from the reaction-diffusion model and 𝜎_0_is the conversion factor from molecular activity to pressure. Here, we used 𝑅(𝜃) = min(max(PKB(𝜃) − 0.3,0), 0.25). This formulation ensures that the protrusive forces are within the experimentally observed range of 0.5–5 nN/μm².

#### Implementing the simulations

Simulations were performed in MATLAB 2025a (MathWorks, Natick, MA) using custom code based on the Itô formulation implemented via the Stochastic Differential Equation Toolbox (http://sdetoolbox.sourceforge.net). The simulations aim to reproduce membrane fluorescence patterns observed in single-cell confocal images. To reflect the spatial scale of a typical cell, we assumed a cell with a radius of 5 µm and 314 grid points along its perimeter under periodic boundary conditions, yielding a spatial resolution of approximately 0.1 µm. Initial conditions were obtained by solving for the steady state of the full model, assuming that the dynamics of PIP2, Actin, Actomyosin, PIP5K, and membrane tension (T_mem_) are sufficiently slow to be in quasi-steady state. The temporal resolution was 1 ms. The total simulation time was 900 s, with kymograph results shown over a 300 s window following the exclusion of initial transients. Level set simulations were performed using the Level Set Toolbox for MATLAB with first-order forward Euler time integration. Parameters used in these simulations are provided in Supplementary Table 1.

### Image Analysis

All images were analysed with Fiji/ImageJ 1.52i (NIH) and/or Python v.3.10 (with numpy, scipy, pandas, scikit-image, opencv, pillow, along with standard Python libraries). We utilized Python v3.10 (with Seaborn and/or Matplotlib library), GraphPad Prism 10 (GraphPad), OriginPro v.9.0 (Originlab Corporation), LibreOffice Calc (The Document Foundation) and Microsoft Excel (Microsoft) for plotting our results.

#### Hierarchical classification of cellular morphologies

First, for each of the N cells, time-lapse image series were obtained from microscope raw data. Second, each of these image series were segmented with a custom-written segmentation pipeline that involved ROI selection (to segregate cells coming in contact with each other), thresholding/binarization, multiple morphological processing steps including dilation, erosion, fill hole, and despeckle. Third, each of these binarized image series were processed with a custom-written montaging pipeline that augmented different image series and provided a specified padding between each of these cells to ultimately result a single large time-series of images for further unbiased processing. In the final of preprocessing, a custom-written Jython script was used to generate and export “labelled” images (where each label corresponds to a particular cell across time-series) using Trackmate Nearest-Neighbor tracker in a headless, automated way. In the hierarchical classification workflow, for each cell across the entire time series, multiple morphological parameters are obtained using scikit-image measure.regionprops function and further secondary metrics were computed from there to finally obtain the values of areas, rolling coefficient of variation of areas, circularity, aspect ratio, angle of orientation, angle displacement, angle between the displacement vector and orientation vector (and its rolling coefficient of variation). Based on the kernel density estimation plots of these parameters, specific threshold values were decided for each, as shown in Figure S1 and S2 (please see figure legends there). We used a combination of these parameters to categorize cells into one of the five final categories (based on the modal value for each cell over time): Amoeboid, Spiky, Round, Oscillator, and Fan/Keratocyte like. For more details and complete code, please visit the GitHub repo: https://github.com/tatsatb/PIP5K-Ras-bistability-symmetry-breaking.

#### Kymographs

For the membrane kymographs, the cells were segmented against the background following standard image-processing steps with custom code written in Python v3.10 (along with numpy, scipy, pandas, scikit-image, opencv, pillow, tkinter, Seaborn, and Matplotlib library). The kymographs were created from the segmented cells as previously described, where consecutive lines over time were aligned by minimizing the sum of the Euclidean distances between the coordinates in two consecutive frames using a custom-written Python function. A linear colourmap was used for the normalized intensities in the kymographs. In colored kymographs, the lowest intensity is indicated by blue and the highest by yellow. Line kymographs that accompanied ventral waves were generated in Fiji/ImageJ by drawing a thick line with a line width of 8–12 pixels and processing the entire stack in the ‘KymographBuilder’ plugin.

#### t-stacks

The ImageJ (NIH) software was used for image processing and analysis. For t-stacks, TIRF time-lapse videos were opened and converted to greyscale in ImageJ. The ImageJ 3D Viewer plugin was then used to stack the frames from the video. The resampling factor is set to 1 to avoid blurring of activities between frames.

#### Line-scan intensity profile

Line scans were performed in Fiji/ImageJ (NIH) by drawing a straight-line segment (using line tool) inside the cells with a line width of 8–12 pixels to obtain an average intensity value. The intensity values along that particular line were obtained in the green and red channels using the ‘Plot Profile’ option. The values were saved and normalized in OriginPro 9.0 (OriginLab). The intensity profiles were graphed and finally smoothened using the adjacent-averaging method, using proper boundary conditions. The plots were then normalized by dividing by the maximum value. For a particular line scan, the green and red profiles were smoothed and processed using the exact same parameters to maintain consistency.

#### Membrane to cytosol ratio quantification

The cytosol of cells was first outlined with freehand selection tool in Fiji/ImageJ (NIH). The mean intensity of the cytosol was measured using analyze>>measure function. Then make band function was applied to the outline and proper band size was applied according to the cells. Then mean membrane intensity was measured using the same method. Using mean membrane to cytosol intensity to divide mean cytosol intensity, we can get the membrane to cytosol ratio for each biosensor. Each data was then plotted as box-and-whisker graphs in GraphPad Prism 10.

#### Cell migration analysis

Analysis was performed by segmenting *Dictyostelium* or neutrophil cells in Fiji/ImageJ 1.52i software. For this, image stack was thresholded using the ‘Threshold’ option. ‘Calculate threshold for each image’ box was unchecked, and range was not reset. Next, cell masks were created by size-based thresholding using the ‘Analyzed particles’ option. To optimize binarized masks, ‘Fill holes’, ‘Dilate’, and ‘Erode’ were done several times. For creating temporal color-coded cell outlines, ‘Outline’ was applied on binarized masks, followed by ‘Temporal-Color Code’ option. Next, ‘Centroid’ and ‘Shape descriptors’ boxes were checked in ‘Set Measurements’ option under ‘Analyze’ tab. This provided us with values for centroid coordinates and aspect ratio. Mean and SEM from replicates of aspect ratio values were determined and plotted in GraphPad Prism 10. The starting point for centroid values was set to zero for each track, and these new coordinates were plotted in Microsoft Excel to generate migration tracks. Velocity was calculated by computing displacement between two consecutive frames. Displacement was then divided by time interval to obtain speed for each cell. These speed values were then time-averaged over all frames to produce data points for cell speed which were plotted as box-and-whisker graphs. Alternatively, a custom-written Python based workflow, which used similar segmentation and tracking strategy, was also used for quantifying migration velocity. Please see the GitHub repo for details.

---

**Figure S1.**
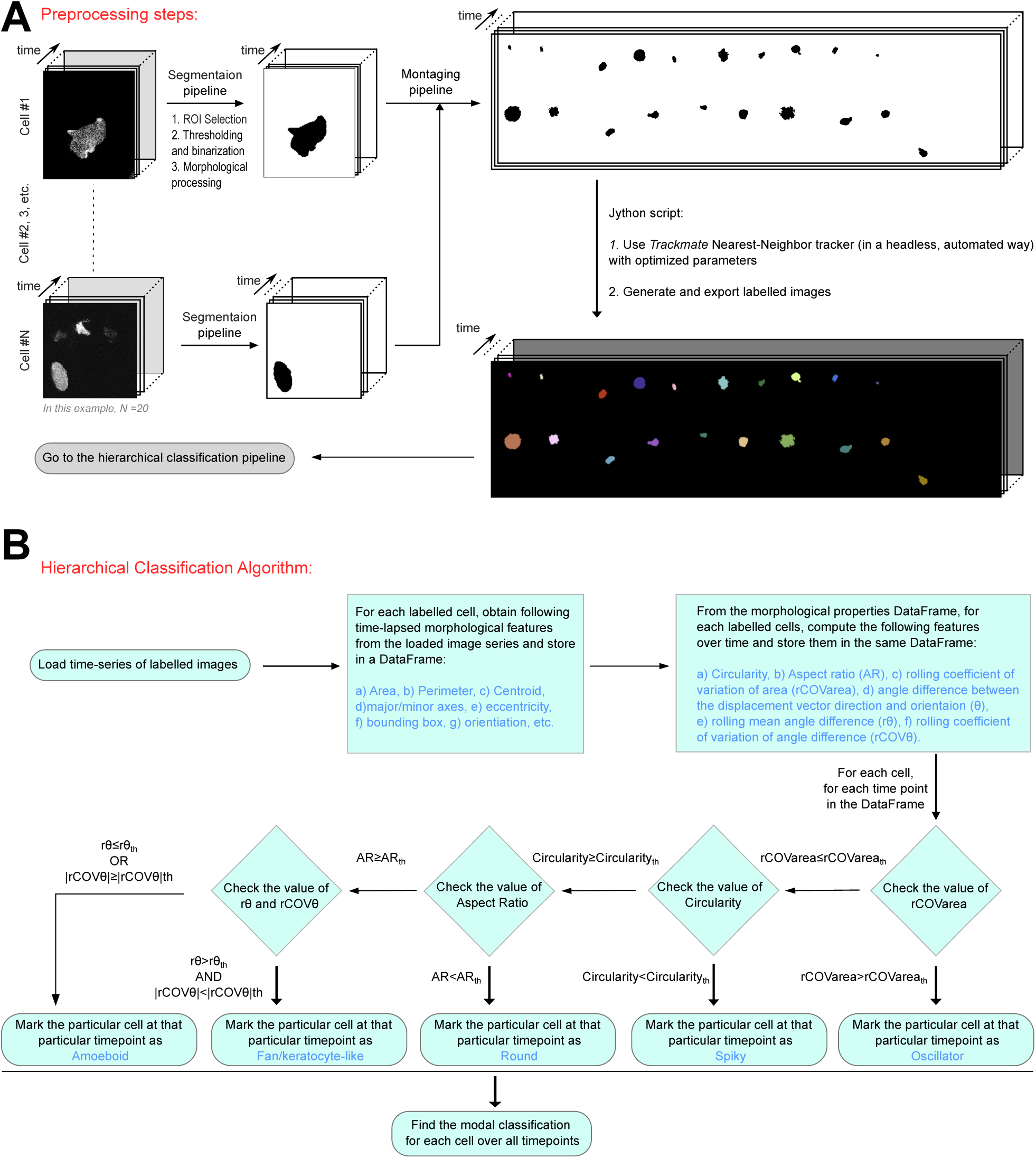
Automated classification of cellular morphology and migration mode by hierarchical clustering. **(A)** Preprocessing steps before cell classification that accept multiple time-lapse image series as input and generate a combined “labelled” image time-series where each label corresponds to a particular cell across the time-series. Throughout Figure S1 and S2 examples, N=20 cell images were used and each cell image series contains 25-30 time points. For details on segmentation pipeline, montaging pipeline, and Jython-based please see Methods and code provided (we are providing the full Jupyter Notebooks and Jython codes on GitHub: https://github.com/tatsatb/PIP5K-Ras-bistability-symmetry-breaking). **(B)** Hierarchical clustering algorithm delineating how the **“**labelled” cells were classified into “*oscillator*”, “*spiky*”, “*round*”, “*fan/keratocyte-like*”, and “*amoeboid*” categories, based on multiple metrics. Following threshold values are selected based on Kernel Density Estimation plots (please see figure S2B-S2F): rCOVarea_th_: 18; Circularity_th_: 0.4; AR_th_: 1.2; rθ_th_: 40; |rCOVθ|_th_: 100. Each cell in every frame of labelled image series has been classified into one of these categories and finally the modal values are chosen to finally categorize each cell. Please see Figure S2, methods, and GitHub repository for more details.

**Figure S2.**
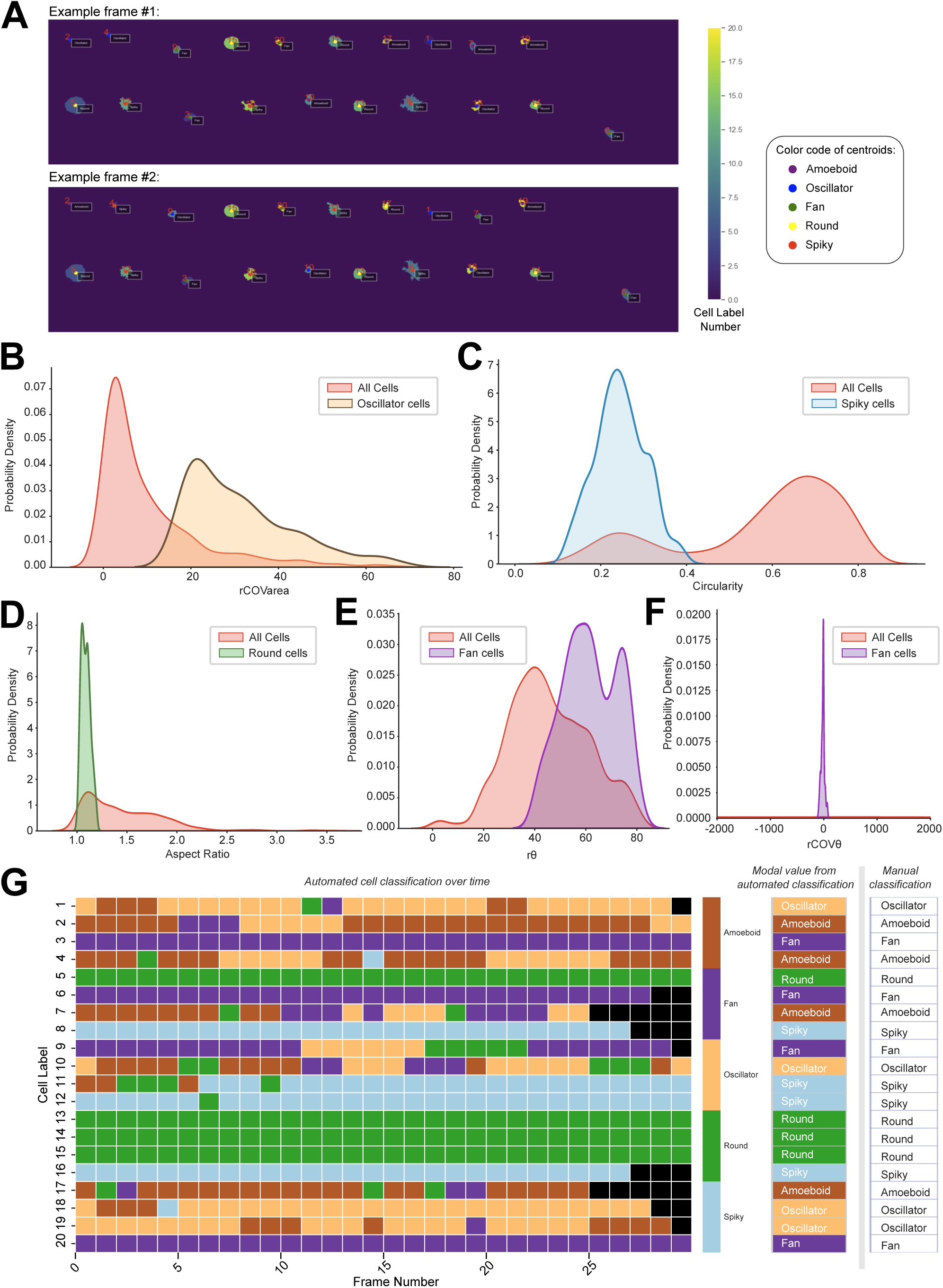
Metrics for the unbiased hierarchical classification of cells and comparison with manual classification. **(A)** Two example time-points from the “labelled” image time-series. Note that each cell has a specific label number between 1-20 (corresponding to matplotlib viridis colormap is shown on right) and each cell has been categorized into one of the five categories and their centroids are marked accordingly with color-code. **(B–F)** Kernel density estimation plots of rolling coefficient of Variation of area (B), circularity (C), aspect ratio (D), rolling mean angle difference (E), rolling coefficient of variation of angle difference (F) for entire population across all time frames are shown in red. The population that got selected into a particular category based on specific threshold values for each metric (see Figure S1B, methods, and https://github.com/tatsatb/PIP5K-Ras-bistability-symmetry-breaking for details) is shown in another color. **(G)** Heatmap showing classification of cells into one of the five categories across different time points. Black represents time frames where a particular cell was not present in the labelled image canvas. For comparison purpose, Manual classification (which was done independently of automated classification) categories are shown on the right of modal values from automated classification.

**Figure S3.**
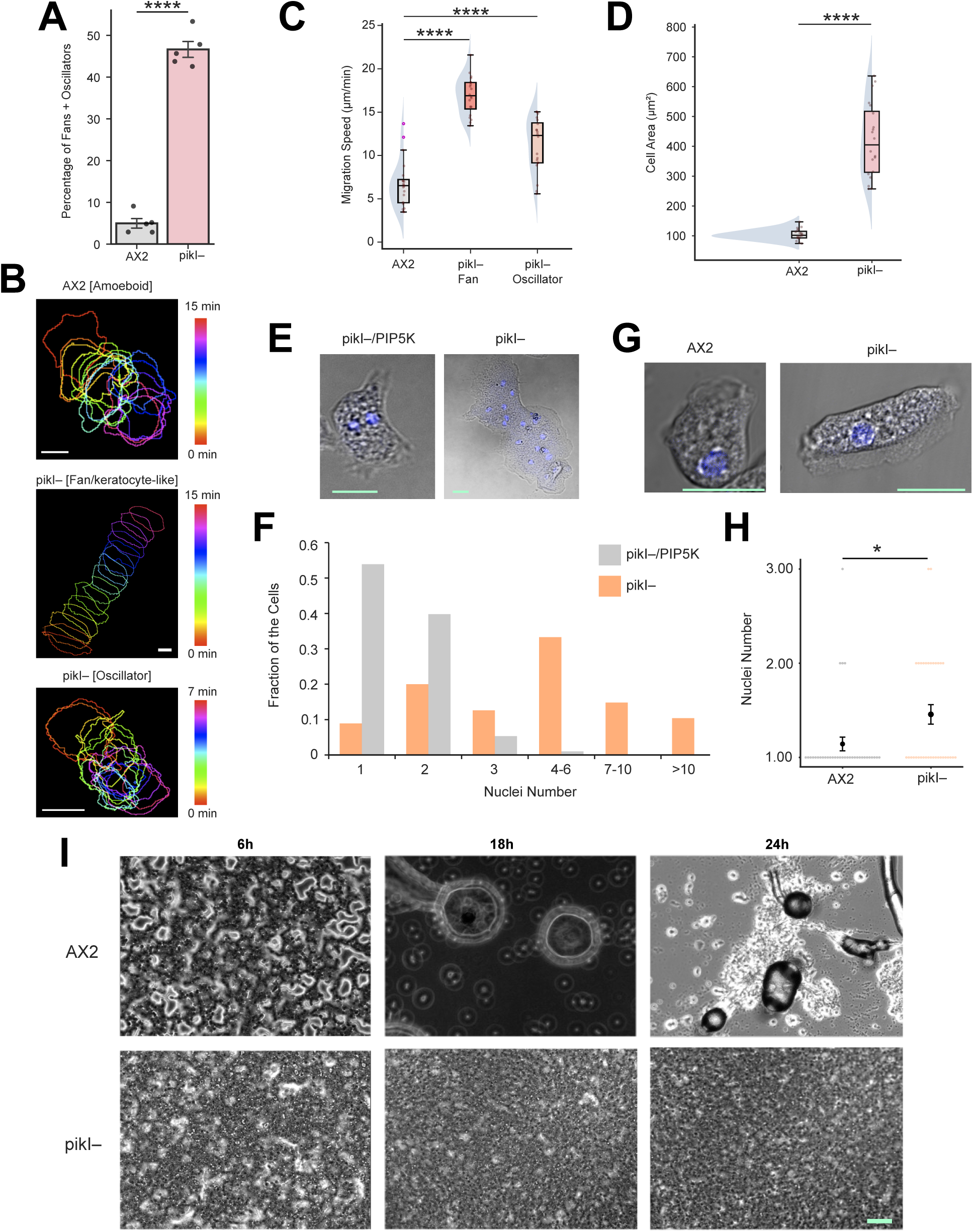
PIP5K knockout cells exhibit faster migration, impaired cytokinesis, and defective aggregation. **(A)** Representative color-coded temporal overlay profiles of migrating Dictyostelium cells; wild-type AX2 cells (top), *pikI–* fan/keratocyte-like, (middle) and *pikI–* oscillatory cells (bottom) are shown which are analogous to Figure 1A. The color maps denoting the time are shown on the right. **(B)** Quantification of percentage of cells migrating as fans and oscillator in wild-type AX2 and *pikI– Dictyostelium* cell populations. Each data point indicates one independent experiment where at least 46 cells were profiled. The p-values by Welch’s t-test. **(C)** Quantification of cell migration speed of AX2 cells (amoeboid mode migration) and *pikI–* cells (migrating as either fans or oscillators), showing that fan-shaped cells migrate the fastest, while oscillator cells migrate faster than the wild-type cells but slower than the fan-shaped cells. Here n_c_ =20 cells for each case; p values by Mann-Whitney U test. **(D)** Quantification of cell area of AX2 cells and *pikI–* cells showing *pikI–* cells are flatter compared to AX2 cells. Here n_c_ =20 cells for each case; p values by Mann-Whitney U test. **(E, F)** Representative images (E) and histogram of nuclei number (F) of AX2 cells and *pikI–* cells grown in suspension culture for 65 hours. In (E), DIC channels and nuclear staining (Hoescht) channels are merged. In (F) n_c_ ≥ 135 cells were analyzed for each case; data is from N = 2 independent experiments. **(G, H)** Representative images (G) and swarmplot of nuclei number (H) of AX2 cells and *pikI–* cells grown in adherent culture, showing significant mononucleated population in either case. In (G), DIC channels and nuclear staining (Hoescht) channels are merged; p values by Mann-Whitney U test. **(I)** AX2 (top) and *pikI–* cells (bottom) were plated on development buffer agar for starvation induced aggregation and imaged at 6h, 18h, and 24h, showing *piki-* cells have severe aggregation defects.

**Figure S4.**
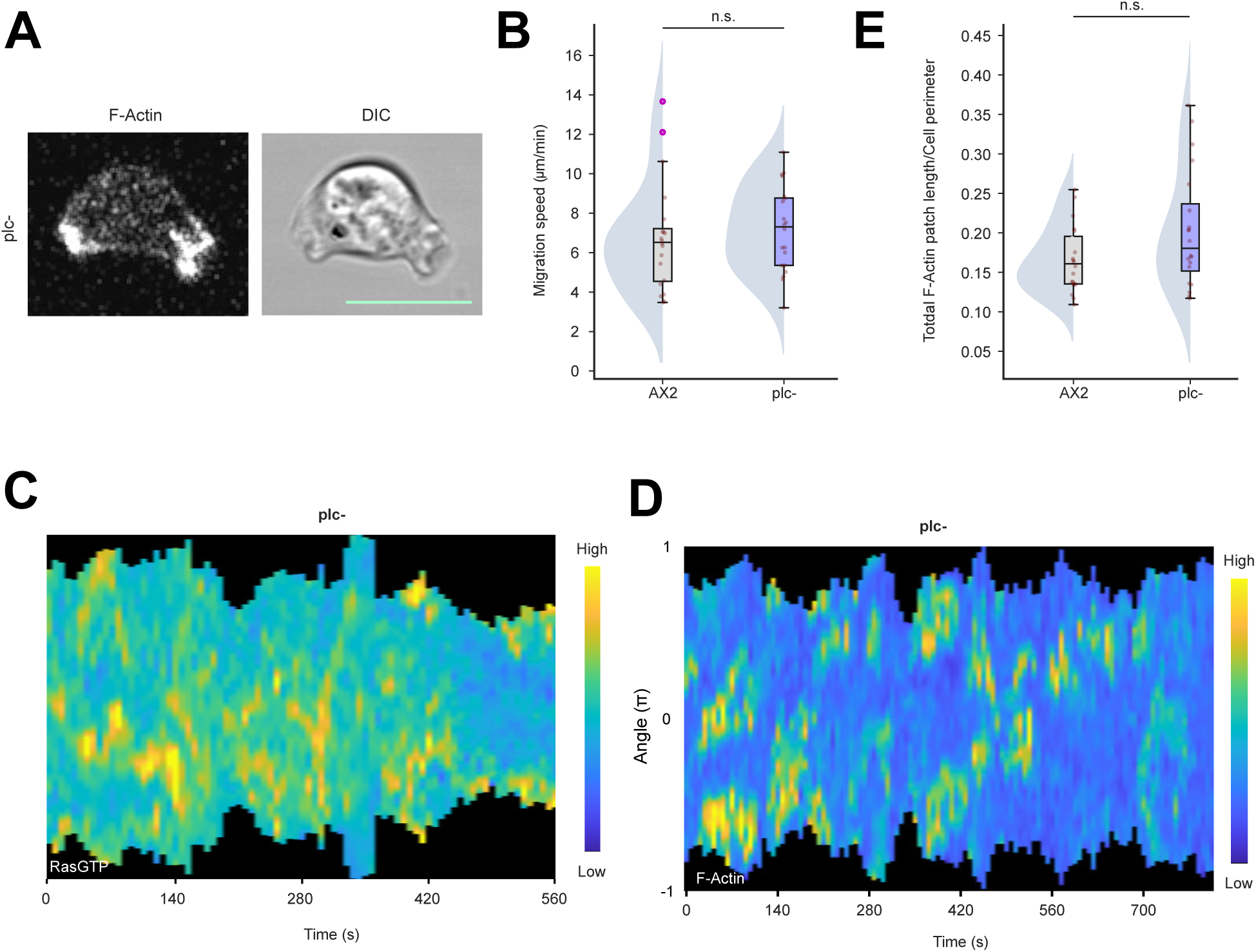
PLC knockout cells have no substantial phenotypic defect in terms of morphology, migration, or signaling and cytoskeletal activities. **(A)** Live-cell confocal images of *plc– Dictyostelium* cells expressing LimE-mCherry (LimE_Δcoil_, ‘LimE’ hereafter, is a biosensor for newly polymerized F-actin). **(B)** Quantification of cell migration speed of AX2 cells (amoeboid mode migration) and *plc–* cells, showing that there is no significant difference in speed between these two populations. Here n_c_ =20 cells for each case; p values by Mann-Whitney U test. **(C, D)** Representative 360° membrane kymographs from *plc– Dictyostelium* cells, expressing either RBD-GFP (RBD:Ras-Binding Domain of Raf1 that serves as a biosensor of Ras activation) (C) or LimE-mCherry (D). (C) corresponds to cells shown in Figure 1D and (D) corresponds to cells shown in Figure S4A. **(E)** Quantification of overall newly polymerized F-actin level on the cell membrane in AX2 and *plc–* cells, in terms of the total LimE patch length/cell perimeter, demonstrating no significant difference between these two populations. Here n_c_ =20 cells for each case (here and everywhere else in this study, for patch length quantification each of the n_c_ cells were tracked for at least n_f_=5 frames and the average values were obtained). P-values by the Mann-Whitney U test.

**Figure S5.**
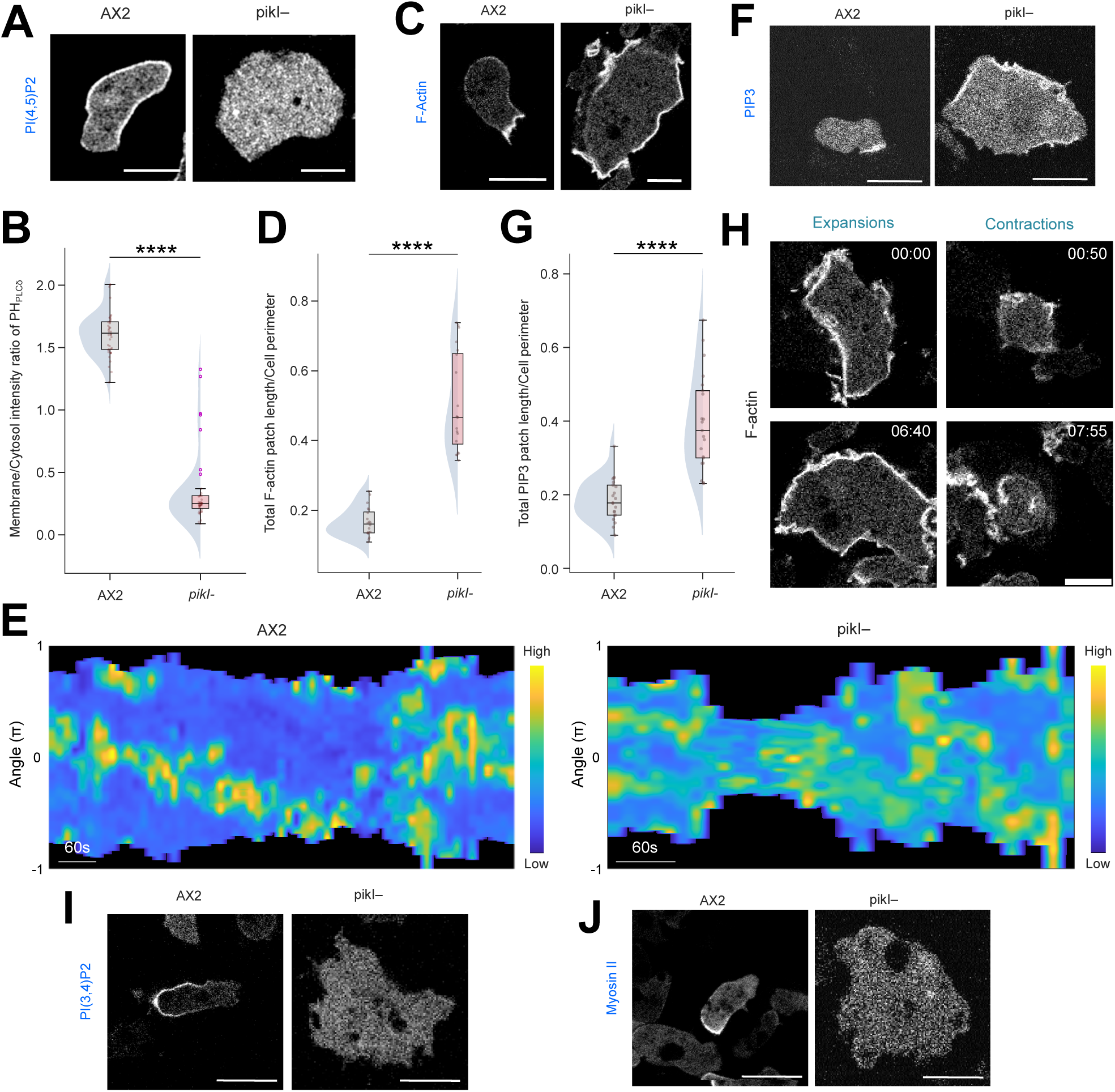
PIP5K knockout cells display hyperactivated Ras/PI3K/F-actin network activities. (A, C, **F)** Representative live-cell images of WT *Dictyostelium* AX2 cells (left panels) and *pikI– Dictyostelium* cells (right panels) expressing PH_PLCδ_-YFP (PH_PLCδ_ is the biosensor for PI(4,5)P2) (A), or LimE-mCherry (C), or PH_crac_-mCherry (PH_crac_ is the biosensor for PI(3,4,5)P3). **(B, D, G)** Quantification of PI(4,5)P2 level (B), newly polymerized F-actin level (D), and PIP3 level (G) in the membranes-cortex of AX2 and *pikI–* cells, in terms of total PH_PLCδ_ (B)/LimE(D)/ PH_crac_ (G) patch length/cell perimeter. Boxplots demonstrate that PI3K/F-actin axis is highly active in *pikI–* cells, compared to AX2 cells, whereas PI(4,5)P2 levels are severely low. Here, for each case, n_c_ =40 cells (B), or 20 cells (D), or 21 cells (G); p values by Mann-Whitney U test. **(E)** Representative 360° membrane kymographs of AX2 and *pikI– Dictyostelium* cells, expressing either LimE-mCherry showing consistently enhanced F-actin polymerization in *pikI– cells*. **(H)** Live-cell time-lapse images of *pikI– Dictyostelium* oscillatory cells expressing LimE- mCherry undergoing typical oscillation. Left-panel time points are showing expanded state and right-panel time points are showing the contracted state. **(I, J)** Representative live-cell images of WT *Dictyostelium* AX2 cells (left panels) and *pikI– Dictyostelium* cells (right panels) expressing PH_CynA_-KikGR ( PH_CynA_ is the biosensor for PI(3,4)P2) (I), or Myosin II-GFP (J), demonstrating low PI(3,4)P2 level and impaired myosin II assembly in *pikI–* cells.

**Figure S6.**
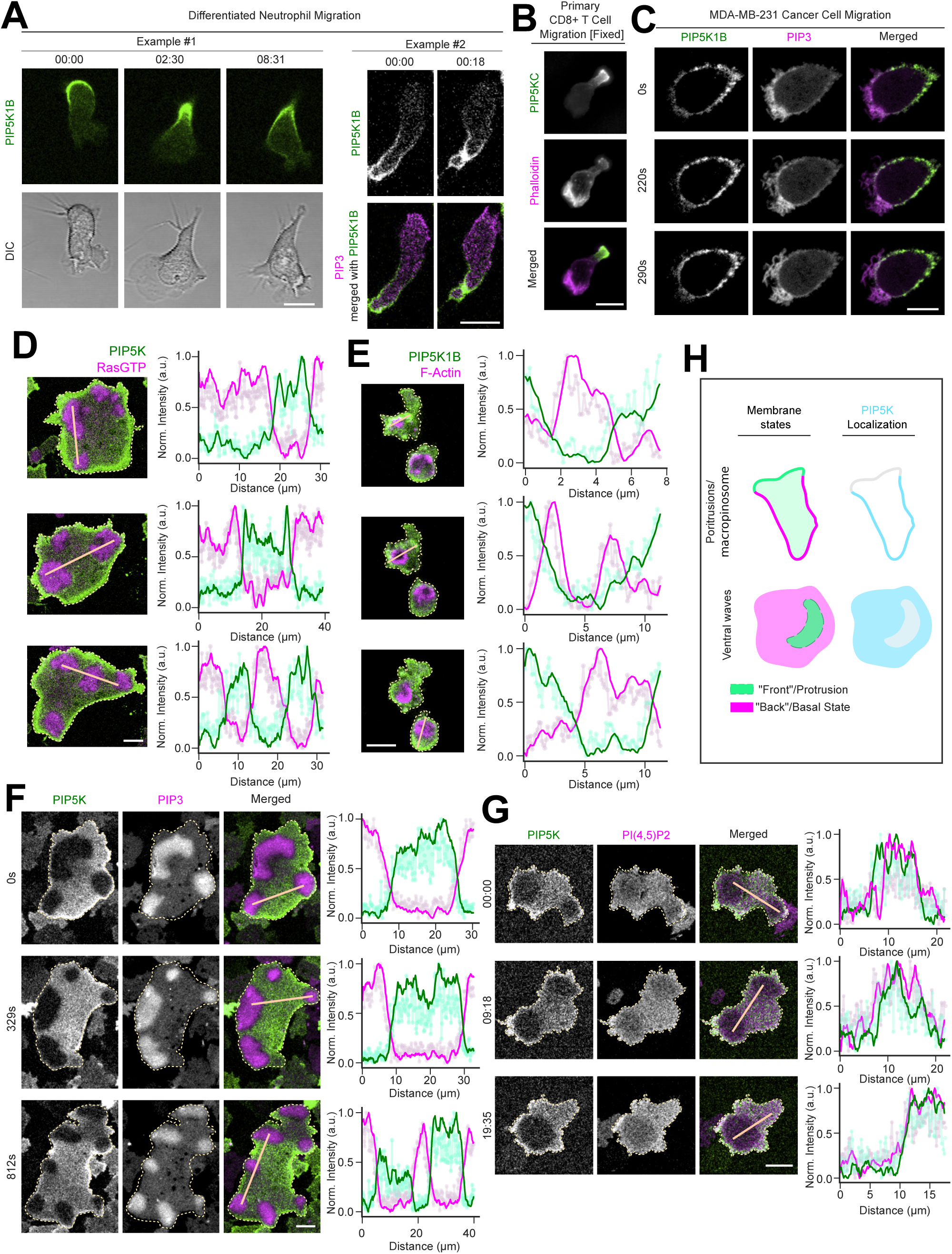
PIP5K dynamically localizes within the back-state regions of the plasma membrane under different physiological scenarios in different cell types. **(A)** Live-cell time-lapse images of differentiated HL-60 neutrophils expressing PIP5K1B- GFP (along with PH_Akt_-mCherry in Example #2; PH_Akt_ is a biosensor for PIP3) during migration, showing PIP5Ks dynamically move away from protrusions in migrating cells which are enriched in PIP3. Time in min:sec format. **(B)** Primary mouse CD8+ T cells expressing PIP5K1C-mNeon were allowed to migrate on ICAM-1 coated coverslips for 20mins, fixed, and stained with phalloidin **(C)** Live-cell time-lapse images of MDA-MB-231 breast cancer cells expressing PIP5K1B- GFP and PH_Akt_-mCherry during migration, showing PIP5Ks are depleted from protrusions in migrating cells. Time in sec format. **(D, E)** Line-scan intensity profiles are shown which correspond to Figure 1J and 1K, respectively. Intensity profiles were obtained across the thick lines shown in the panels; lighter colored lines join the raw data points whereas darker colored lines display the moving averages. Note that PIP5Ks maintain complementarity with the membrane regions that have high RasGTP activities or the regions that are enclosed by newly polymerized F-actin waves. **(F, G)** Ventral wave propagation in *Dictyostelium* cells coexpressing PIP5K-GFP and PH_crac_- mCherry (F) or mRFP-PIP5K and PH_PLCδ_-GFP (G) showing PIP5K and PH_PLCδ_ dynamically and consistently localizes to the back-state regions in ventral waves. Line-scan intensity profiles (similar to D, E) are shown in the rightmost panels. **(H)** Schematic illustrating the spatial distribution of PIP5K in the plasma membrane during protrusion/macropinosome formation and ventral wave propagation. Demarcated front and back state regions of the plasma membrane in these physiological scenarios are shown in the left panels.

**Figure S7.**
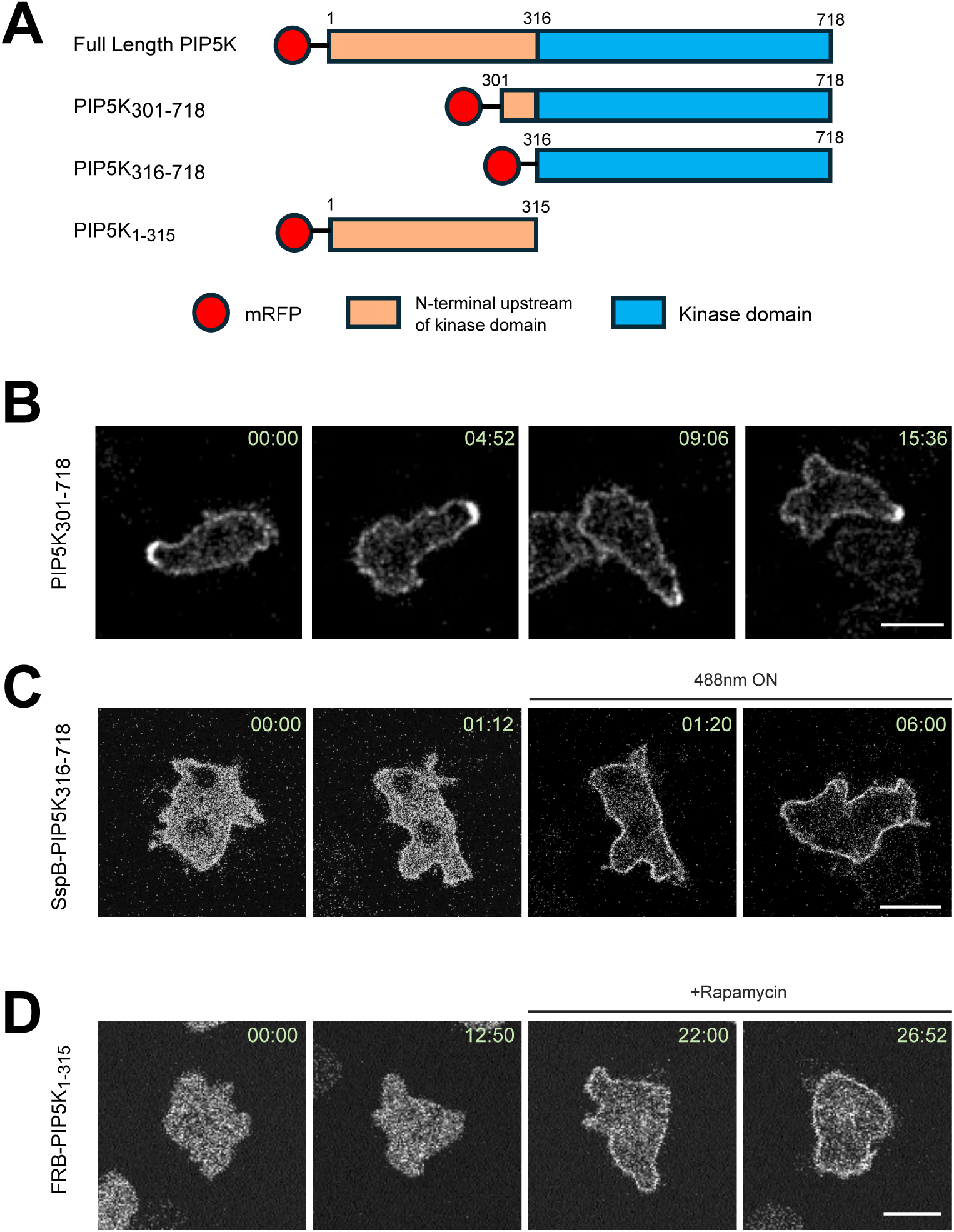
Profiling the ability of different regions of PIP5K protein in sorting into the back-state of the membrane. **(A)** Diagram representing the design of different truncated versions of PIP5K. **(B)** Live-cell images of migrating *Dictyostelium* cells expressing mRFP-PIP5K_301-718_ during migration, showing PIP5K_301-718aa_ dynamically localizes at the trailing edge. In this figure, all times are in min:sec format. **(C)** Live-cell images of migrating *Dictyostelium* cells expressing mRFP-SspB_R73Q_- PIP5K_301-718_ and cAR1-iLiD, before and after globally turning on 488 nm laser to facilitate recruitment of cytosolic PIP5K_301-718_. **(D)** Live-cell images of migrating *Dictyostelium* cells expressing mCherry-FRB-PIP5K_316- 718_ and cAR1-FKBP-FKBP, before and after addition of 5 μM Rapamycin to facilitate recruitment of cytosolic PIP5K_316-718_.

**Figure S8.**
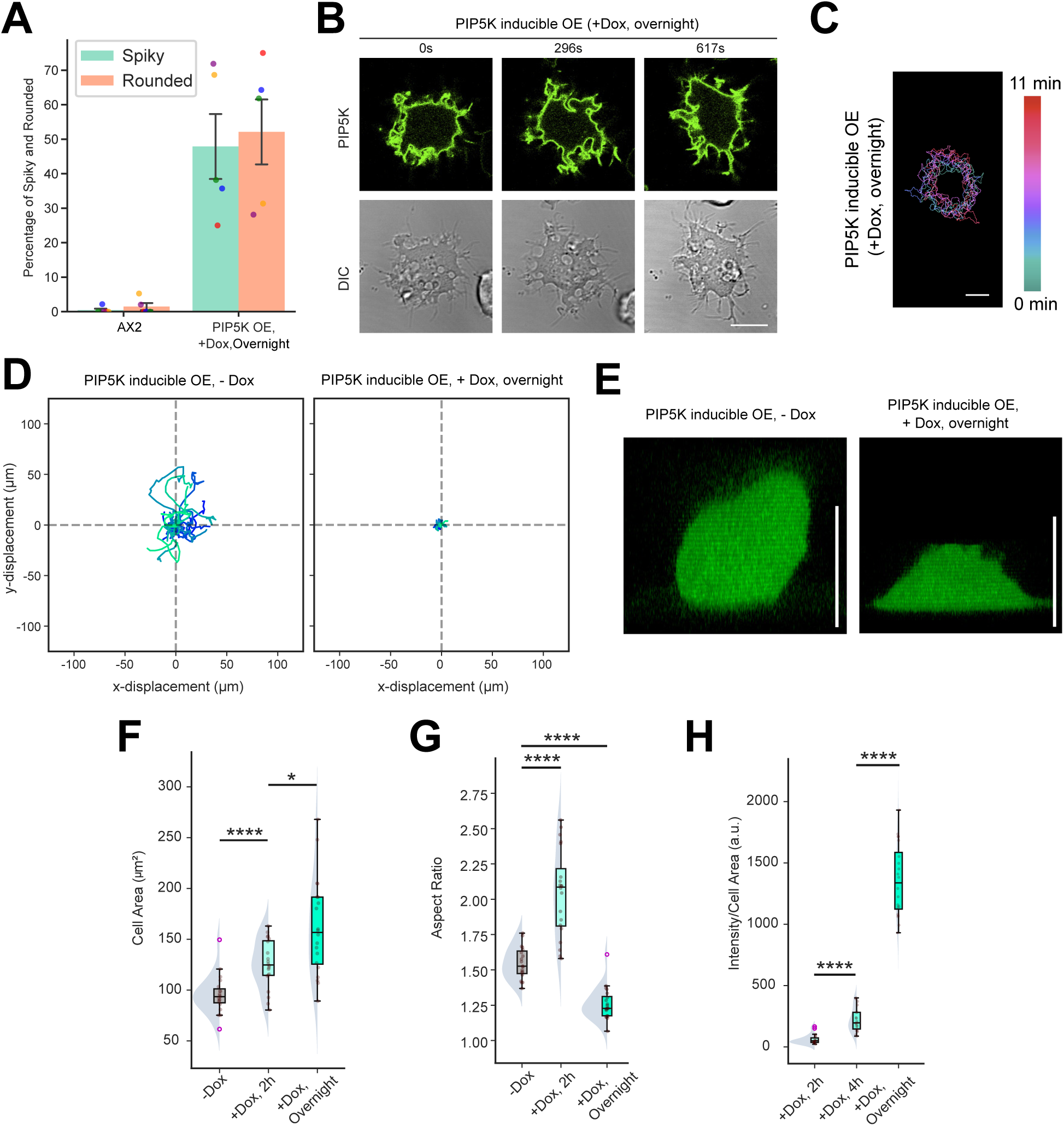
Overnight doxycycline treatment driven PIP5K inducible overexpression substantially alter *Dictyostelium* cell migration mode and morphological features. **(A)** Quantification of percentage of cells behaving as “spiky” and “rounded” in wild-type AX2 and PIP5K-GFP OE (upon overnight Dox treatment) *Dictyostelium* cell populations. Each data point indicates one independent experiment where at least 55 cells were profiled. Data points are color-coded so that, for either AX2 or PIP5K OE group, same color represents the same experiment, providing a way to track fraction of “spiky” vs “rounded” in each experiment. **(B, C)** Live-cell time-lapse images (B) and Color-coded temporal overlay profiles (C) of *Dictyostelium* cells expressing Dox-inducible PIP5K with overnight Dox treatment, which resulted a spiky cell phenotype that barely migrate. **(D)** Centroid tracks of randomly migrating *Dictyostelium* cells (n_c_=20) expressing Dox-inducible PIP5K, without (left) or with overnight (right) Dox treatment. Each cell was tracked for 10 minutes and was reset to the same origin. **(E)** Representative z-stack imaging (rendered with *Volume* view of Fiji/ImageJ 3D Viewer plugin) showing the height of *Dictyostelium* cells expressing Dox-inducible PIP5K-GFP without Dox induction (left) or with overnight Dox induction (right), showing PIP5K OE cells are substantially flatter. **(F-H)** Quantification of cell area (F), aspect ratio (G), and PIP5K-GFP Intensity per unit cell area (H) of *Dictyostelium* cells expressing Dox-inducible PIP5K-GFP, either without or with Dox treatment from specified amount of times. Noticeably, upon increased expression of PIP5K-GFP, cells spread significantly and also, cells eventually lost polarity after going through a phase of elongation when PIP5K was moderately overexpressed. Here, n_c_=19 for -Dox population and n_c_=20 for +Dox population; data from N=3 independent experiments; p value by the Mann-Whitney U test.

**Figure S9.**
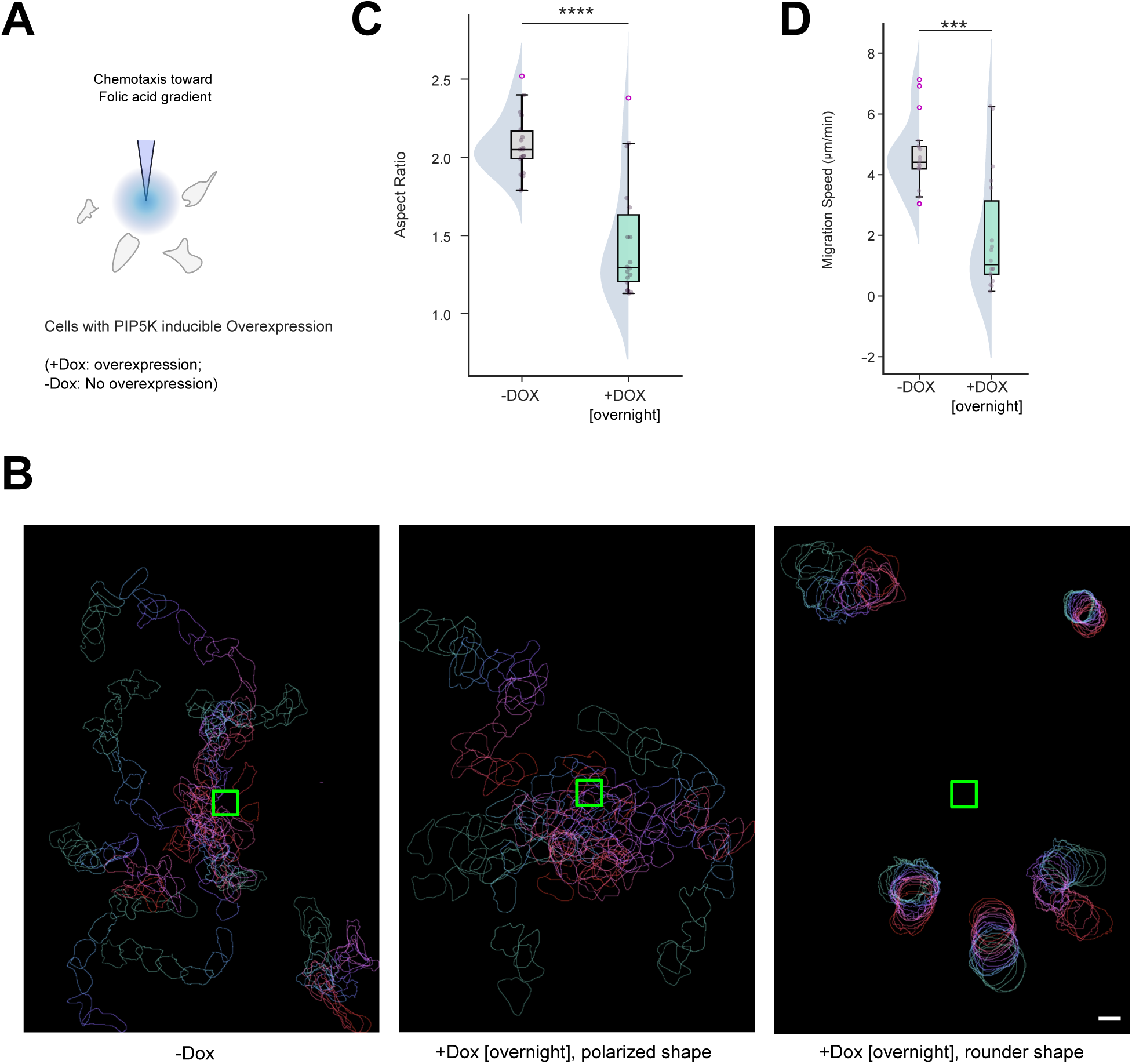
Inducible PIP5K overexpression impairs chemotaxis. **(A)** Illustration of chemotaxis of *Dictyostelium* cells towards folate gradient, generated by placing a folate-filled micropipette connected to a controllable microinjector. Chemotaxis assay was performed with *Dictyostelium* cells expressing Dox-inducible PIP5K-GFP, either without or with overnight Dox treatment. **(B)** Color-coded temporal overlay of non-Dox treated cells (left panel), Dox-treated cells which acquired a polarized shape upon gradient stimulation (middle panel), and Dox-treated cells which maintained round-shape upon gradient stimulation (right panel). Green box is showing the tip of the needle where folate is coming from. **(C, D)** Quantification of the aspect ratio (C) and migration speed (D) of non-Dox treated and overnight Dox-treated cells under chemotactic gradient, showing PIP5K OE cells abrogate their polarity and cannot chemotax effectively. Here n_c_=18 for either -Dox population or +Dox population; p value by the Mann-Whitney U test.

**Figure S10.**
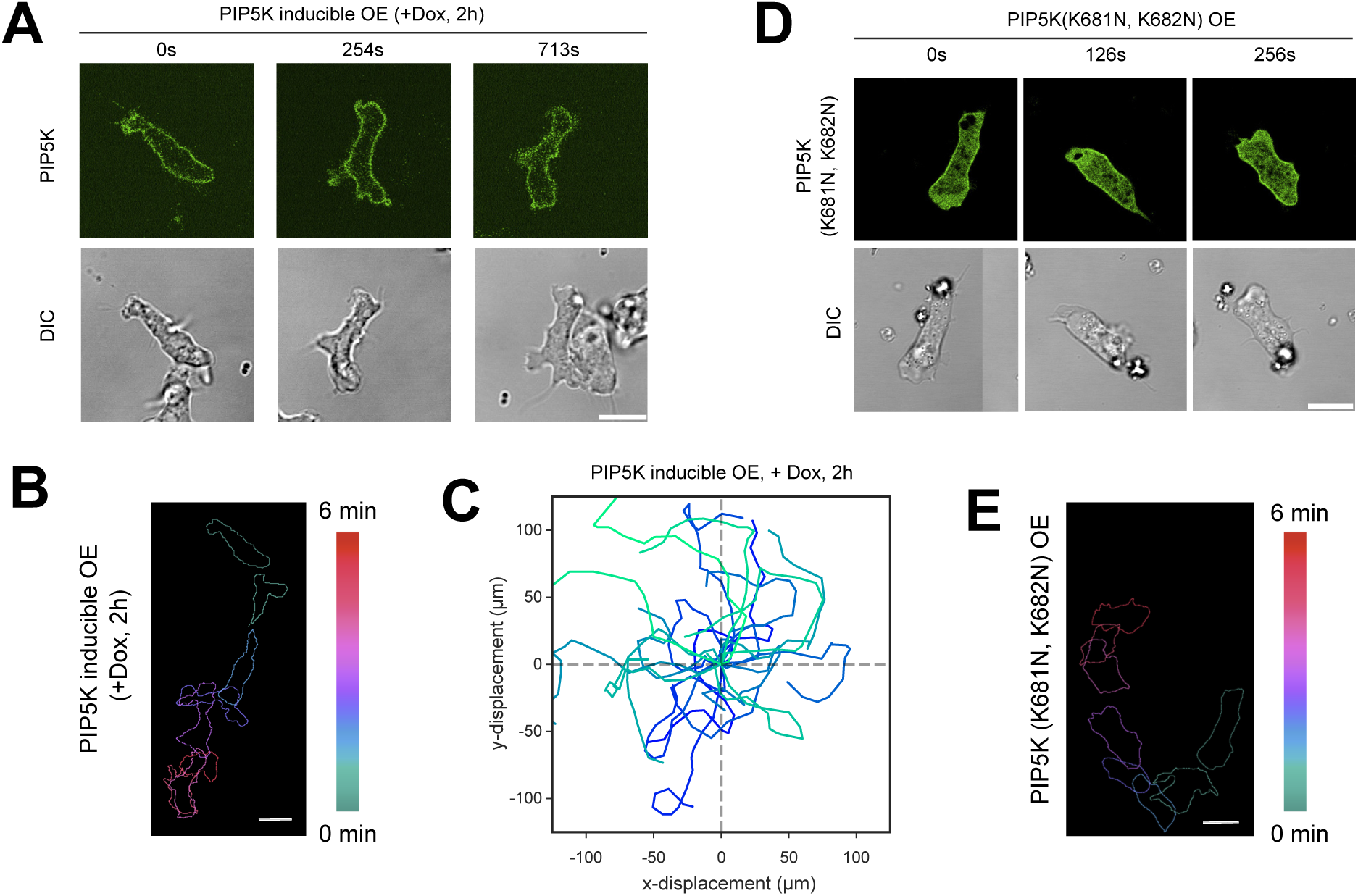
Moderate overexpression of PIP5K polarizes cells and make them migrate faster. **(A, D)** Representative live-cell images of *Dictyostelium* cells expressing Dox-inducible PIP5K-GFP with 2h Dox induction (A) or expressing PIP5K (K681N, K682N)-GFP mutant (D), showing both of these populations are more elongated and polarized. **(B)** Color-coded temporal overlay profiles of *Dictyostelium* cells expressing doxycycline-inducible PIP5K-GFP with 2 hour long Dox induction, showing that this moderate overexpression of PIP5K made these cells polarized and fast. **(C)** Centroid tracks of randomly migrating *Dictyostelium* cells (n_c_=20) expressing Dox-inducible PIP5K, with 2 hour long Dox treatment. Each cell was tracked for 10 minutes and was reset to the same origin. **(E)** Color-coded temporal overlay profiles of *Dictyostelium* cells expressing PIP5K (K681N, K682N)-GFP mutant, showing that this mutant PIP5K expression made cells elongated and fast.

**Figure S11.**
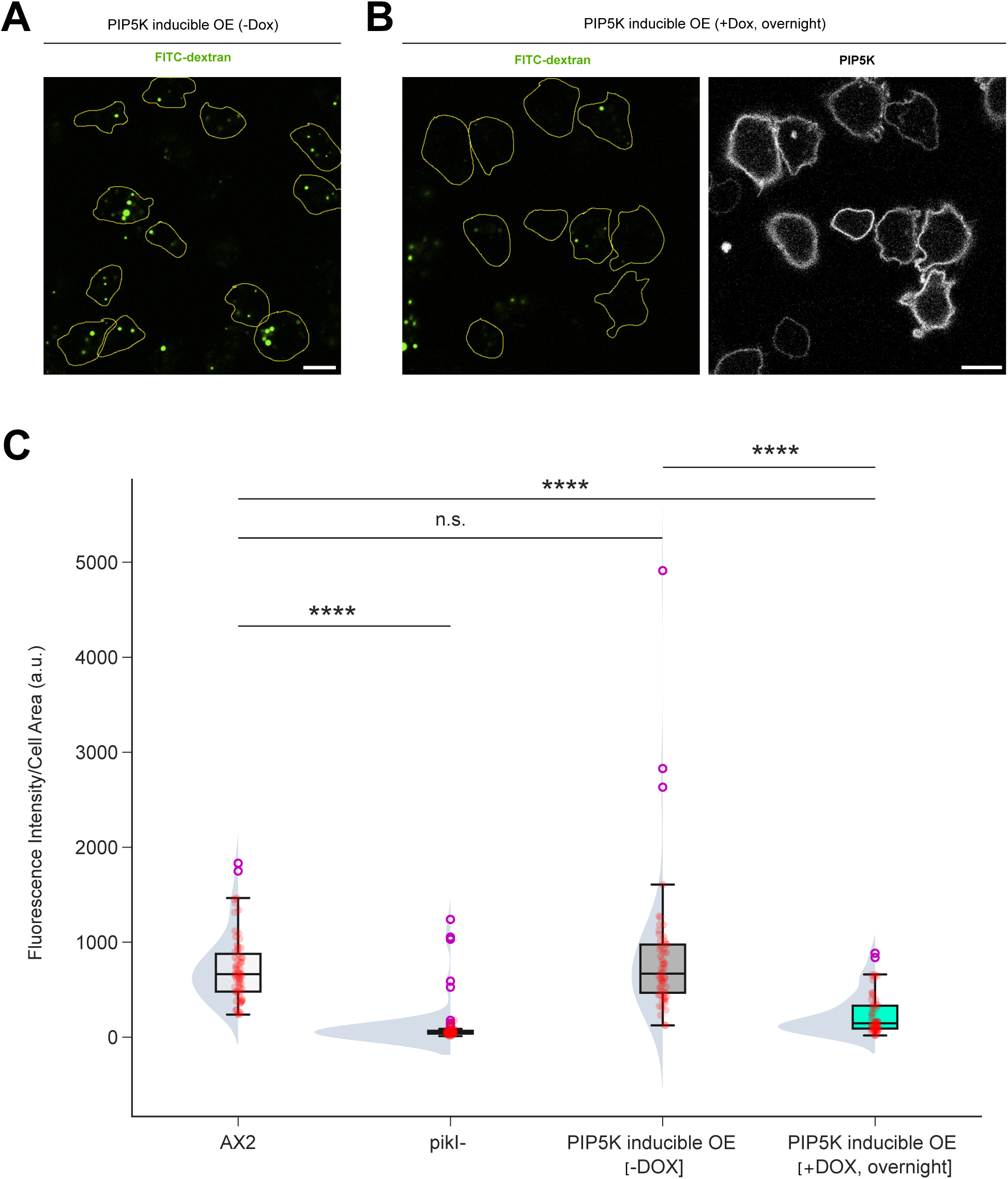
Both PIP5K knockout and PIP5K overexpression result in decrease in macropinocytosis. **(A, B)** Representative images of *Dictyostelium* cells, which are expressing Dox-inducible mRFP-PIP5K, without (A) or with overnight (B) Dox induction, performing macropinocytosis. Cells were treated with FITC-dextran (green) for 10mins before imaging. The yellow lines show the cell outlines. **(C)** Quantification of macropinocytosis in AX2, *pikI–,* PIP5K inducible OE (-Dox), and PIP5K inducible OE (+Dox, overnight) *Dictyostelium* cells, in terms of green fluorescence intensity per unit cell area, demonstrating that either the absence of PIP5K or the increased level of PIP5K result in strong macropinocytosis defects. Here, n_c_=66 (AX2), 70 (*pikI–*), 57 (PIP5K inducible OE (-Dox)), or 58 (PIP5K inducible OE (+Dox, overnight)); p value by the Mann-Whitney U test.

**Figure S12.**
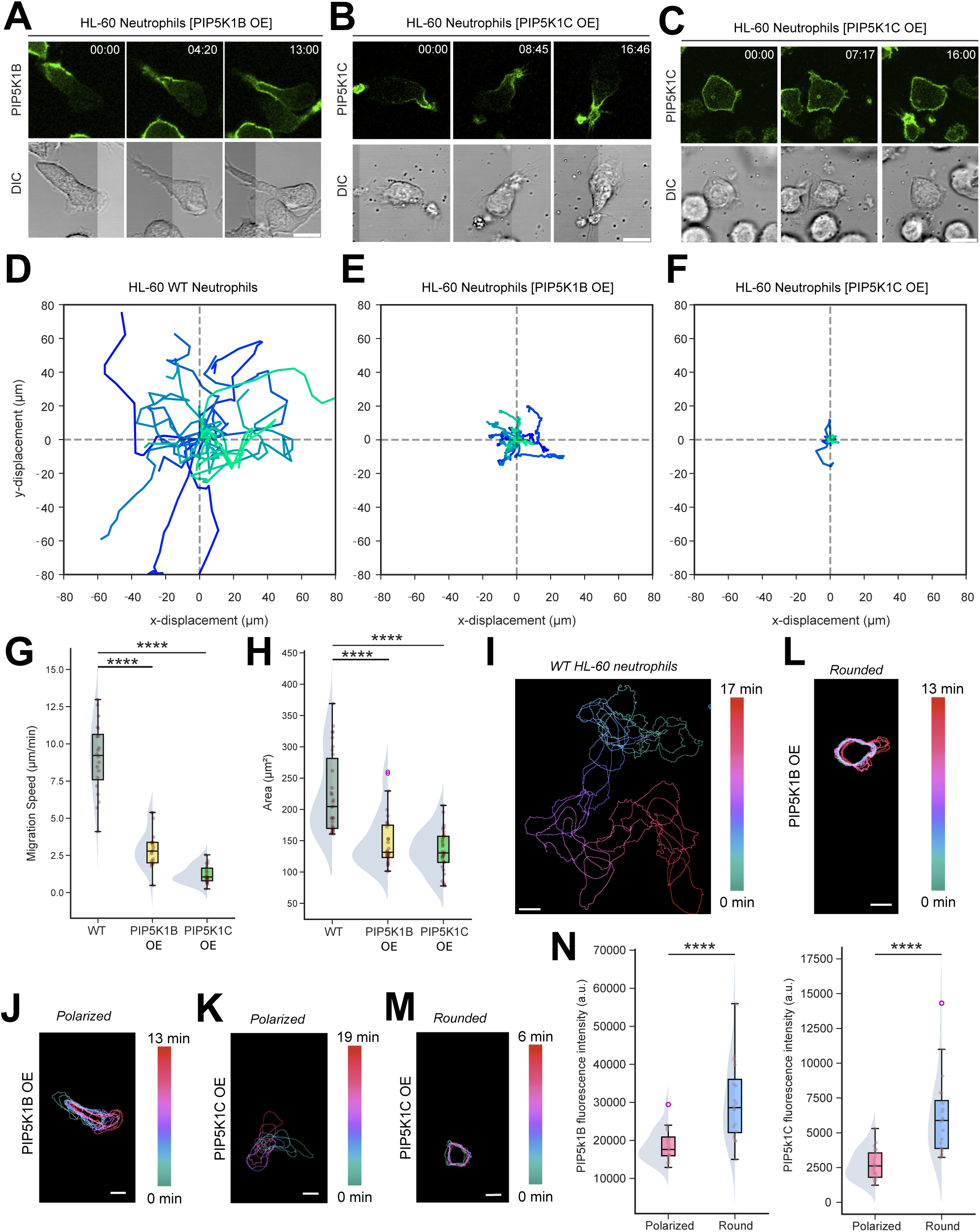
Overexpression of PIP5K impairs HL-60 neutrophil polarization and migration. **(A-C)** Representative live-cell time-lapse images of HL-60 cells differentiated into neutrophils, expressing PIP5K1B-GFP (A) or PIP5K1C-GFP (B, C), demonstrating that they can exhibit either polarized (with long PIP5K enriched tail) (A, B) or rounded (with PIP5K present nearly uniformly on the membrane) morphology. **(D-F)** Centroid tracks of randomly migrating differentiated HL-60 cells with WT background (D), PIP5K1B-GFP overexpression (E), and PIP5K1C-GFP overexpression (F), showing both PIP5K1B OE and PIP5K1C OE cells have drastic migration defects. Each of these n_c_=20 tracks lasted for 12 minutes and was reset to the same origin. **(G, H)** Quantification of migration speed and area of differentiated WT HL-60 neutrophils, HL-60 neutrophils with overexpressing PIP5K1B-GFP, and HL-60 neutrophils with overexpressing PIP5K1C-GFP. Here, n_c_=20 (G) or 30 (H) from at least N=3 independent experiments; p value by the Mann-Whitney U test. **(I-M)** Color-coded temporal overlay profiles of differentiated WT HL-60 neutrophils (I), or neutrophils overexpressing PIP5K1B-GFP (polarized) (J), or neutrophils overexpressing PIP5K1C-GFP (round) (K), or neutrophils overexpressing PIP5K1B-GFP (round) (L), or neutrophils overexpressing PIP5K1C-GFP (polarized) (M), showing that overexpressing PIP5Ks inhibited migration in both round and polarized cells comparing with WT, and round cells barely moved at all. **(N)** Quantification of the mean fluorescence intensity of PIP5K1B-GFP (left) and PIP5K1C-GFP (right) between polarized and round phenotypes, showing round OE cells have higher PIP5K expression level. Here n_c_=20 for either polarized or round population; p values by the Mann-Whitney U test.

**Figure S13.**
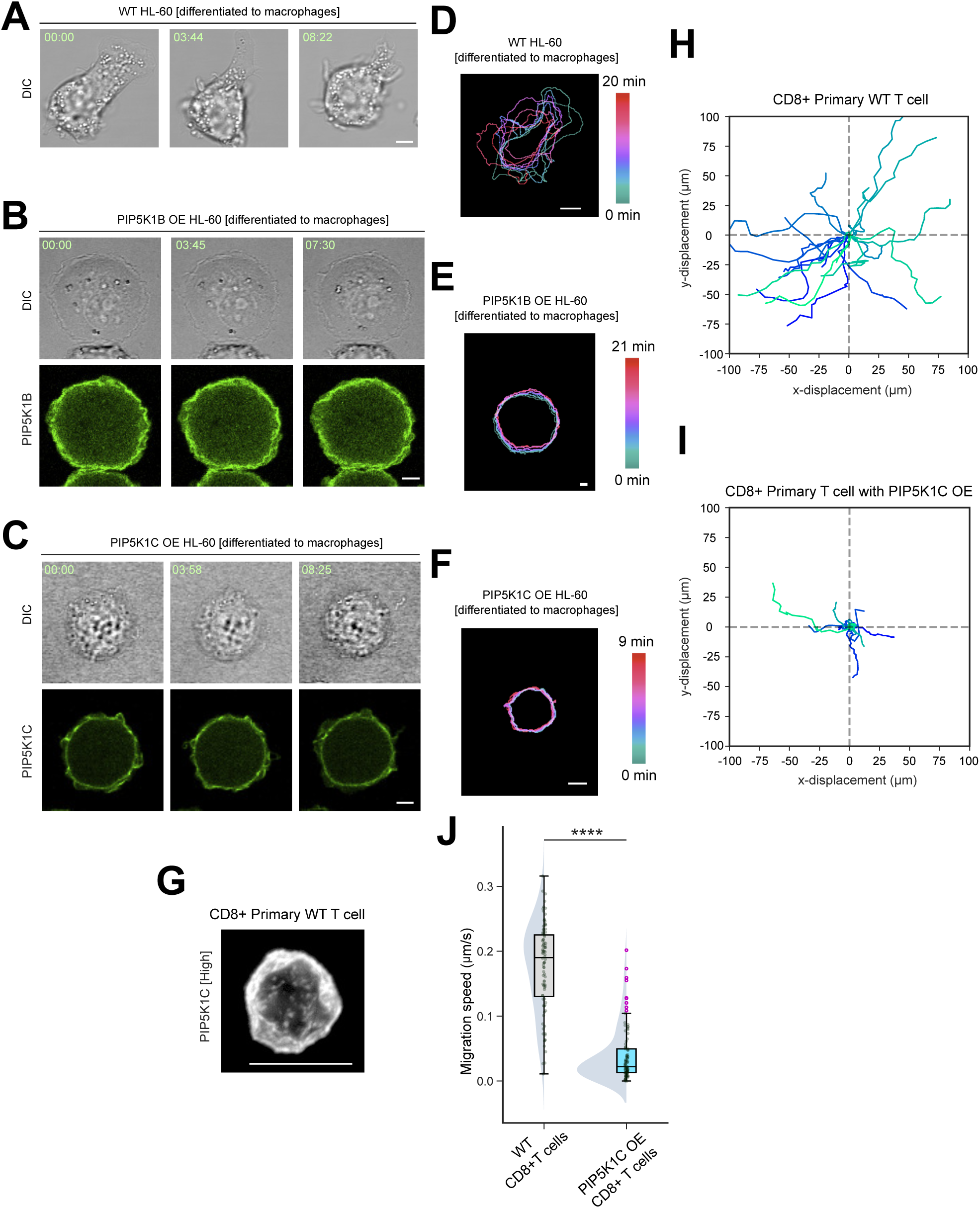
Overexpression of PIP5K impairs HL-60 macrophage protrusion formation and mouse primary CD8+ T cell migration. **(A-C)** Representative live-cell time-lapse images of differentiated WT HL-60 macrophages (A), or macrophages overexpressing PIP5K1B-GFP (B) or PIP5K1C-GFP (C), demonstrating that overexpression of PIP5Ks make cells flatter and rounded. **(D-F)** Color-coded temporal overlay profiles of macrophage cells shown in (A-C) respectively, showing that overexpressing PIP5Ks diminished protrusion formation and impaired migration. **(G)** Representative live-cell images of mouse primary CD8+ T cells expressing PIP5K1C-mNeon at higher level (compared to cell shown in Figure 1I), demonstrating a rounder cell shape. **(H-I)** Centroid tracks of randomly migrating mouse primary CD8+ T cells with WT background (H) or PIP5K1C-mNeon overexpression (I). Tracks collectively demonstrate that PIP5K1C OE cells have drastic migration defects. Each of these n_c_=20 tracks lasted for 10 minutes and was reset to the same origin. **(J)** Quantification of migration speed of WT mouse primary CD8+ T cells, or cells with PIP5K1C-mNeon overexpression. Here, n_c_=92 for WT cells, and 98 for PIP5K1C OE cells from at least N=3 independent experiments; p value by the Mann-Whitney U test.

**Figure S14.**
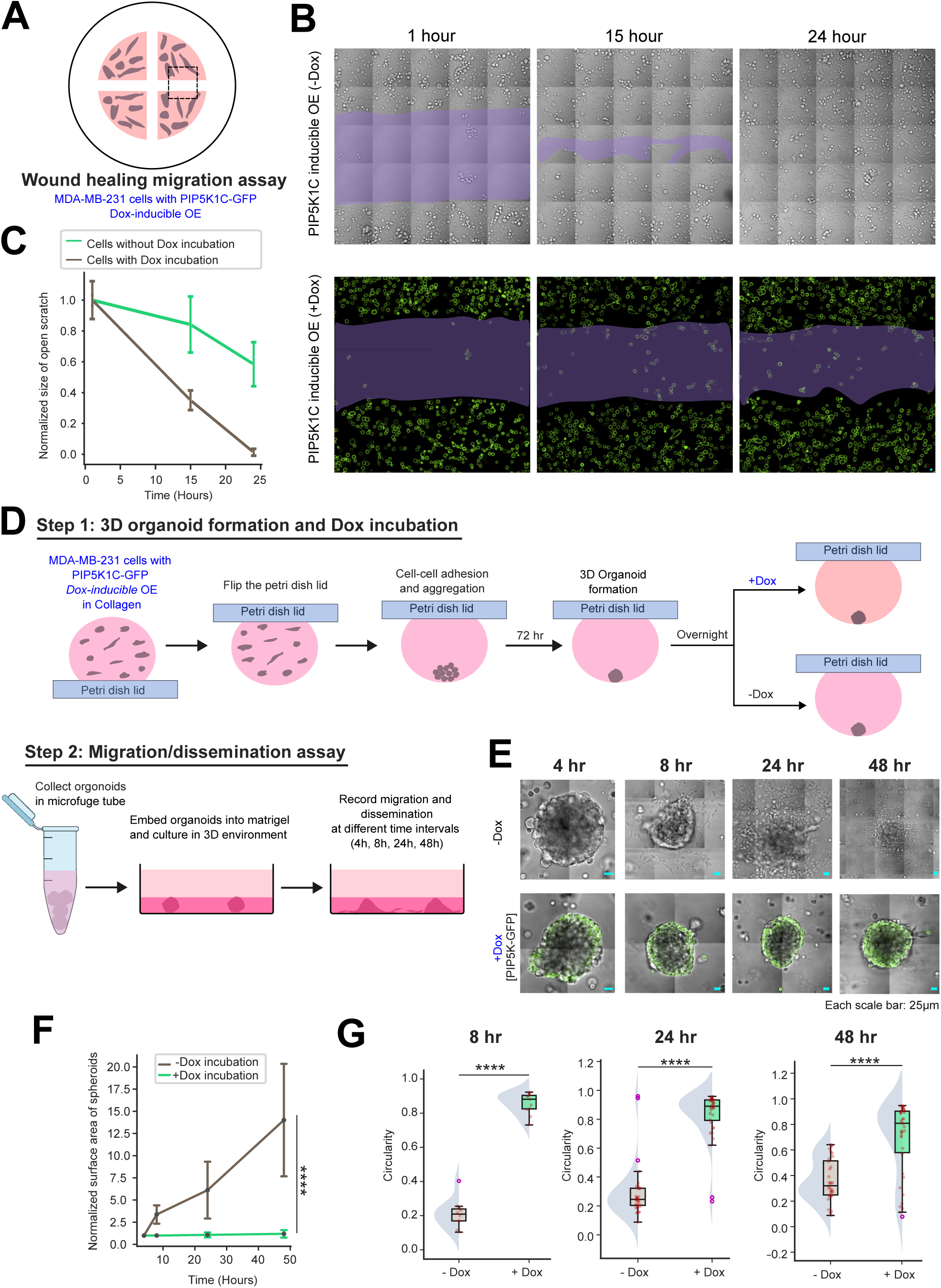
Overexpression of PIP5K inhibits the migration of MDA-MB-231 cancer cells in 2D wound healing and 3D organoid dissemination assays. **(A)** Illustration of 2D wound healing assay with MDA-MB-231 cells expressing Dox-inducible PIP5K1C-GFP; before plating the cells on wound healing chamber, they were incubated without or with Dox overnight. After plating the cells, the inserts were lifted to create an open scratch. **(B)** Representative live-cell images of MDA-MB-231 cells expressing Dox-inducible PIP5K1C-GFP, either without or with overnight Dox treatment. DIC images are shown for cells that were incubated without Dox and GFP channel are shown for cells that were incubated overnight with Dox. Images were acquired at respective time points. **(C)** Normalized size of the open scratch at different timepoints, demonstrating impeded wound closure speed in Dox incubated OE cells. Responses were computed in terms of open scratch area at each time point, normalized to scratch area at 1h. Here, n≥7 wounds for each case, for each time-point, from N≥3 independent experiments. **(D)** Schematic of dissemination assay from 3D organoids. Step 1 shows the 3D organoid formation from MDA-MB-231 cells (expressing Dox inducible PIP5K1C-GFP) using hanging drop cell culture protocol. After 3 days when organoids were formed, they were incubated without or with Dox overnight. In Step 2, 3D organoids were collected and embedded into Matrigel, and dissemination was recorded at the shown time points. **(E)** Representative live images of organoids at 4hr, 8hr, 24hr, and 48 hr from embedding. DIC images are shown for the organoids that were incubated without Dox and merged (GFP+DIC) images are shown for cells that were incubated overnight with Dox. **(F)** Normalized surface area of organoids at different time points after embedding in Matrigel, demonstrating PIP5K1C overexpressing organoids exhibit negligible dissemination, compared to their Dox untreated counterpart. **(G)** Circularity of the organoids incubated with or without Dox at different time-points showing that PIP5K1C-GFP overexpression significantly inhibits invasion and dissemination. Here, n=10 (-Dox, 8h), 11(+Dox, 8h), 33 (-Dox, 24h), 36 (+Dox, 24h), 32 (-Dox, 48h), 38 (+Dox, 48h) organoids; p value by the Mann-Whitney U test.

**Figure S15.**
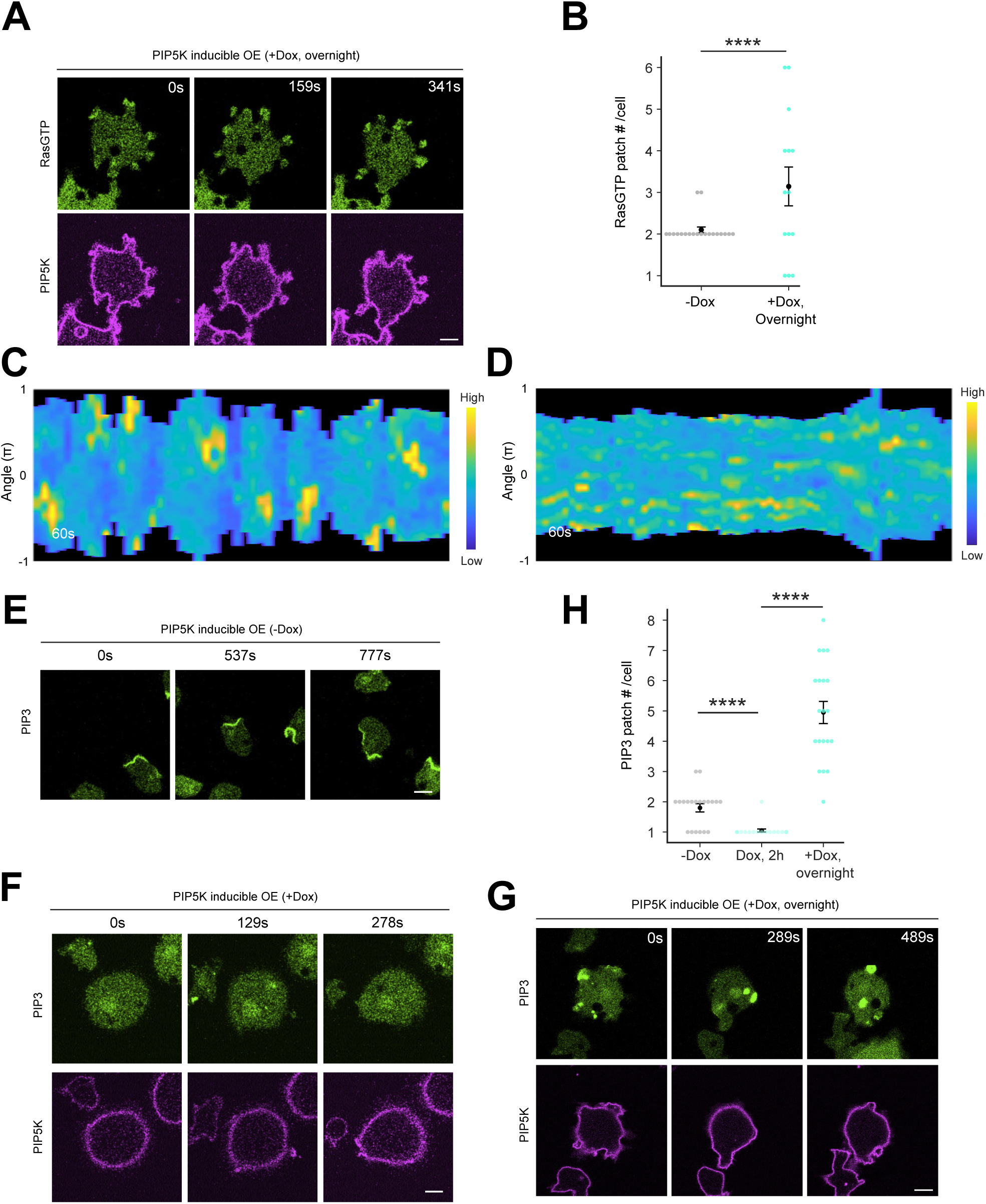
Inducible PIP5K overexpression inhibits Ras and PI3K signaling activities. **(A)** Live-cell images of *Dictyostelium* cells co-expressing RBD-GFP and Dox-inducible PIP5K-mCherry with overnight Dox induction, showing multiple small RasGTP patches (top panels) in “Spiky” PIP5K OE cells (bottom panels). **(B)** Quantification of RasGTP patch number of *Dictyostelium* cells expressing Dox-inducible PIP5K-GFP, either without or with overnight Dox treatment. In case of with Dox treatment, onlyspiky population has been included and round population (showing no patches) has been excluded. Here, n_c_=20 for -Dox population and n_c_=14 for +Dox population; data from N=3 independent experiments; p value by the Mann-Whitney U test. **(C, D)** Representative 360° membrane kymograph of RBD-GFP intensity in -Dox population (C) and +Dox population (D), respectively, corresponding to Figure 2J and Figure S15A, respectively. **(E-G)** Live-cell images of *Dictyostelium* cells co-expressing PH_crac_-YFP and Dox-inducible mRFP-PIP5K without Dox induction (E) or with overnight Dox induction (F-G), showing PIP3 activities nearly completely vanished in rounded PIP5K OE cells (F), or appeared as multiple small patches in spiky cells (G). **(H)** Quantification of PIP3 patch number of *Dictyostelium* cells expressing Dox-inducible PIP5K-GFP, either without or with Dox treatment for different amount of time periods. For overnight Dox treatment case, only spiky populations are included and round population (showing no patches) has been excluded. Here, n_c_=20 for all three populations; data from N=3 independent experiments; p value by the Mann-Whitney U test.

**Figure S16.**
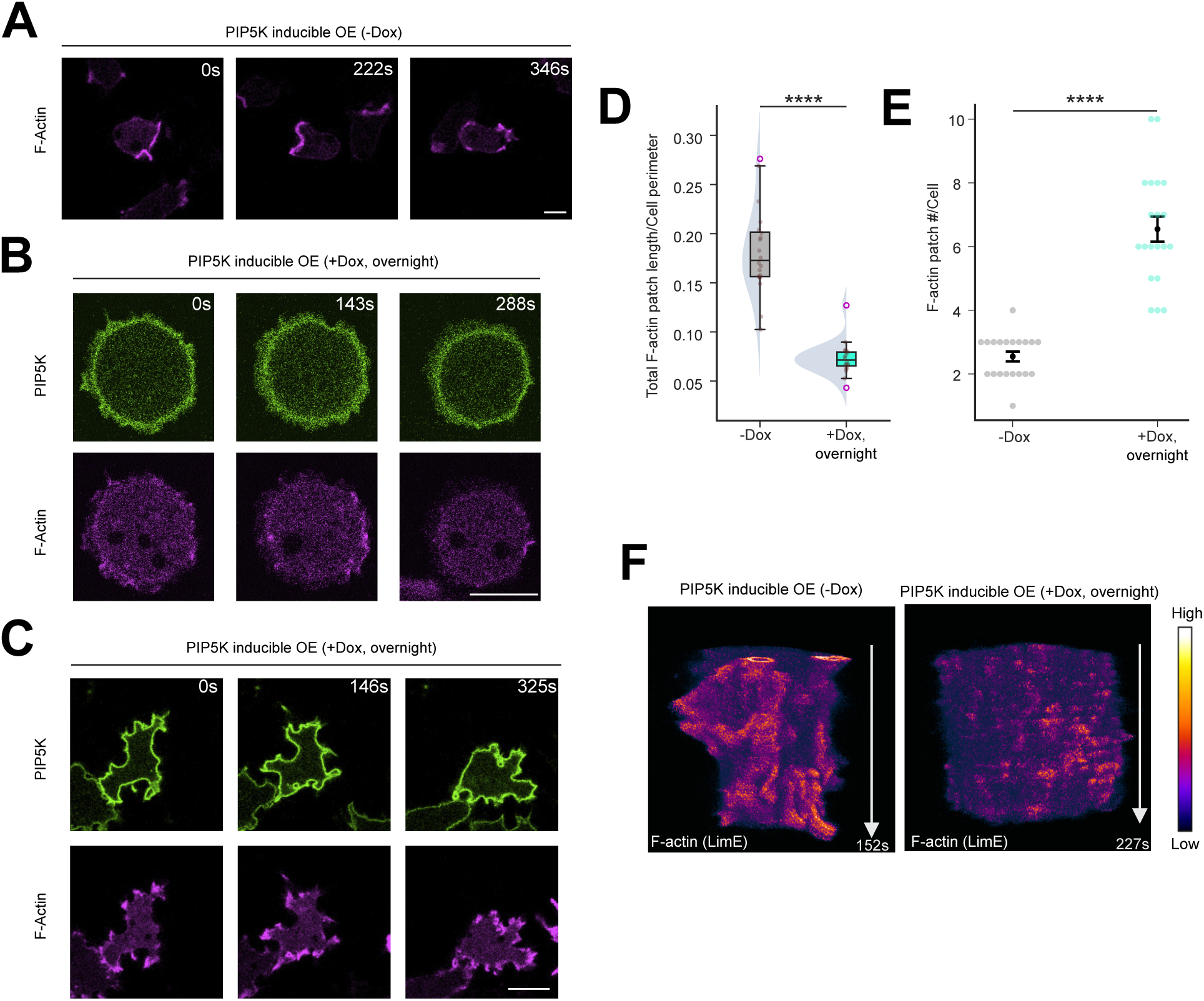
Inducible PIP5K overexpression impairs F-actin polymerization. **(A-C)** Live-cell images of *Dictyostelium* cells co-expressing LimE-mCherry and Dox-inducible GFP-PIP5K without Dox induction (A) or with overnight Dox induction (B-C), showing F-actin polymerization was nearly completely abolished in rounded PIP5K OE cells (B), or appeared as multiple small actin patches in spiky cells (C). **(D-E)** Quantification of newly polymerized F-actin level on the plasma membrane (D), or actin patch number in *Dictyostelium* cells, either without or with overnight Dox treatment, demonstrating that F-actin patches are reduced dramatically upon overnight Dox induction. The levels are quantified in terms the total actin patch length per unit cell perimeter length. In (E), for overnight Dox treatment case, only spiky populations are included and round population (showing no patches) has been excluded. Here, n_c_=20 for all populations; data from N=3 independent experiments; p value by the Mann-Whitney U test. **(F)** Two t-stacks from cells co-expressing LimE-mCherry and Dox-inducible GFP-PIP5K without Dox induction (left) or with overnight Dox induction (right), showing F-actin patches are much smaller and transient in PIP5K OE cells. The white arrow corresponds to the time duration of the t-stack.

**Figure S17.**
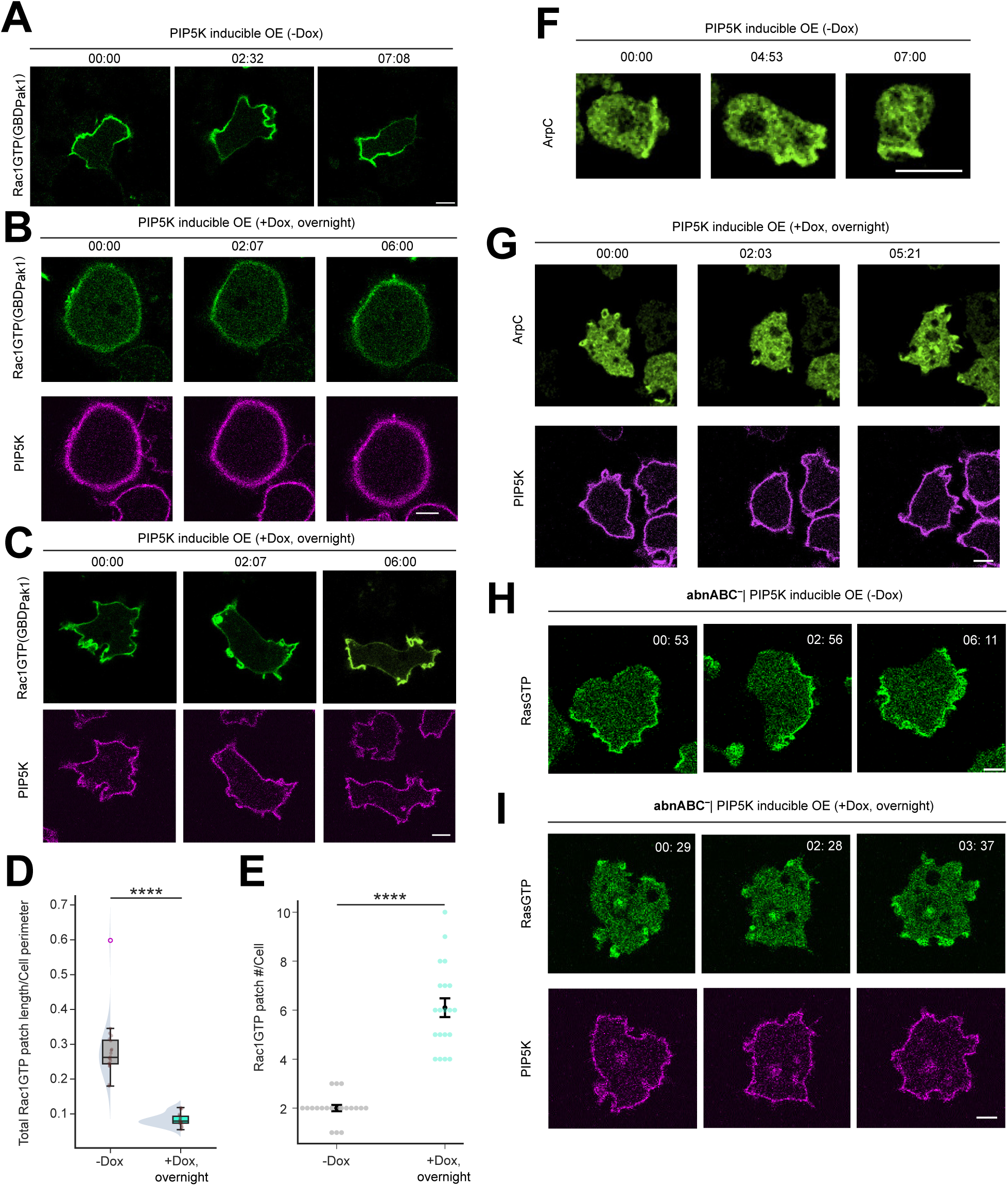
Inducible PIP5K overexpression impairs Rac1/Arp2/3 complex/F-actin axis. **(A-C)** Representative live-cell images of *Dictyostelium* cells co-expressing GBD_Pak1_-GFP (biosensor for Rac1) and Dox-inducible mRFP-PIP5K, without DOX induction (A) or with overnight Dox induction (B-C), showing Rac1 patches nearly vanished in rounded PIP5K OE cells (B), or appeared as multiple small Rac1 patches in spiky cells (C). **(D, E)** Quantification of overall Rac1GTP level on the cell membrane (D), or Rac1 patch number in *Dictyostelium* cells, either without or with overnight Dox treatment, showing Rac1 patches are reduced dramatically upon overnight Dox induction. The levels are quantified in terms the total Rac1 patch length per unit cell perimeter length. In (E), for overnight Dox treatment case, only spiky populations are included and round population (showing no patches) has been excluded. Here, n_c_=20 for all populations; data from N=3 independent experiments; p value by the Mann-Whitney U test. **(F, G)** Live-cell images of *Dictyostelium* cells co-expressing ArpC-GFP and Dox-inducible mRFP-PIP5K without Dox induction (F) or with overnight Dox induction (G), showing ArpC activities are impaired strongly in PIP5K OE cells. **(H, I)** Live-cell images of *Dictyostelium abnABC-* cells co-expressing RBD-GFP and Dox-inducible mRFP-PIP5K without Dox induction (H) or with overnight Dox induction (I), showing RasGTP activities are still ilow in PIP5K OE cells, even when positive feedback from branched actin is enhanced.

**Figure S18.**
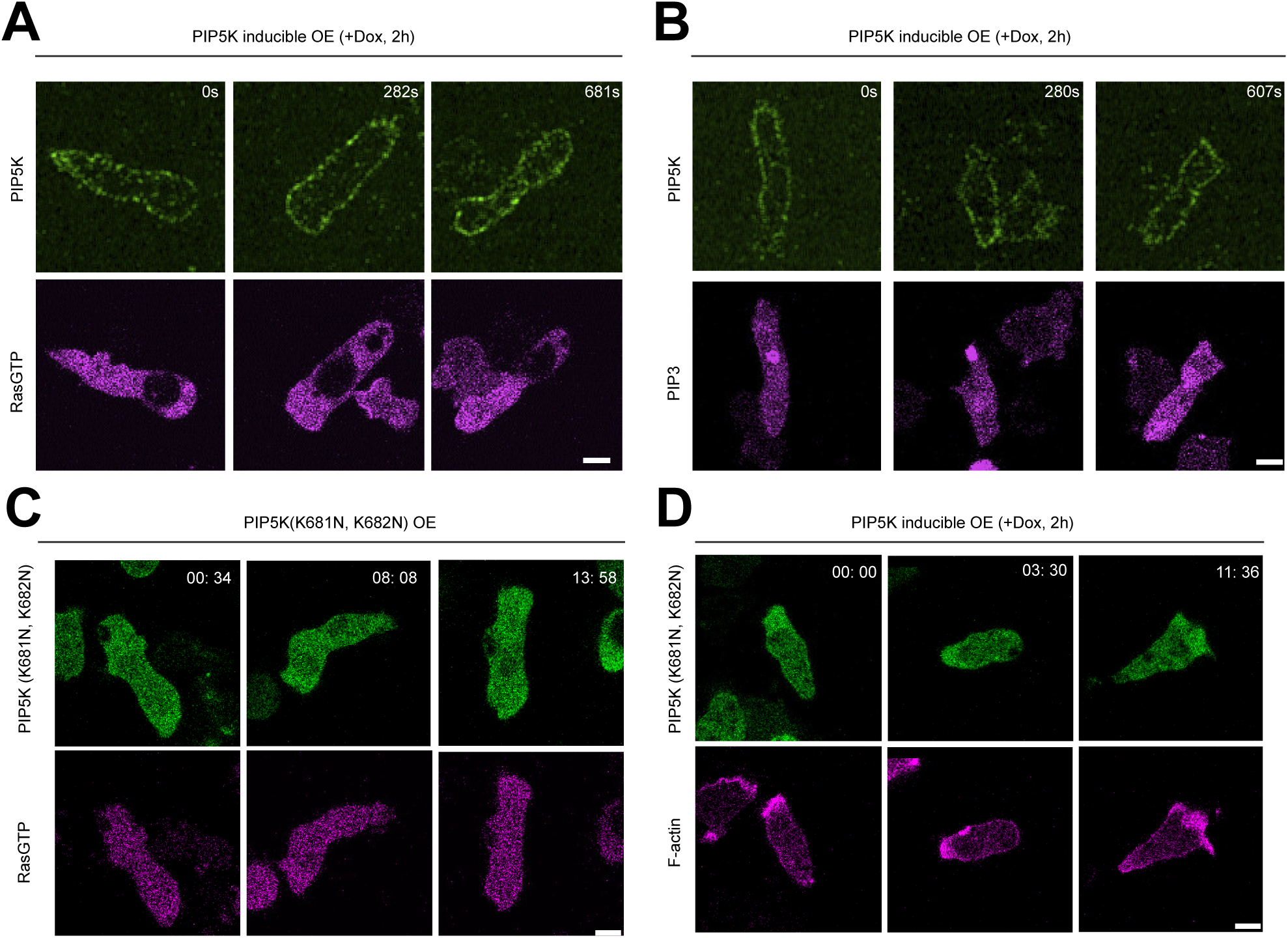
Moderate overexpression of PIP5K polarizes cells but inhibits Ras/PI3K/F-actin signaling activities. **(A, B)** Representative live-cell images of *Dictyostelium* cells co-expressing Dox-inducible PIP5K-GFP with 2h Dox induction, along with RBD-GFP (A), or PHcrac-YFP (B), showing impaired RasGTP and PIP3 activities. **(C, D)** Representative live-cell images of *Dictyostelium* cells co-expressing PIP5K(K681N, K682N)-GFP mutant with RBD-mRFP (C), or LimE-mCherry (D), showing impaired PIP3 activities and spatially confined F-actin polymerization.

**Figure S19.**
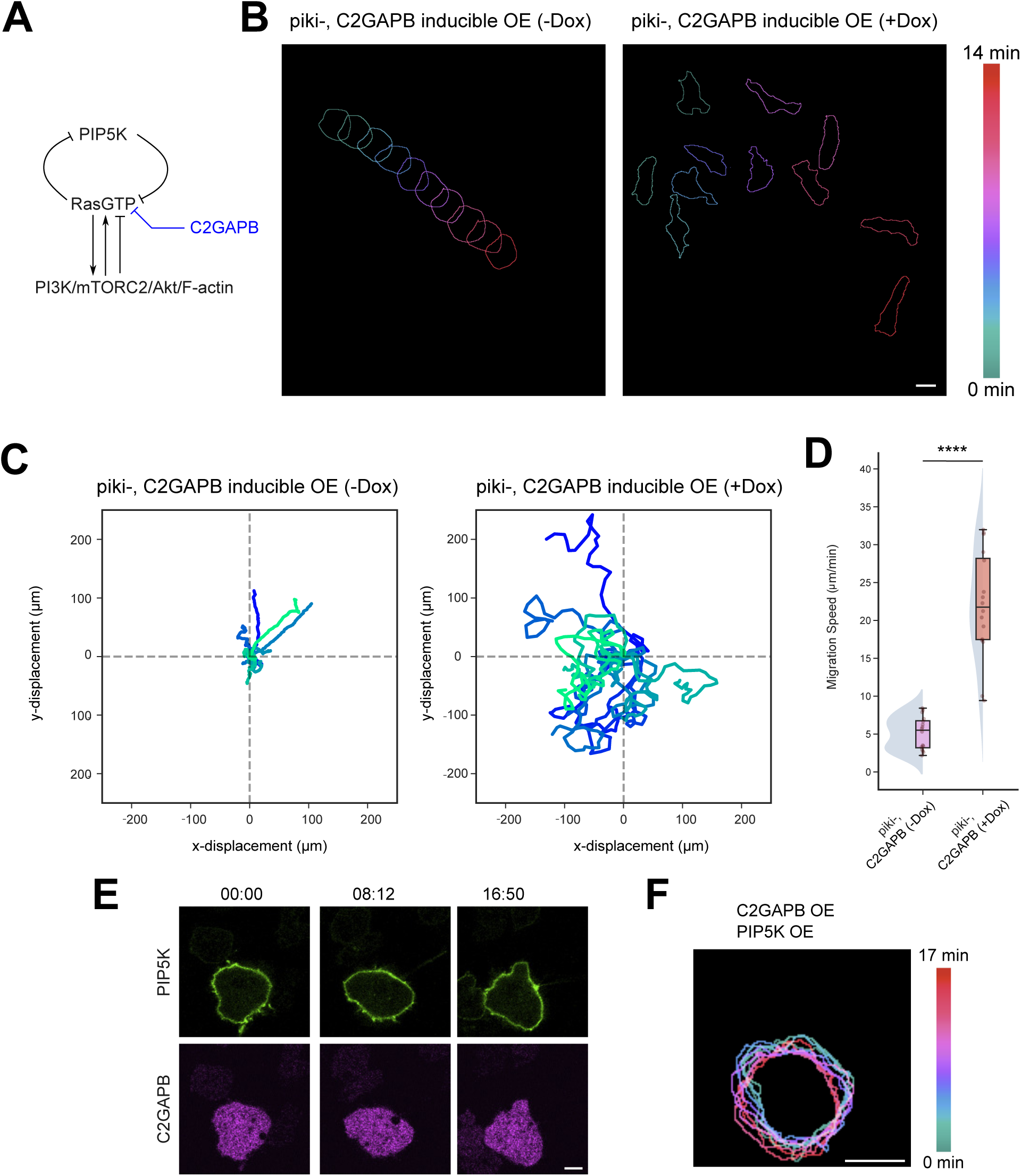
Attenuating Ras activity with C2GAPB partially restores cell polarity in *pikI-* cells. **(A)** Schematic of signal transduction network showing two pathways inhibiting RasGTP. Note that, C2GAPB is a RasGAP, which is known to inhibit Ras activities and polarizes cells upon overexpression. **(B)** Color-coded temporal overlay profiles of migrating *piki- Dictyostelium* cells expressing doxycycline-inducible mRFP-C2GAPB without Dox induction (left) or with overnight Dox induction (right), showing that attenuation of Ras activity with C2GAPB can restore polarity in *piki-* cells. **(C)** Centroid tracks of randomly migrating *Dictyostelium piki-* cells (n_c_=16 for -Dox populations, n_c_=15 for +Dox populations) expressing Dox-inducible C2GAPB, without (left) or with overnight (right) Dox treatment. Each cell was tracked for 18 minutes and was reset to the same origin. **(D)** Quantification of the migration speed of non-Dox treated and overnight Dox-treated cells corresponding to Figure S19C, showing inducible overexpression of C2GAPB significantly speeds up pikI- cells. Here n_c_=16 for either -Dox population or +Dox population; p value by the Mann-Whitney U test. **(E, F)** Representative live-cell images (E) and color-coded temporal overlay (F) of *Dictyostelium* cells overexpressing both PIP5K-GFP and mRFP-C2GAPB. In (E), time is in mm:ss format.

**Figure S20.**
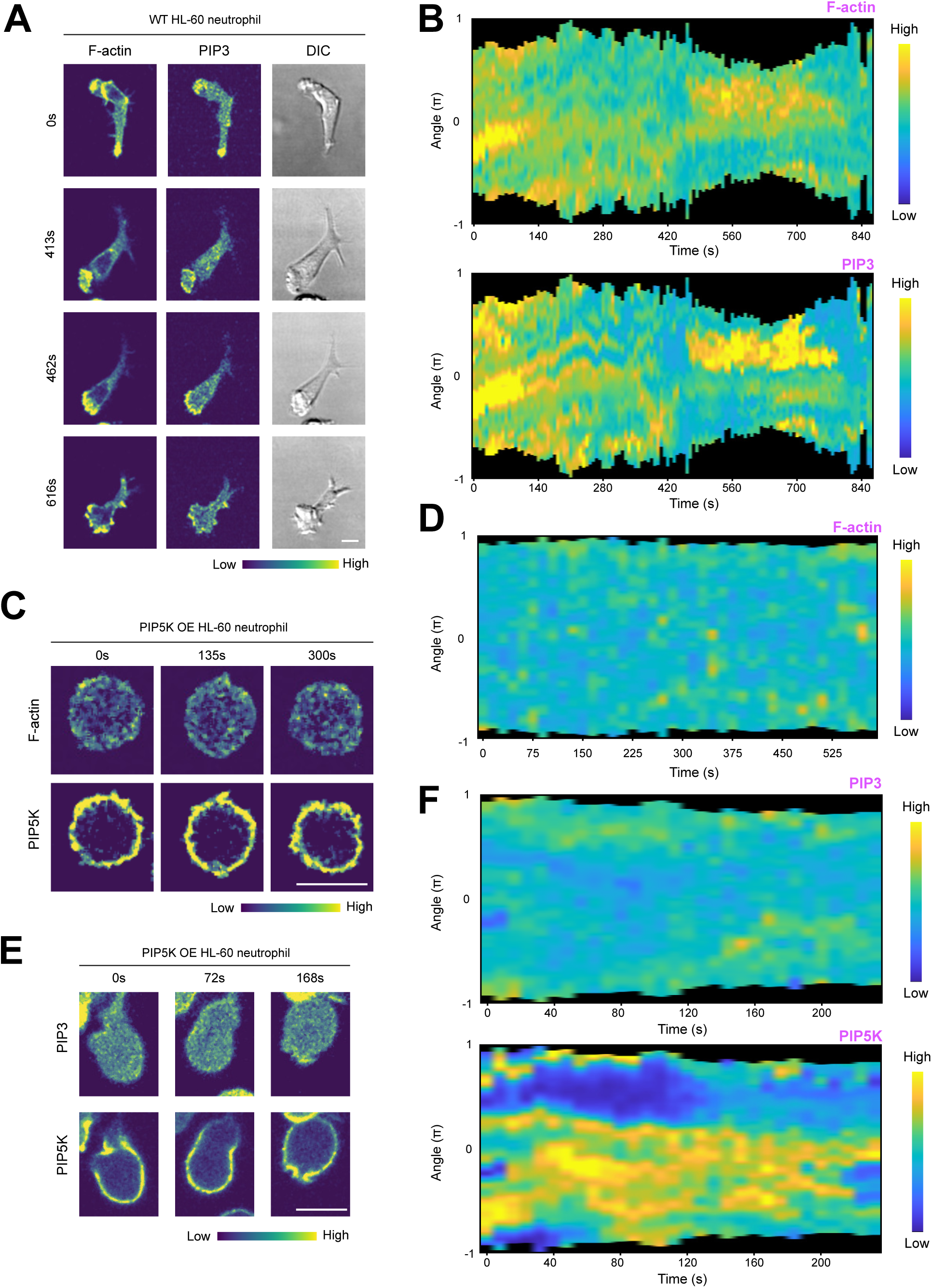
Overexpressing PIP5K in HL-60 neutrophils inhibits PIP3 signaling activities and F-actin polymerization. **(A, C, E)** Live-cell images of differentiated HL-60 neutrophils cells co-expressing LifeAct- mCherry and PH_Akt_-GFP (A), or cells co-expressing LifeAct-iRFP703 and PIP5K1C-GFP, or cells co-expressing PH_Akt_-RFP and PIP5K1C-GFP. Note that overexpression of PIP5Ks in HL-60 neutrophils strongly inhibited PIP3 signaling and F-actin polymerization, and thereby impaired cell polarity and protrusion formation. **(B, D, F)** Representative 360° membrane kymographs of indicated components; these correspond to cells shown in (A, C, E), respectively. Note that, unlike WT HL-60 neutrophils, in PIP5K1C overexpression population, both PIP3 activities and F-actin polymerization levels remained consistently low.

**Figure S21.**
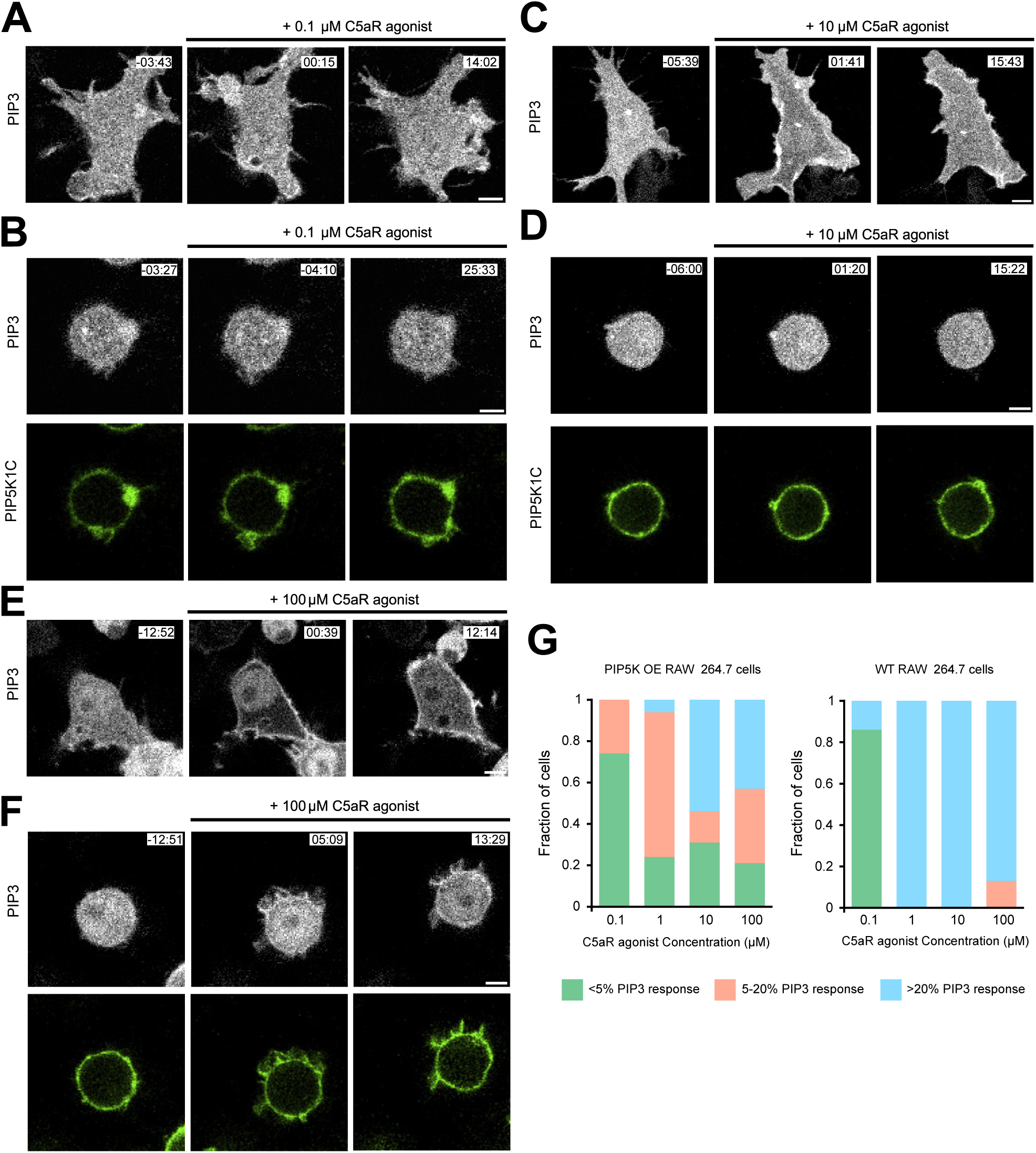
Overexpressing PIP5Ks in RAW 264.7 macrophages can resist receptor input activation of downstream signaling activities. **(A-F)** Representative live-cell time-lapse images of RAW 264.7 macrophage cells, wild type (A, C, E) or overexpressing PIP5K1C-GFP (B, D, F) where they were globally stimulated with 0.1 μM (A-B), or 10 μM (C-D), or 100 μM (E-F) of C5aR agonist, showing cells with high PIP5K expression levels require higher concentration chemoattractant to activate PIP3 signaling activities. Cells were expressing PH_Akt_-mCherry whose cytosol to membrane translocation was monitored over time. Time in min:sec format and C5aR agonist was added at time t=0. **(G)** Quantification of cell fractions for different categories: <5% PIP3 response (green column), 5-20% PIP3 response (salmon column), and >20% PIP3 response (blue column), between PIP5K1C OE and WT cell populations, showing at all C5aR concentrations, PIP5K1C OE cells have exhibited substantially subdued PIP3 response.

**Figure S22.**
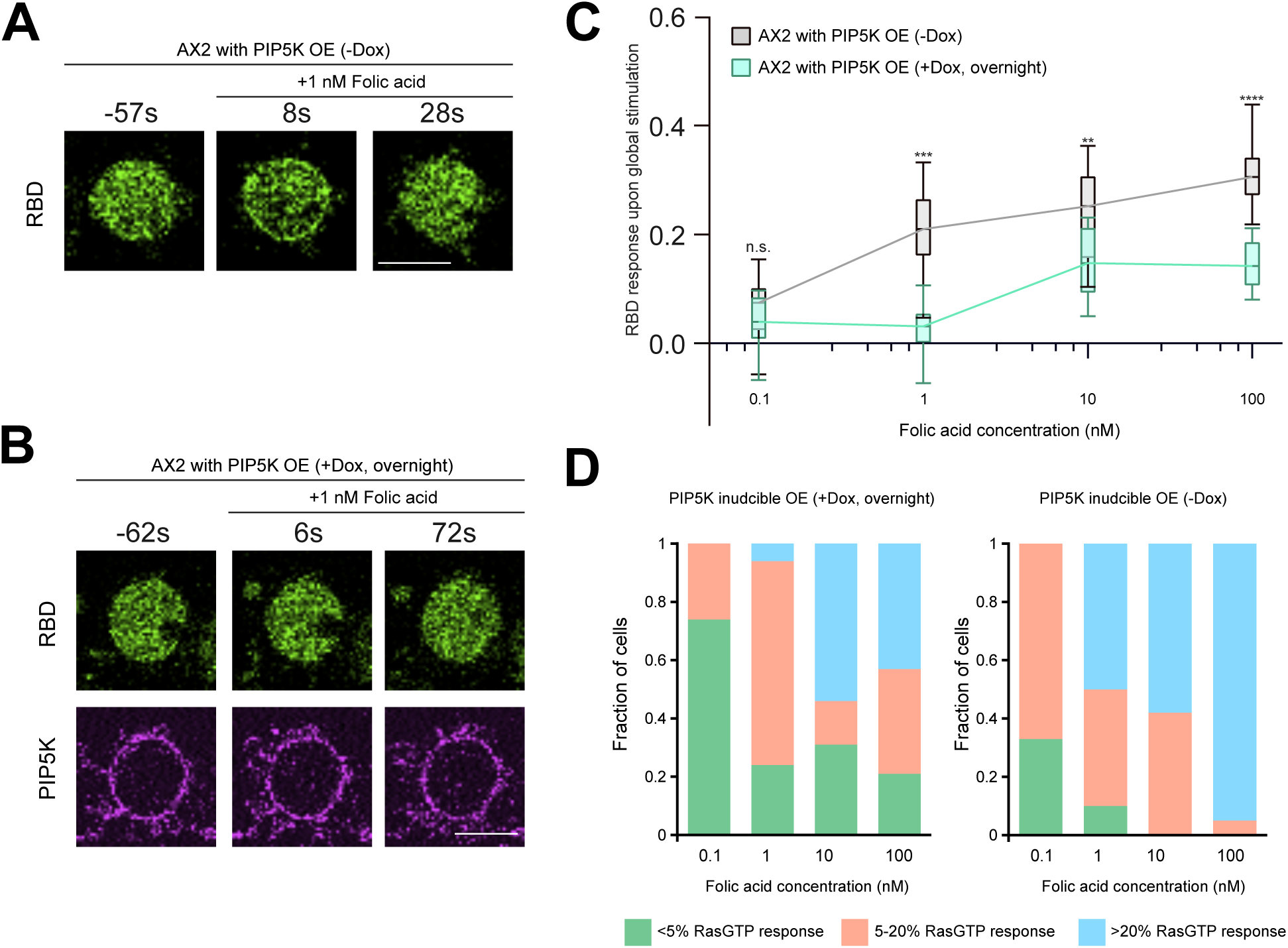
Overexpressing PIP5K in *Dictyostelium* can subvert receptor input driven activation of downstream signaling activities. **(A, B)** Representative live-cell time-lapse images of *Dictyostelium* cells, expressing doxycycline-inducible mRFP-PIP5K without Dox induction (A) or with overnight Dox induction (B), where they are responding to global simulation with 1 nM of folic acid. Cells were expressing RBD-GFP, whose cytosol-to-membrane translocation was monitored over time. Time in min:sec format and C5aR agonist was added at time t=0. **(C)** Normalized responses shown by WT (grey) and PIP5K-GFP overexpressing (mint green) *Dictyostelium* cells to different doses of folic acid, demonstrating impeded receptor activation-driven signaling activation in PIP5K OE cells. Responses were computed in terms of (1 - decrease in cytosolic intensity of RBD-GFP). Here, n_c_≥12 for -Dox populations and n_c_≥10 for -Dox populations for each concentration; for either population, from N≥3 independent experiments. The p-values are computed by the Mann-Whitney U test. **(D)** Quantification of cell fractions for different categories: <5% RasGTP response (green column), 5-20% RasGTP response (salmon column), and >20% RasGTP response (blue column), between -Dox and +Dox populations, showing at all folic acid concentrations, PIP5K OE cells exhibited substantially subdued RasGTP response.

**Figure S23.**
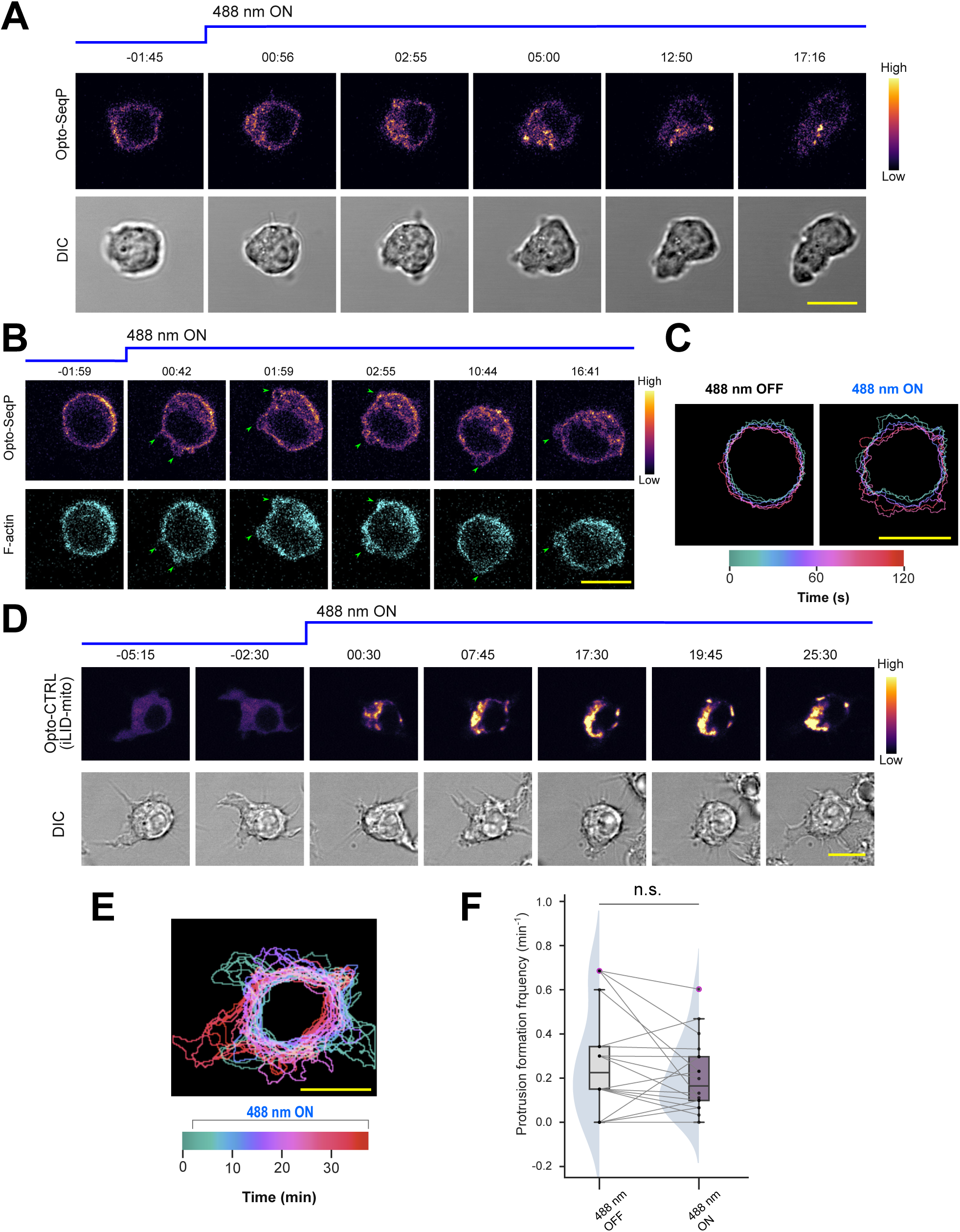
Optogenetic recruitment of plasma membrane PIP5K to mitochondria elicits symmetry breaking and triggers protrusion formation, whereas control recruitment has no effect. **(A, B)** Representative time-lapse images of RAW 264.7 macrophages expressing Opto-SeqP, before and after 488 nm illumination driven recruitment of PIP5K1B from plasma membrane to mitochondrial outer membrane (488 nm was turned ON at t=0), demonstrating that cells generally formed more protrusions (A, B) and often assumed a polarized morphology (A), upon depletion of PIP5K from the plasma membrane. Time in min:sec format. **(C)** Color-coded temporal overlay profiles of RAW 264.7 macrophages before and after Opto-SeqP recruitment, showing that substantially more protrusion formed as plasma membrane PIP5K is recruited to mitochondria. **(D-F)** Representative live-cell time-lapse images, color-coded temporal overlay, and quantification of protrusion formation frequency in RAW 264.7 macrophage cells, where Opto-CTRL (mCrismon-NES-SspB) was recruited to mitochondrial outer membrane by anchor iLID-mito, upon 488 nm laser illumination. Note that, expression of Opto-CTRL has not stopped protrusion formation and no significant increase in protrusion frequency was observed as Opto-CTRL was sequestered in mitochondria. In (D), time in min:sec format. In (F), for each of the n_c_ = 18 cells, protrusion formation events were tracked, before and after the recruitment. The p values are computed by the Wilcoxon matched-pairs signed-rank test. For pairwise comparison, data from the same cell are connected by grey lines.

**Figure S24.**
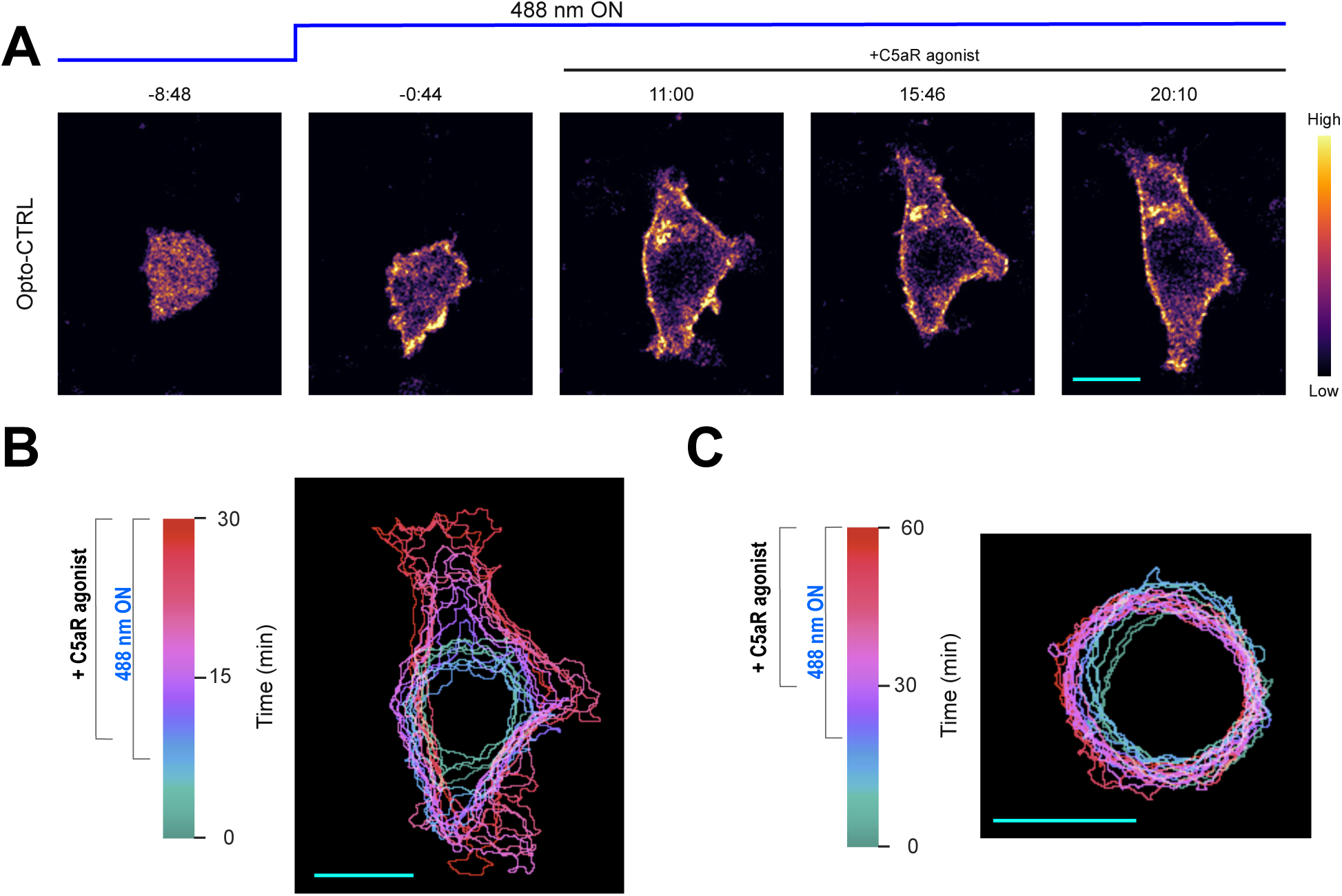
Opto-CTRL recruitment from cytosol to plasma membrane did not inhibit protrusion formation induced by C5aR agonist stimulation in RAW 264.7 cells. **(A)** Representative live-cell time-lapse images of RAW 264.7 macrophages undergoing blue light-triggered global recruitment of Opto-CTRL to plasma membrane, followed by global stimulation with C5aR agonist, demonstrating that cells generated new protrusions and spread substantially upon receptor input. Cells were expressing Opto-CTRL (mCrismon-NES-SspB) with iLID-CAAX, Time in min:sec format. **(B, C)** Color-coded temporal overlay profiles of RAW 264.7 macrophages before and after optogenetic recruitment and C5aR agonist stimulation. In (B) Opto-CTRL was used, and in (C), Opto-RecP was used.

**Figure S25.**
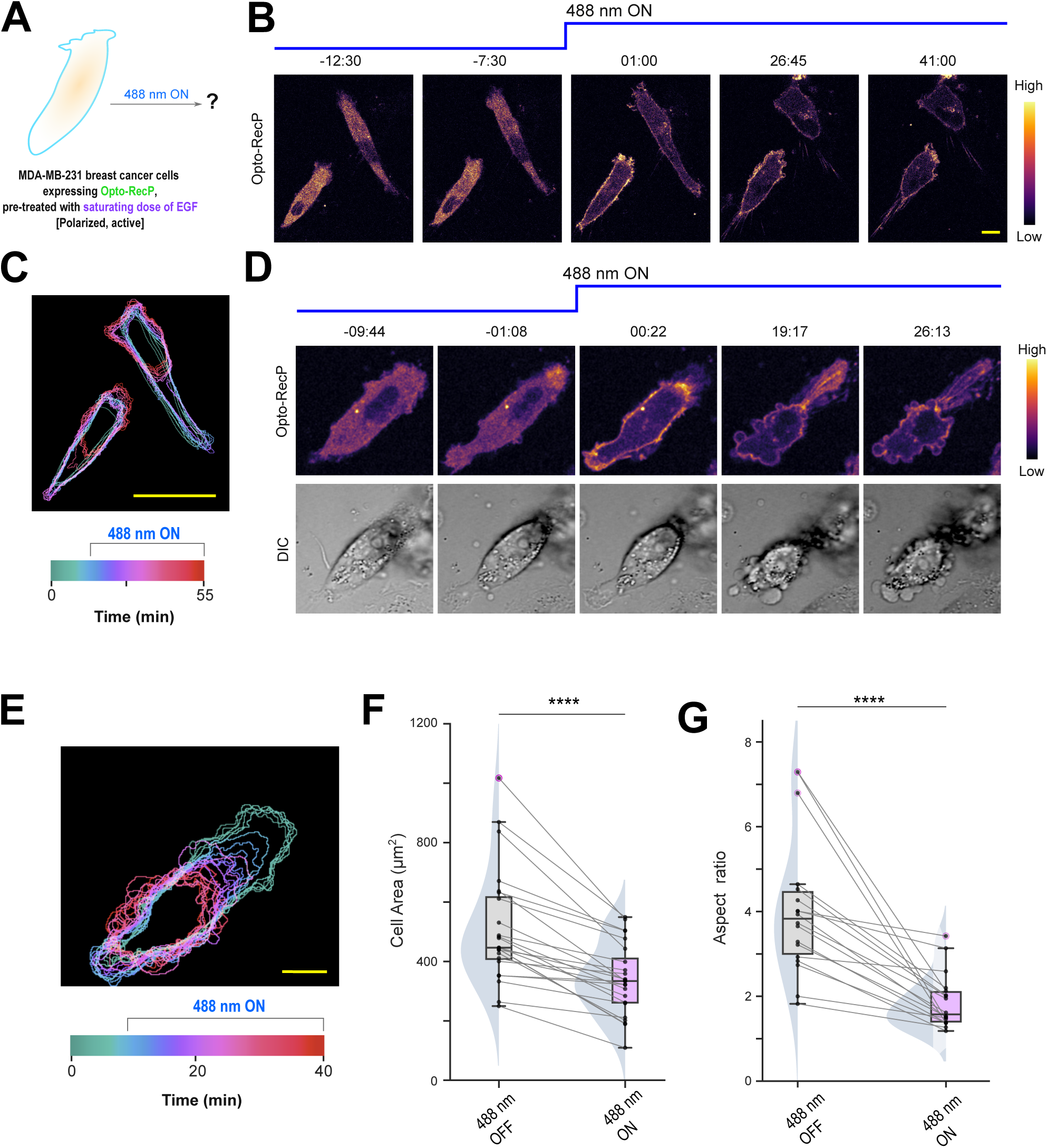
Rapid optogenetic increase of PIP5K level on the plasma membrane induces cell contraction in MDA-MB-231 cancer cells. **(A)** Illustration of the Opto-RecP recruitment system in MDA-MB-231 cells. Note that, cells were pretreated with a saturating dose of EGF. **(B, D)** Two representative live-cell time-lapse image series of MDA-MB-231 cells, before and after Opto-RecP recruitment from cytosol to membrane, demonstrating that cells contracted and blebs were induced upon recruitment. Time in min:sec format. **(C, E)** Color-coded temporal overlay profiles of MDA-MB-231 cells before and after Opto-RecP recruitment (corresponding to (B) and (D) respectively), showing that acutely increasing PIP5K on the plasma membrane substantially induces cell retraction, even when EGF was is present in saturating dose. **(F, G)** Quantifications of cell area (F) and aspect ratio (G) of MDA-MB-231 cells, before and after recruiting cytosolic PIP5K1C to the plasma membrane, showing that acutely increasing PIP5K on the membrane induces cell retraction and cells abrogate their elongated, polarized shape. Here, n_c_=24 for (F) and n_c_= 18 for (G); for both populations; data from N=3 independent experiments; p value by the Mann-Whitney U test.

**Figure S26.**
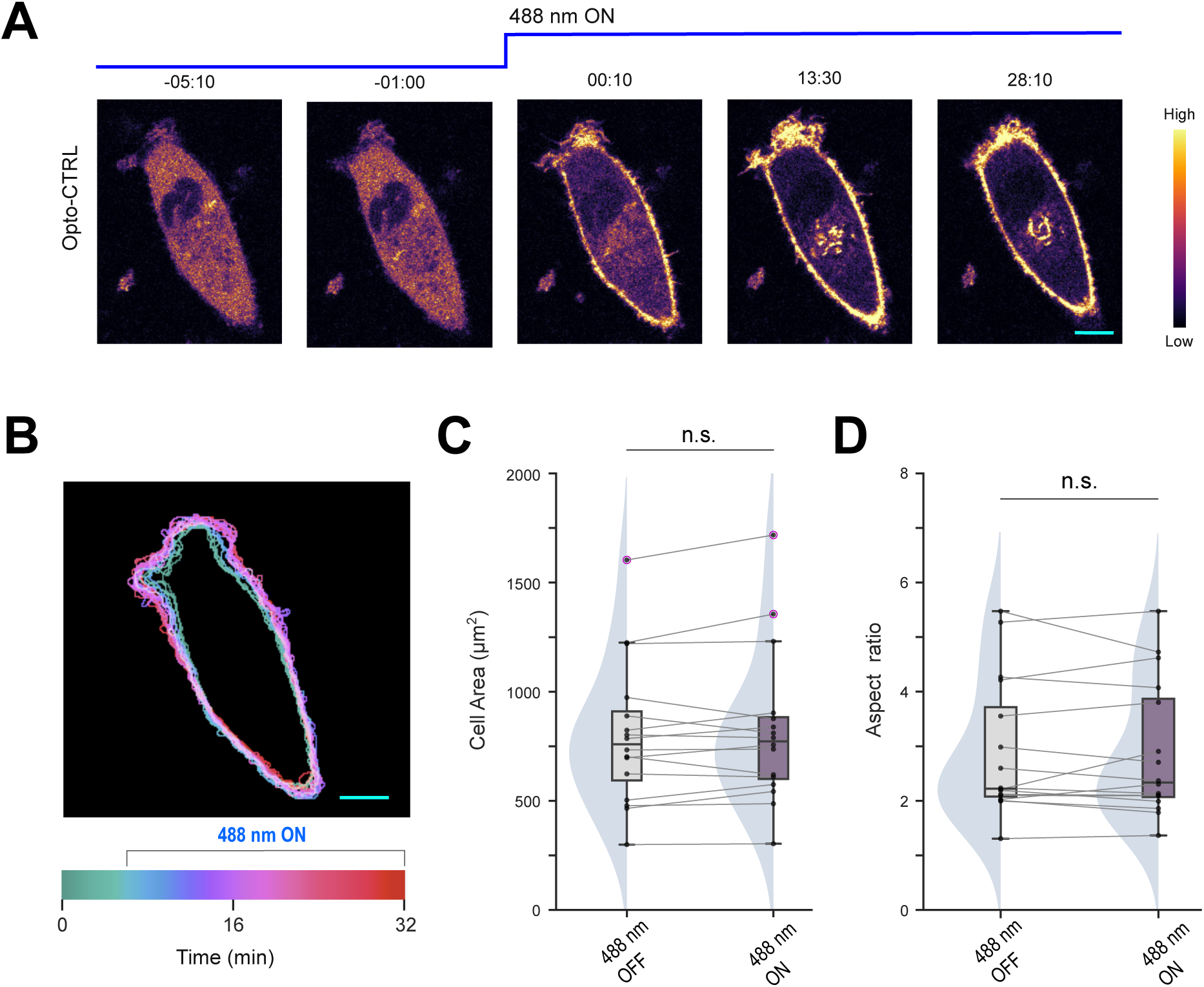
Opto-CTRL recruitment from cytosol to plasma membrane did not induce cell contraction in MDA-MB-231 cancer cells. **(A, B)** Representative live-cell time-lapse image (A) and color-coded of temporal overlay (B) of MDA-MB-231 cells, before and after Opto-CTRL recruitment from cytosol to membrane, demonstrating that cell morphology underwent negligible changes. Cells were expressing Opto-CTRL (mCrismon-NES-SspB) with iLID-CAAX, Time in min:sec format. **(C, D)** Quantifications of cell area (C) and aspect ratio (D) of MDA-MB-231 cells, before and after recruiting Opto-CTRL from cytosol to the plasma membrane, revealing no significant phenotypic transitions upon recruitment. Here, n_c_=16 for (C) and (D), for both populations; p-value by the Mann-Whitney U test.

**Figure S27.**
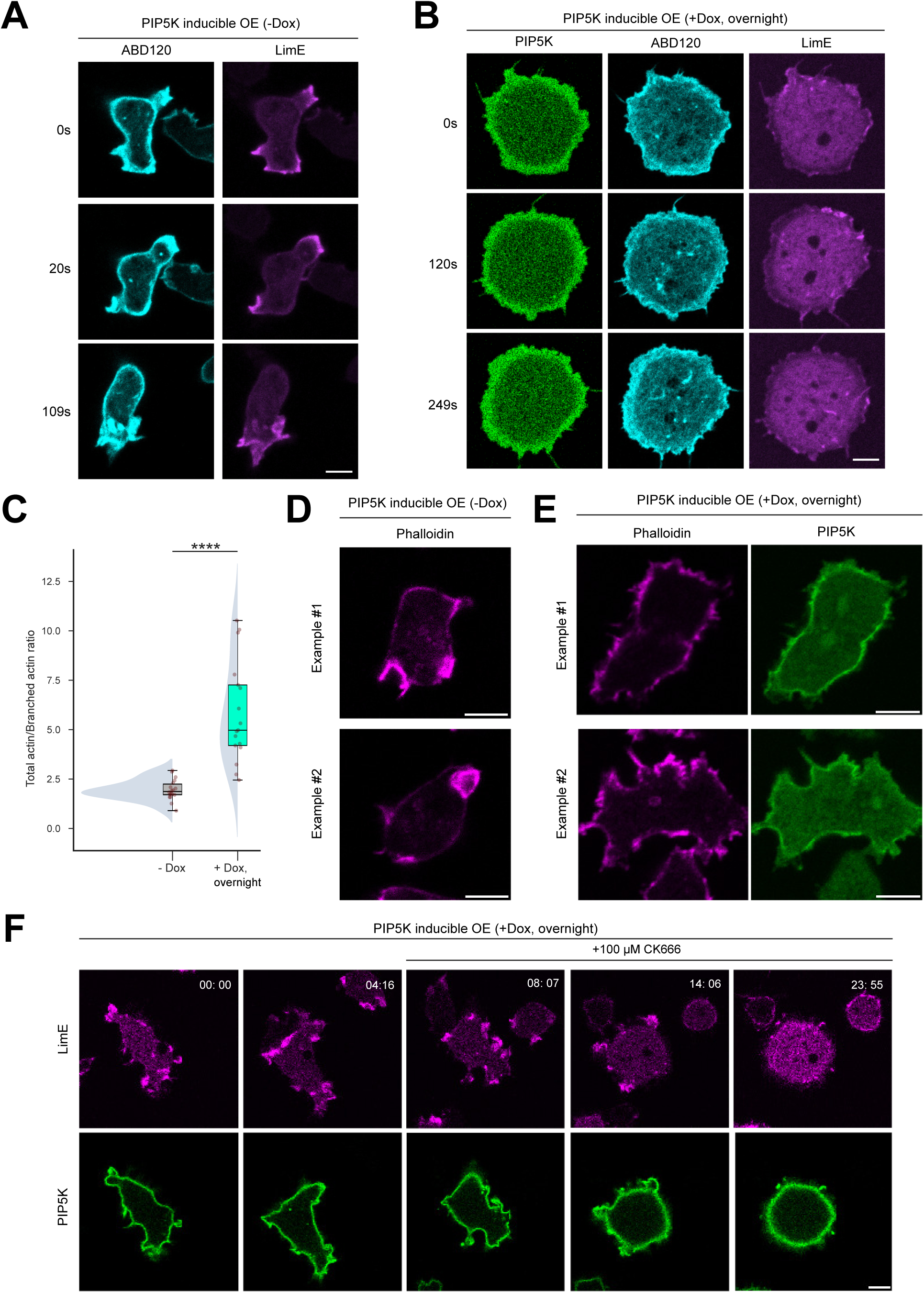
Inducible PIP5K overexpression increases linear actin assembly. **(A, B)** Live-cell images of *Dictyostelium* cells co-expressing ABD120-GFP, LimE-Halo (TMR conjugated), and Dox-inducible mRFP-PIP5K without Dox induction (A) or with overnight Dox induction (B). **(C)** Quantification of the total actin patch to branched actin patch ratio, showing that cells shift the equilibrium to more linear actin when PIP5K is overexpressed. The levels are quantified in terms of the total ABD120 patch length divided by the total branched actin patch (LimE) in the same cell. Here, n_c_=18 for -Dox populations, and n_c_=17 for +Dox populations; data from N=3 independent experiments; p value by the Mann-Whitney U test. **(D, E)** Image of *Dictyostelium* cells expressing Dox-inducible GFP-PIP5K without Dox induction (D) or with overnight Dox induction (E); cells were fixed and stained with Phalloidin, showing more total actin in PIP5K OE cells. **(F)** Representative live-cell time-lapse images of a spiky *Dictyostelium* cells co-expressing LimE-mCherry and Dox-inducible PIP5K-GFP (with overnight DOX induction), before and after 100 μM Arp2/3 inhibitor CK666 treatment, showing spiky cells can shift to rounded cells when branched actin is further inhibited.

**Figure S28.**
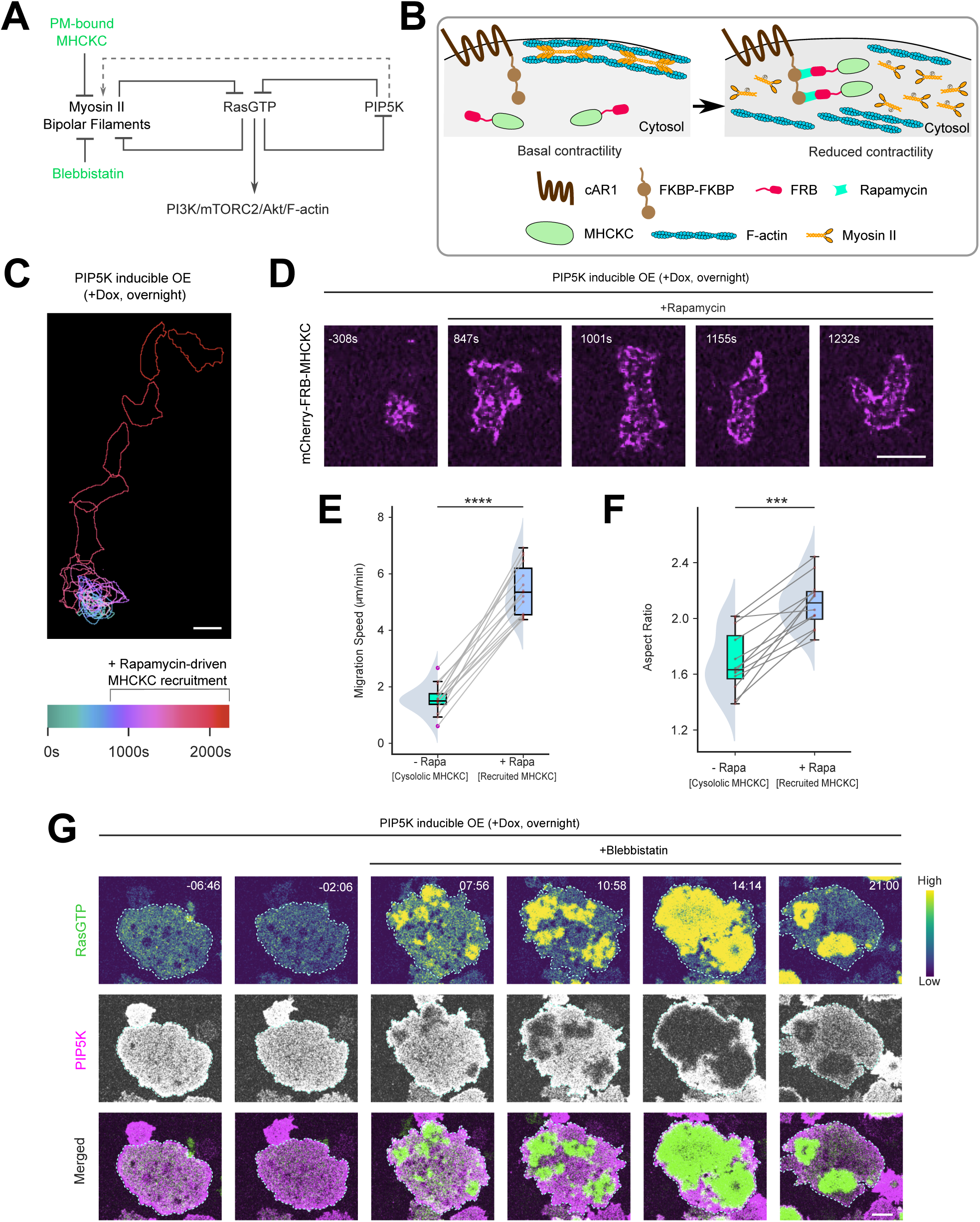
Inhibiting actomyosin contractility counteracts PIP5K overexpression-induced phenotypes. **(A)** Schematic of perturbations used to inhibit myosin II activities, in context of the signal transduction network. **(B)** Illustration of chemically inducible dimerization recruitment system to increase of myosin heavy chain kinase C (mCherry-FRB-MHCKC) to plasma membrane with cAR1-FKBP-FKBP as the membrane anchor. Upon recruitment to membrane, MHCKC phosphorylates myosin heavy chains and causes disassembly of myosin bipolar thick filaments from the membrane-cortex. **(C, D)** Color-coded temporal overlay profiles (C) and live-cell images (D) of *Dictyostelium* cells co-expressing doxycycline-inducible GFP-PIP5K (with overnight Dox induction) and recruitable MHCKC system (cAR1-FKBP-FKBP and mCherry-FRB-MHCKC; as shown in (B)). Note that, upon MHCKC recruitment, cells switched from round, immobilized state to polarized shape and migrated much faster. In (D), Rapamycin was added at t=0 to induce the recruitment. **(E, F)** Quantifications of migration speed (E) and aspect ratio (F) of *Dictyostelium* cells co-expressing doxycycline-inducible GFP-PIP5K (with overnight Dox induction) and recruitable MHCKC system (cAR1-FKBP-FKBP and mCherry-FRB-MHCKC), before (- Rapa) or after recruitment (+Rapa), showing that reduction in myosin II-based contractility can counteract PIP5K overexpression-driven cellular phenotypes. Here n_c_=15 for (E) and n_c_=12 for (F), for either -Rapa population or +Rapa population; For pairwise comparison between before and after recruitment, data from the same cell are connected by gray lines; p value by the Mann-Whitney U test. **(G)** Representative live-cell images of *Dictyostelium* cells co-expressing RBD-GFP and Dox-inducible mRFP-PIP5K (with overnight Dox induction), before or after 50 μM myosin II inhibitor blebbistatin addition, showing that inhibition of myosin II dramatically activates Ras signaling in PIP5K OE cells and induces the appearance and propagation of large ventral waves. Time in min:sec format.

**Figure S29.**
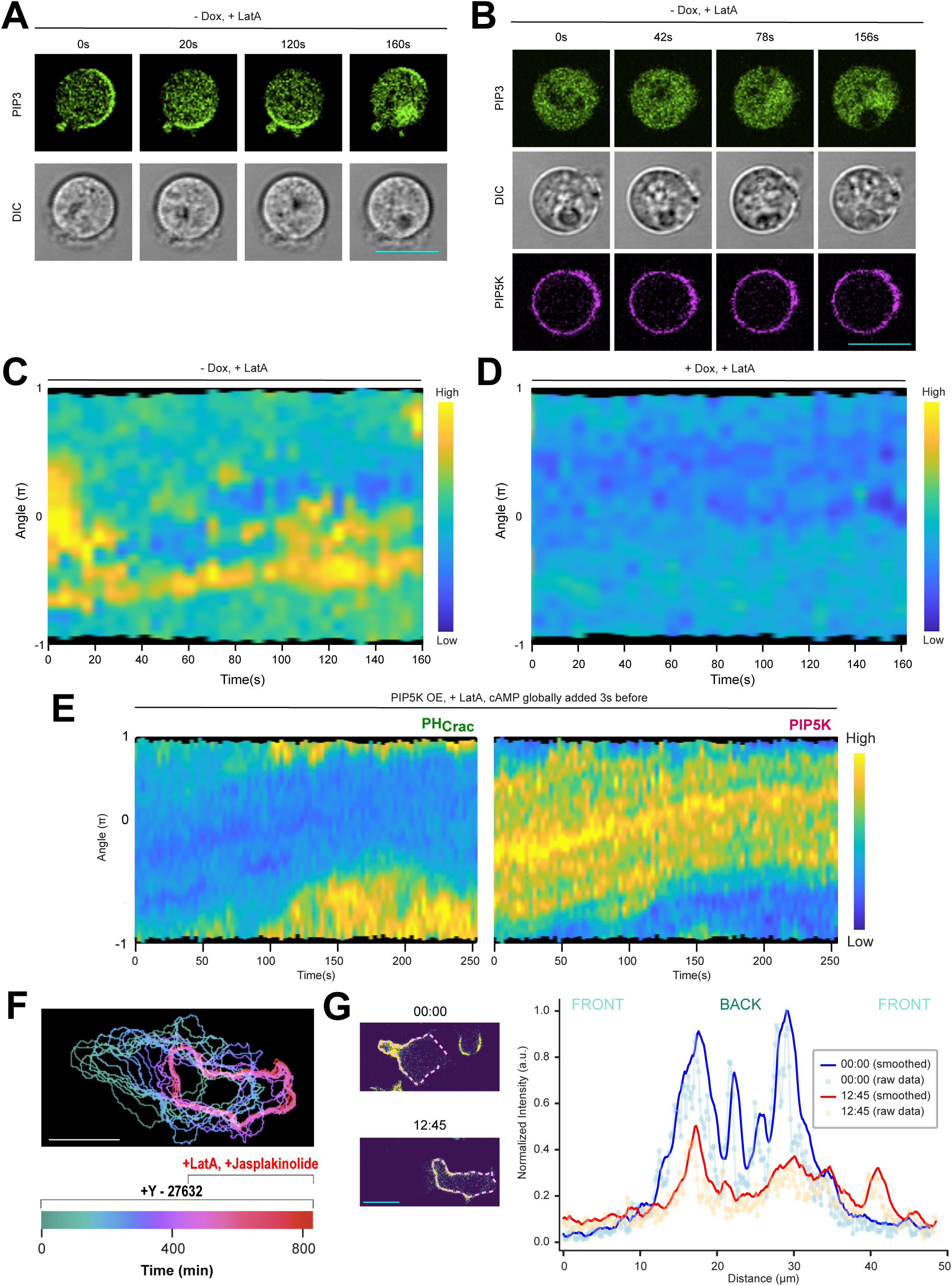
PIP5K can inhibit signaling activities and maintain back localization independent of cytoskeletal dynamics. **(A, B)** Live-cell images of *Dictyostelium* cells co-expressing PHcrac-YFP and Dox-inducible GFP-PIP5K without Dox induction (A) or with overnight Dox induction (B). Cells were pre-treated with Latrunculin A (final concentration 5µM) and caffeine (final concentration 4mM) for 20min. The PIP3 activities nearly vanished in PIP5K OE cells (B), compared to Dox-untreated counterpart (A). **(C, D)** Representative 360° membrane kymographs of PH_crac_-YFP, corresponding to cells shown in (A) and (B), respectively. Note that, PIP3 activities nearly vanished in PIP5K OE cells (B, D), compared to Dox-untreated counterpart (A, C). **(E)** Representative 360° membrane kymograph of PHcrac-YFP and PIP5K-GFP, respectively, corresponding to Figure 4J. **(F)** Color-coded temporal overlay profiles of differentiated WT HL-60 neutrophils expressing GFP-PIP5K, before and after JLY cocktail treatment, corresponding to Figure 4K-L, showing that while the cell froze and stopped migrating, it maintained polarized state upon JLY treatment. **(G)** Line-scan intensity profile of GFP-PIP5K, along with the dotted magenta line on the membrane, showing that although upon JLY treatment, the intensity profile of PIP5K diffuses a bit upon JLY treatment (12:45 timestamp), compared to untreated state (00:00 timestamp), PIP5K remain substantially enriched at the back-state regions of the membrane. In both timestamps, intensity profiles were normalized by dividing by the maximum intensity value at the 00:00 timestamp.

**Figure S30.**
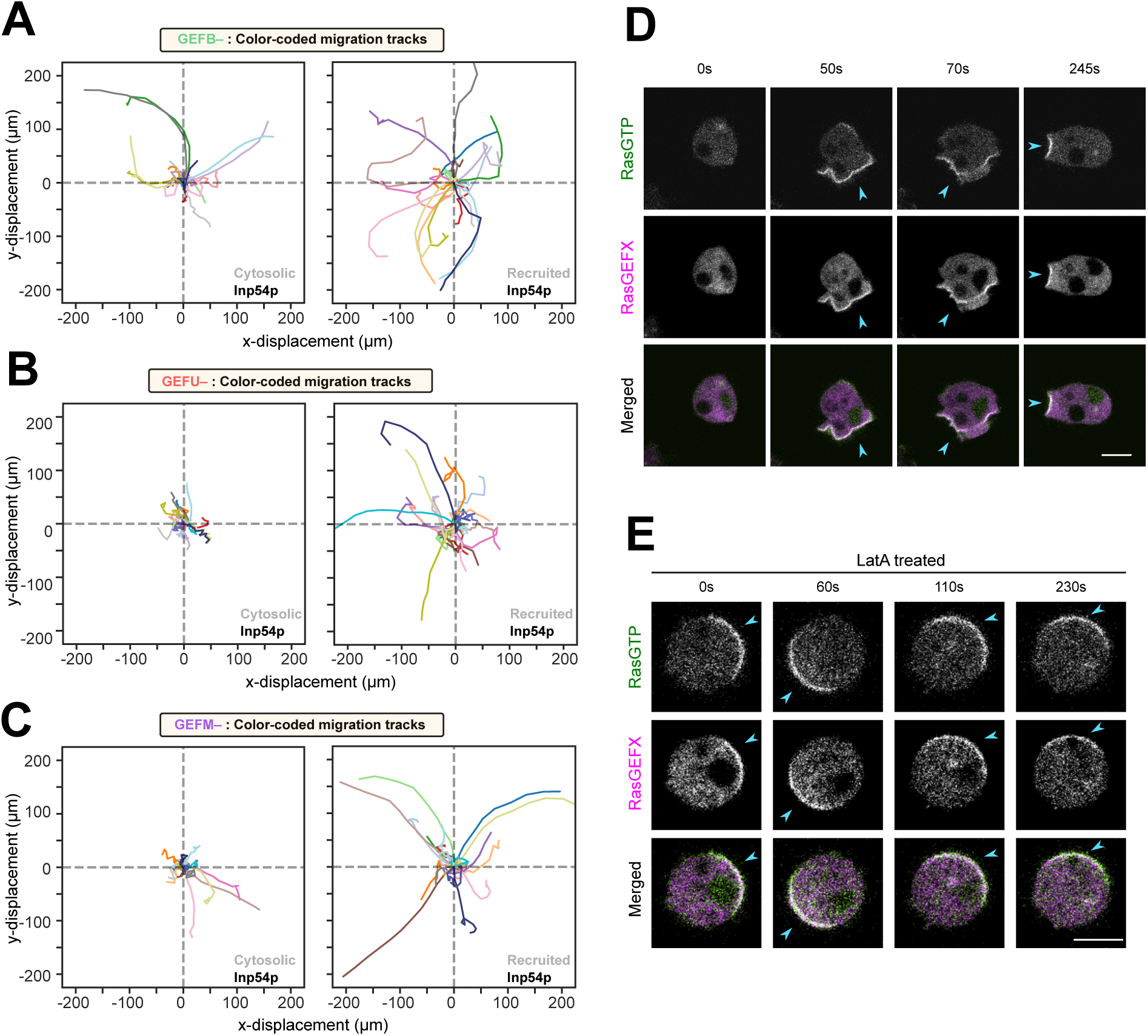
RasGEFX mediated PIP5K/PI(4,5)P2 dependent inhibition of Ras signaling and cell migration. **(A-C)** Centroid tracks of migrating *RasGEFB–* (A), *RasGEFU–* (B), and *RasGEFM–* (C) *Dictyostelium* cells, before and after rapamycin induced synthetic recruitment of Inp54p from cytosol to plasma membrane. For pairwise comparison, tracks from the same cell are plotted in same color for the before and after recruitment populations. Here, n_c_= 21 (*RasGEFB–*), 23 (*RasGEFU–*) 22 (*RasGEFM–*) cells and each cell has been tracked for t=22 min for before and after recruitment. **(D, E)** Representative live-cell time-lapse images of wild-type AX2 *Dictyostelium* cells co-expressing RBD-GFP and Halo-RasGEFX (TMR conjugated). In (E), cells were pre-treated with Latrunculin A and caffeine before starting the image acquisition. Blue arrowheads denote the front-state regions on the membrane where both RasGEFX and RasGTP are enriched.

**Figure S31.**
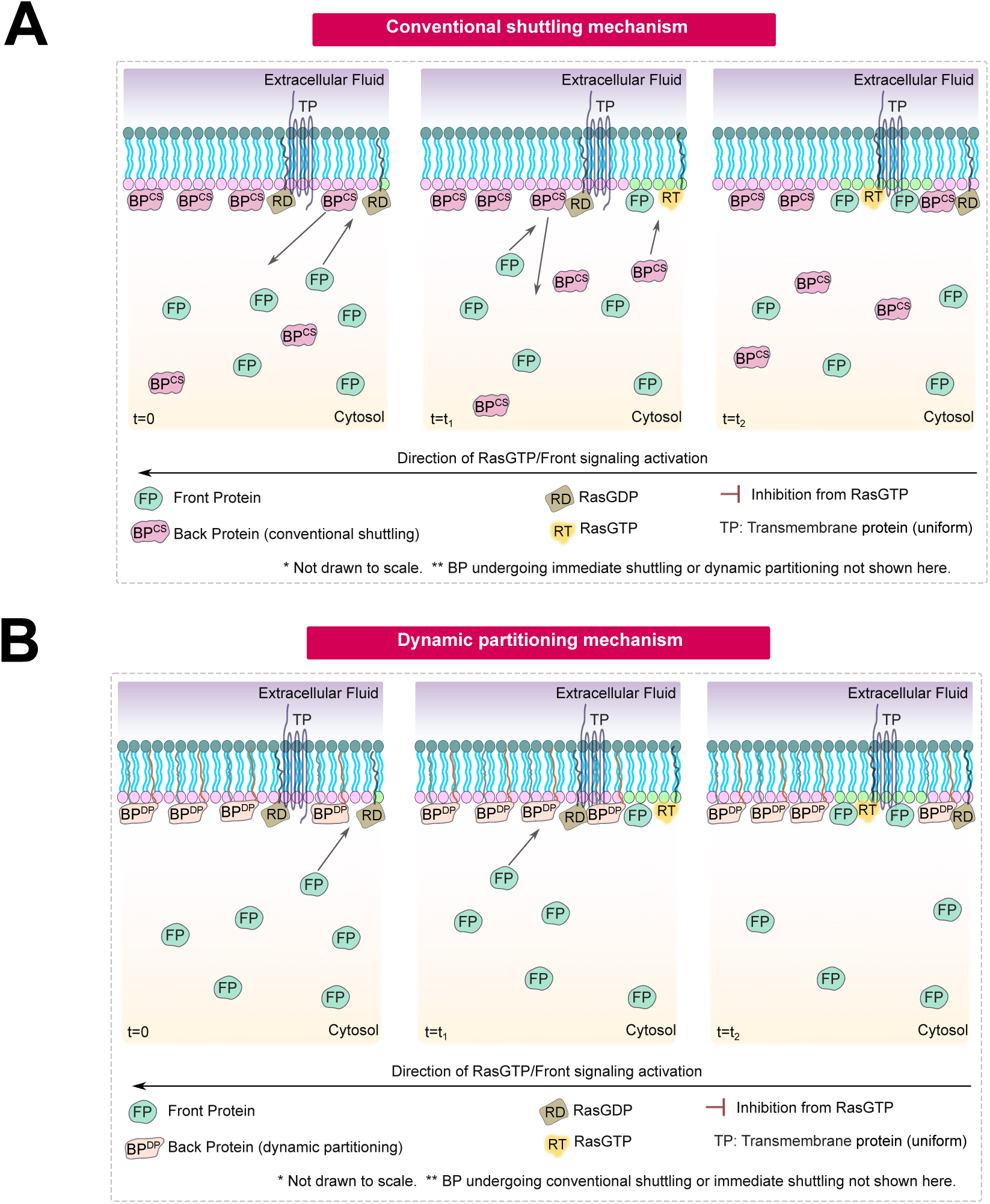
Conventional shuttling and Dynamic partitioning mechanisms for membrane protein compartmentalization. **(A-B)** Schematic showing the Conventional *shuttling* (A) and *Dynamic partitioning* mechanism (B) in plasma membrane of the cells where ventral wave is propagating. In both (A) and (B), lipid headgroups associated with the back-state and front-state are shown in mauve color and green color, respectively. At t=0, the membrane is in complete back-state and wave of signaling activation just initiated at the rightmost point, and as time passed (t_2_>t_1_>0), wave propagated from right to the left. As wave propagates and Ras switches from GDP to GTP bound state in specific membrane domains, signaling and cytoskeletal network triggers. As a result, front proteins from the cytosolic pool translocate to those membrane domains. In (A), simultaneously, membrane-bound BP^CS^ molecules move to cytosolic pool of BP^CS^. As wave propagates and some membrane domain revert to basal state, new BP^CS^ molecules from cytosolic pool get recruited there. On the other hand, in (B), simultaneously, membrane-bound BP^DP^ molecules stay on the membrane, but increase their diffusion there and quickly rearranges themselves into the other regions of the membrane where they diffuse slowly. As wave propagates, BP^DP^ molecules continue to dynamically rearrange by diffusing heterogeneously. TP molecules are shown as fixed, fiducial component. Solid arrows are shuttling between the membrane and cytosol.

**Figure S32.**
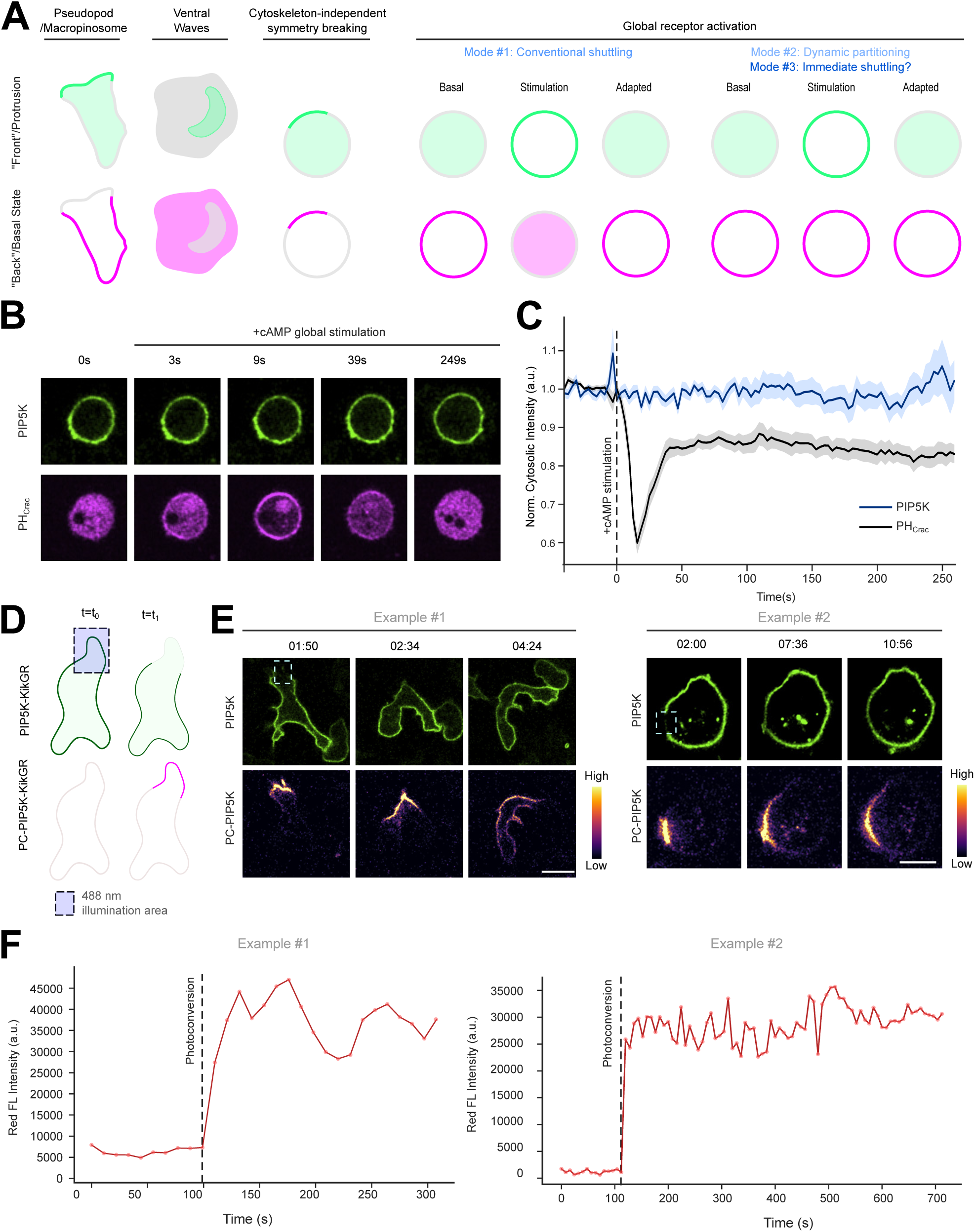
Selective photoconversion and global receptor activation assays collectively demonstrate that PIP5K does not undergo conventional shuttling. **(A)** Left three panels of the diagram illustrate the front-back complementarity during protrusion formations, ventral wave propagation, and cytoskeletal dynamics independent signaling events. In right panels of the diagram is demonstrating two distinguishable responses observed during global receptor agonist stimulation experiments, indicating the existence of at least two, and potentially three, different mechanisms that can dynamically compartmentalize proteins into the back-state regions. In contrast to *Conventional shuttling*-based polarization (Mode 1), back membrane proteins that undergo dynamic partitioning (Mode 2) or high-affinity driven immediate shuttling (Mode 3), do not dissociate from the membrane upon global stimulation, but can still compartmentalize in different physiological scenarios (also see Figure 6K for details on immediate shuttling mechanism). **(B, C)** Responses of *Dictyostelium* cells co-expressing PIP5K-GFP and PH_crac_-mCherry upon global stimulation with cAMP. Representative live-cell time-lapse images (B) and temporal profile of the normalized cytosolic intensities (C) established that front-marker PH_crac_ was recruited to the membrane during receptor activation (before going through the adaptation), whereas PIP5K remained membrane bound for the time course of the experiment. Here PIP5K-GFP was expressed under non-Dox promoter to circumvent excessive overexpression that can subdue response. The cells were pre-treated with Latrunculin A before the experiment. Vertical dashed line show the time of global cAMP stimulation. Here, n_c_=17 cells; mean ± SEM are shown. **(D)** Setup for the photoconversion experiment. Photoconvertible fluorescent protein KikGR-tagged PIP5K was expressed, a small portion of the membrane was photoconverted by selective illumination with 405 nm laser, and the photoconverted red protein was tracked over time. As cells move and make protrusions, photoconverted population should decrease over time if it generates its asymmetric distribution via the C*onventional shuttling,* but not if it does so via other mechanisms, as described above. **(E, F)** Live-cell time-laspe images (E) and quantification of red fluorescence intensity (F) in two representative cells demonstrate that, as cells migrate, photoconverted PIP5K largely stayed on the membrane over the time-course of the experiments. Light blue boxes on live-cell images show the area of selective 408 nm laser illumination. Photoconversion channel is shown in matplotlib plasma colormap so that a large intensity range can be visualized.

**Figure S33.**
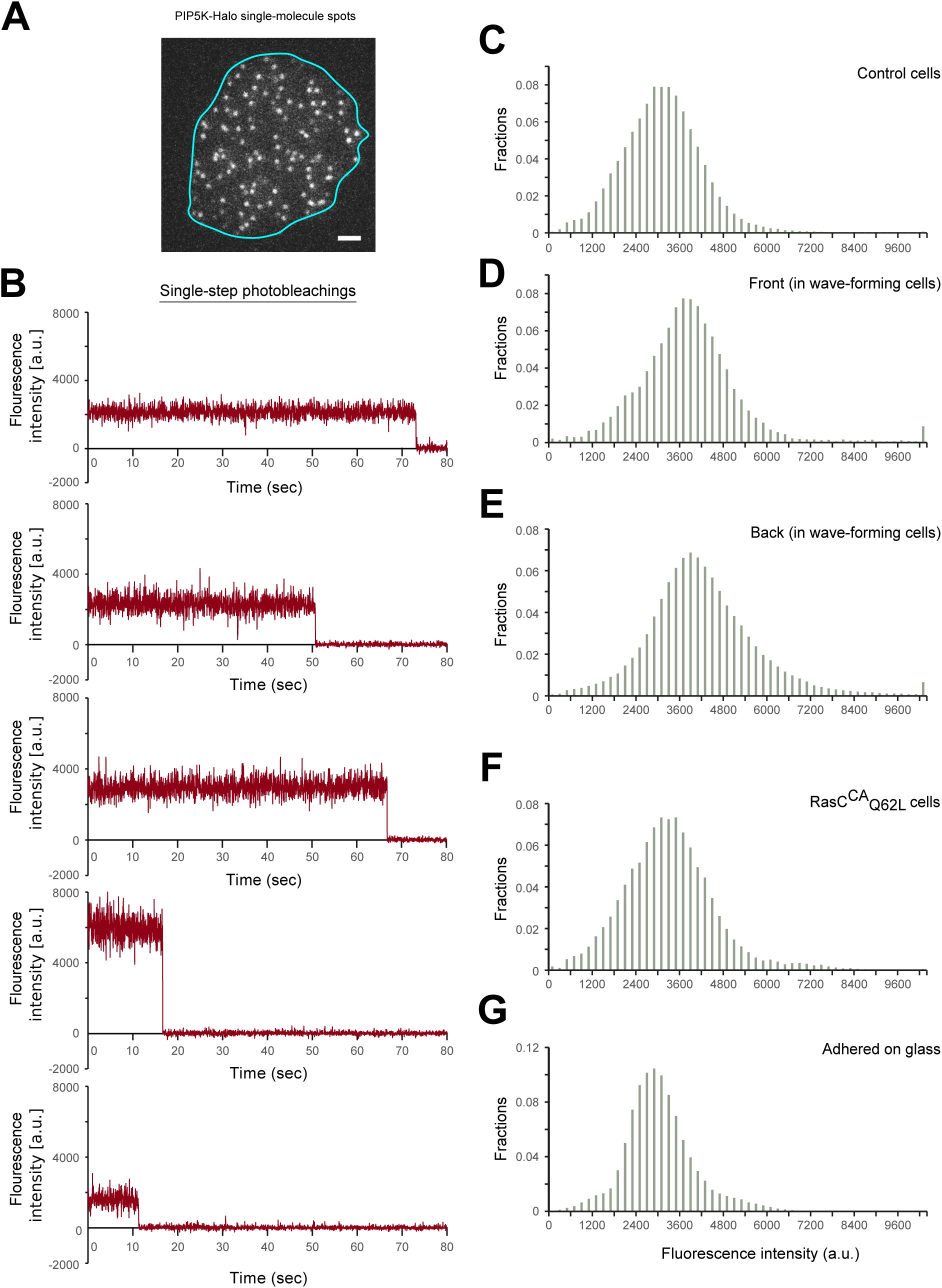
Single-molecule imaging of PIP5K under different conditions. **(A)** A representative live-cell TIRF microscopy image of a *Dictyostelium* cell displaying PIP5K-Halo-TMR molecules. Cyan line shows the outline of the cell. Scale bar indicates 2 μm. **(B)** Five representative time series plots of fluorescent intensities of TMR conjugated PIP5K-Halo exhibiting single-step photobleaching, a characteristic of single-molecule imaging. **(C-G)** Single-peaked distribution of fluorescence intensities of PIP5K-Halo-TMR was observed in control/quiescent cells (C), front-state in wave-forming cells (D), back-state in wave-forming cells (E), cells with constitutively active RasC (F), and Halo molecules on glass substrate (G).

**Figure S34.**
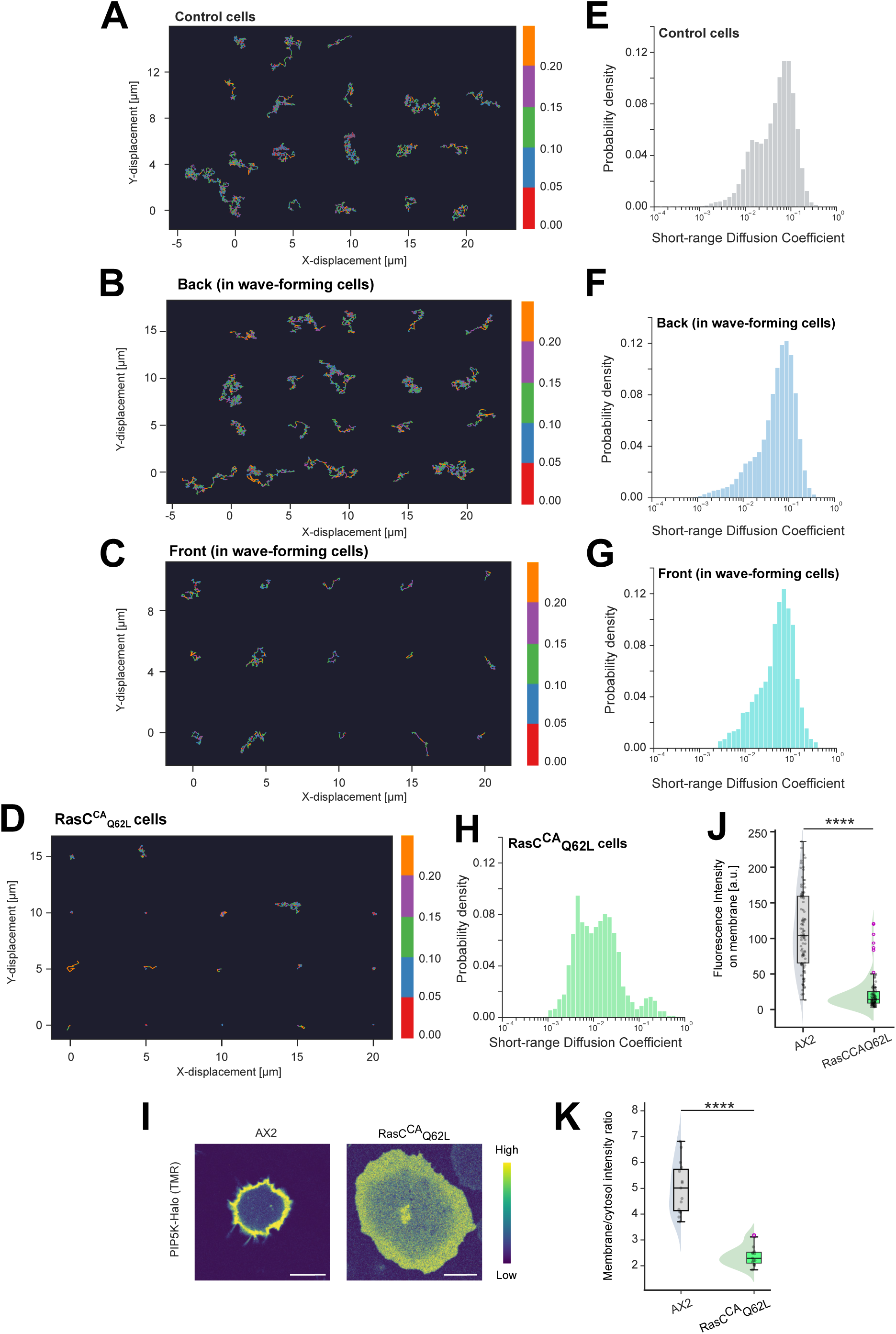
Single-molecule diffusion profiles of PIP5K in membrane domains with different levels of Ras signaling activation. **(A-D)** Representative color-coded trajectories of the PIP5K single-molecules undergoing lateral diffusion on the plane of plasma membrane inside control/quiescent cells (A), front-state in wave-forming cells (B), back-state in wave-forming cells (C), and cells with constitutively active RasC (D). Right side color bar represents the amount of displacement between two consecutive frames (in μm unit). **(E-H)** Histograms of the Short-Range Diffusion (SRD) coefficients of PIP5K molecules associated with control/quiescent cells (E), front-state in wave-forming cells (F), back-state in wave-forming cells (G), and cells with constitutively active RasC (H). SRD values are in μm^2^/sec. Note that (A-H) collectively demonstrate that the compartmentalization of PIP5K molecules inside back-state regions of the membrane do not stem from their ability to diffuse heterogeneously on the membrane. **(I)** Fluorescence intensity of PIP5K-Halo-TMR on plasma membrane in *Dictyostelium* cells with wild-type AX2 background and with constitutively active RasC, demonstrating that PIP5K significantly falls off from the plasma membrane to cytosol in cells where RasC is constitutively active. **(J)** Fluorescence intensity of PIP5K-Halo-TMR on plasma membrane in *Dictyostelium* cells with wild-type AX2 background and with constitutively active RasC. The cells adhered to coverslips were treated with Dox (50 ug/mL) for 8 hours, TMR-conjugated HaloTag ligand (5 uM) for 2 hours, PFA (4%) for 30 min and Triton X-100 (0.1%) for 30 min. Here, n_c_=100 cells for each population; p values are the Mann-Whitney U test. **(K)** Membrane to cytosol fluorescence intensity ratio of PIP5K-Halo-TMR in *Dictyostelium* cells with wild-type AX2 background and with constitutively active RasC. The z-stacks were obtained and n=4 z-sections were averaged for each cell to generate each data point. Here, n_c_=15 cells for each population; p values are the Mann-Whitney U test.

**Figure S35.**
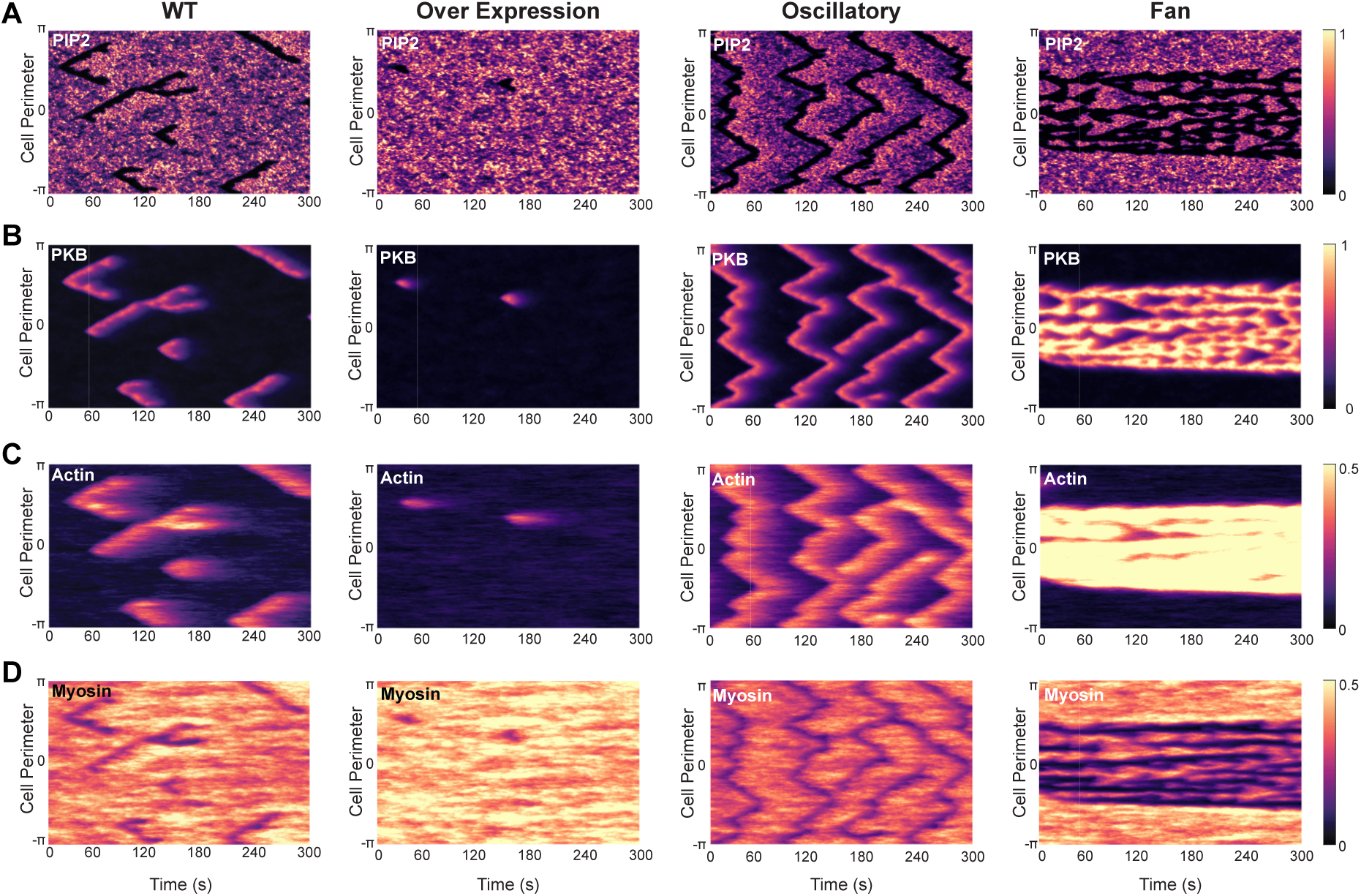
Spatial-temporal dynamics of signaling and cytoskeletal components across different phenotypes. **(A–D)** Kymographs showing molecular dynamics over a 300 s simulation: **(A)** PIP2, **(B)** PKB, **(C)** Actin, and **(D)** Myosin. Each column represents a different condition: wild type (1st), PIP5K overexpression (2nd), *piki-* oscillatory phenotype (3rd), and *fan* phenotype (4th). PKB and Actin are enriched at the front of the cell, while PIP2 and Myosin are depleted from the front and accumulate at the rear, reflecting coordinated front-rear polarization across conditions.

**Figure S36.**
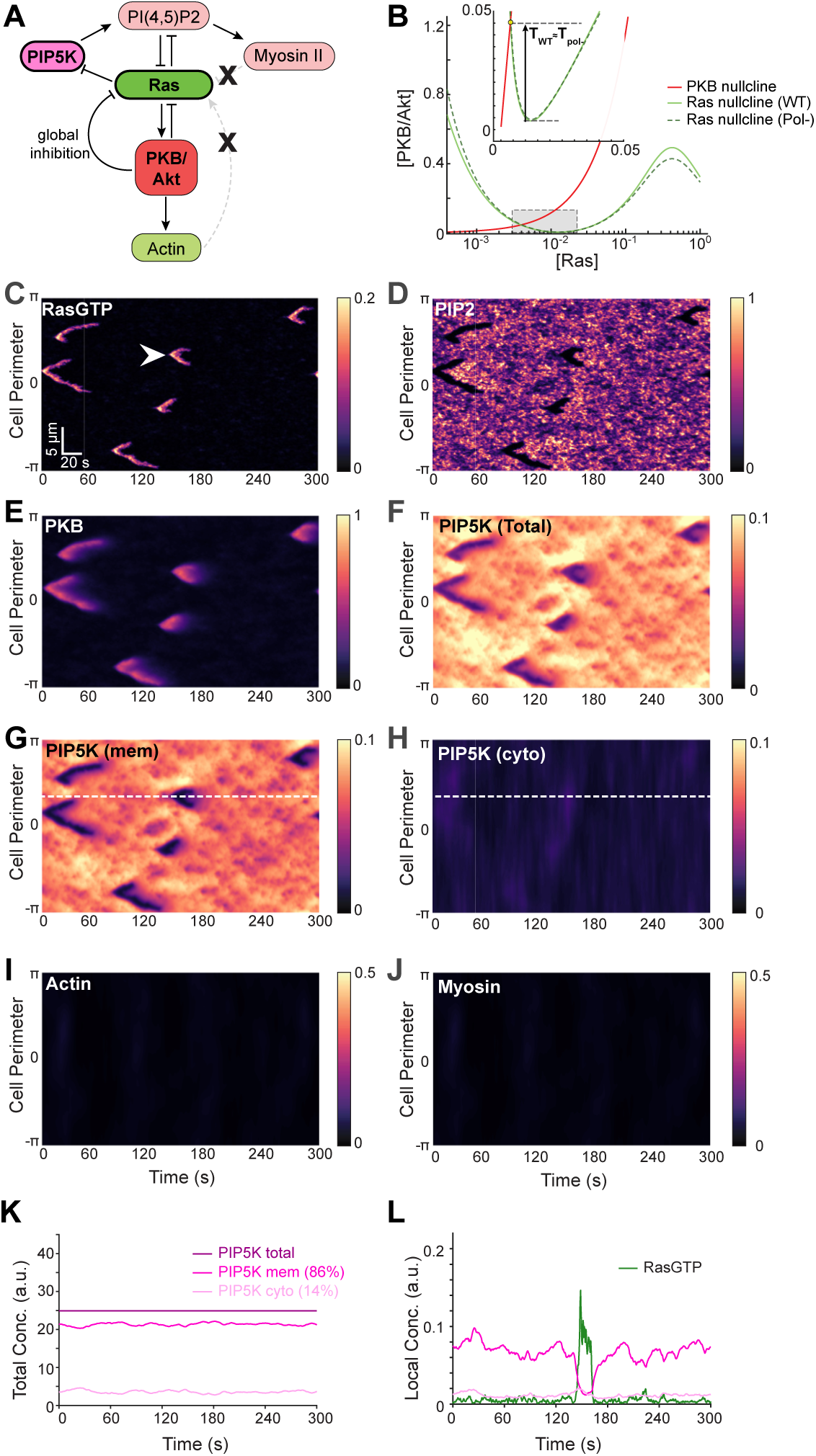
PIP5K–Ras interaction is sufficient to drive excitability in the absence of cytoskeletal feedback. **(A)** Schematic illustrating the key signaling interactions included in the model. To mimic latrunculin treatment, actin and (acto)myosin production were blocked, effectively removing their feedback to Ras signaling. This is represented by cross symbols on the corresponding feedback arrows. **(B)** Phase-portrait showing nullclines for Ras (shades of green) and PKB/Akt (red) for wild type vs latrunculin treated (pol-). Yellow marker denotes equilibrium points; the inset shows the firing threshold (𝑇_∗_), indicated by arrow, which is approximately the same between wildtype and pol-. **(C–J)** Kymographs of molecular dynamics over a 300 s simulation: **(C)** Ras, **(D)** PIP2, **(E)** PKB, **(F)** total PIP5K, **(G)** membrane-associated PIP5K; **(H)** cytosolic PIP5K, **(I)** Actin, and **(J)** Myosin. **(K)** Time series of total PIP5K (dark purple), membrane-associated (purple), and cytosolic (light purple) fractions, summed across the cell perimeter. Total PIP5K remains conserved over time (same as wild type from Fig. 7). **(L)** Local concentration profiles of Ras (green), membrane-associated (purple), and cytosolic (light purple) PIP5K at the spatial location marked by the arrow in **(C)** and dashed lines in **(G,H)**. These dynamics highlight a causal relationship: Ras activation correlates with decreased membrane association and a transient increase in cytosolic PIP5K.

## Notes

### Competing Interest Statement

The authors have declared no competing interest.

### Summary of Updates

All Figures have been updated with new data.

